# Dynamic architecture and regulatory implications of the miRNA network underlying the response to stress in melon

**DOI:** 10.1101/745653

**Authors:** Alejandro Sanz-Carbonell, Maria Carmen Marques, German Martinez, Gustavo Gomez

## Abstract

miRNAs are small RNAs that regulate mRNAs at both transcriptional and posttranscriptional level. In plants, miRNAs are involved in the regulation of different processes including development and stress-response. Elucidating how stress-responsive miRNAs are regulated is key to understand the global response to stress but also to develop efficient biotechnological tools that could help to cope with stress. Here, we describe a computational approach based on sRNA sequencing, transcript quantification and degradome data to analyze the accumulation, function and structural organization of melon miRNAs reactivated under seven biotic and abiotic stress conditions at two and four days post-treatment. Our pipeline allowed us to identify fourteen stress-responsive miRNAs (including evolutionary conserved such as miR156, miR166, miR172, miR319, miR398, miR399, miR894 and miR408) at both analyzed times. According to our analysis miRNAs were categorized in three groups showing a broad-, intermediate- or narrow- response range. miRNAs reactive to a broad range of environmental cues appear as central components in the stress-response network. The strictly coordinated response of miR398 and miR408 (broad response-range) to the seven stress treatments during the period analyzed here reinforces this notion. Although both, the amplitude and diversity of the miRNA-related response to stress changes during the exposition time, the architecture of the miRNA-network is conserved. This organization of miRNA response to stress is also conserved in rice and soybean supporting the conservation of miRNA-network organization in other crops. Overall, our work sheds light into how miRNA networks in plants organize and function during stress.

## INTRODUCTION

Because of their sessile nature, plants interact constantly with a wide array of adverse environmental cues that limit their development and productivity. These interactions with the environment can result in stress if they disturb the plant-cell homeostasis. Indeed, stress induced by both biotic and abiotic factors is one of the primary causes for losses in crops worldwide (Sunkar et al., 2007; Calanca et al., 2017). Consequently, understanding the mechanisms of stress regulation is needed to increase crop production and meet the growing challenges stemming from rapid population growth and climatic change (Brown and Funk, 2008; Takeda and Matsuoka, 2008; Fedoroff, 2010).

Plants reduce the impact of stress through a complex reprogramming of their transcriptome. These transcriptional alterations should be integrated into multi-layered regulatory networks that, collectively, modulate the interplay between the plant and stress (Dangi et al., 2018). Several studies have uncovered complex regulatory processes that coordinate plant adaptation to changing environmental conditions (Shriram et al., 2016; Haak et al., 2017; Song et al., 2019). However, certain mechanistic and structural aspects of the stress-regulatory networks remain to be fully deciphered (Zhang, 2015; Shriram et al., 2016). The elucidation of such networks is pivotal for understanding the complete molecular mechanisms that allow plants to respond and eventually adapt to the environment (Carrera et al., 2009). Using systems biology approaches may help to identify stress-responsive factors and understand their molecular interactions in detail (Dangi et al., 2018).

MicroRNAs (miRNAs) play a versatile role as translational and posttranscriptional regulators of gene expression and regulate essential metabolic processes that control growth and development (Chen, 2009; Voinnet, 2009; Sun, 2012; Xu et al., 2018). Because of this central role in the regulation of development, it was proposed that miRNAs might be good targets for the development of tools to improve crop productivity (Yao et al., 2015; Tang and Chu, 2017; Xu et al., 2018). In plants, miRNA-encoding genes are transcribed as primary transcripts harboring a fold back structure that is processed by DICER-LIKE 1 (DCL1). The outcome of this processing activity is a duplex (normally of 21 or 22 nt in length), which is later 2’-O-methylated by HEN1 and loaded into an AGO complex (Bartel, 2004; Voinnet, 2009; Reis et al., 2015). miRNAs direct target cleavage, and translational repression by means of sequences complementarity, but also other alternative functions such as mediating DNA methylation and pre-mRNA splicing (Voinnet, 2009; Reis et al., 2015; Song et al., 2019). Connected to their role in the regulation of development, miRNAs are modulators of the response to both biotic (e.g., bacterial, fungal, and viral pathogenesis; (Chaloner et al., 2016; Zhang et al., 2016; Liu et al., 2017; Brant and Budak, 2018) and abiotic stress conditions (e.g., drought, salinity, nutrient deprivation, cold, or high temperature (Shriram et al., 2016; Zhu et al., 2016; Baek et al., 2017; Bustamante *et al*., 2018). In addition, recent reports have provided evidence supporting that miRNA biogenesis (Bustamante *et al*., 2018; Manavella *et al*., 2019), miRNA turnover and miRNA-RISC assembly (Song *et al*., 2019) are also susceptible to be controlled by external stimulus. Although stress-responsive miRNAs have been identified in other species like rice and tomato, much more work is needed to decipher the conservation and structure of stress-responsive miRNA-mediated networks in relevant crops (Reis et al., 2015; Tang and Chu, 2017; Shanker et al., 2014). Indeed, most of our knowledge of the relationship between miRNA-mediated responses to stress comes from studies in the model plant *Arabidopsis thaliana*. Melon (*Cucumis melo*) is a crop of great economic importance that is extensively cultivated in semi-arid zones of diverse world regions (Wei et al., 2013). Its economic importance together with the availability of its genome sequence makes it a good target to study miRNA regulatory networks in an agriculturally important crop. Recently, melon has been shown to contain a dynamic stress-responsive miRNA population (Herranz *et al*., 2015; Sattar et al., 2012; Sanz-Carbonell et al., 2019). In this specie, miRNAs can be clustered according to their general range of response to stresses in three different groups: i) broad-, ii) middle- and, iii) narrow response-range miRNAs (Sanz-Carbonell et al., 2019). However, expression of stress-related miRNAs is expected to be dynamic and influenced by diverse factors such as the time of exposition to the adverse condition. Consequently, the predicted clustering of miRNAs may be a static picture not representative of the entire period of exposition to stress.

Here, we analyze the accumulation, functionality and structural organization of the stress-responsive miRNAs in melon plants exposed to diverse biotic and abiotic factors during different time-exposition ranges. For this end, we use sRNA sequencing, transcript quantification, degradome assays, and computational approaches. Our results indicate that although both the amplitude and diversity of the response to stress mediated by miRNAs varies during the plant exposition to the adverse environmental conditions the general architecture of the miRNA-mediated network of response to stress is conserved during the analyzed period.

## RESULTS

### Identification of stress-responsive melon miRNAs

To uncover the dynamic interplay between miRNAs and stress we constructed fifty-one sRNA libraries from melon plants exposed to seven abiotic (cold, drought, salinity and short day) and biotic (Hop stunt viroid, *Agrobacterium tumefaciens*, *Monosporascus cannonballus*) stress conditions at zero (T0), two (T1) and four (T2) days after exposure stress conditions (Table S1). Non-treated plants were used as controls. The analysis of sRNA sequences is described in the File S1. Our sRNA analysis identified a total of 17,882 (T1) and 16,869 (T2) unique reads differentially expressed during at least one of the seven stress conditions (Figure 1a). In both analyzed times cold treatment induced the most drastic alteration in sRNA accumulation (7,138 and 8,298 differential sequences in T1 and T2, respectively), whereas drought-exposed plants showed the lowest change in sRNA accumulation (Figure S3). By homology with miRNAs present in the miRNA repository miRBase (Kozomara and Griffiths-Jones, 2014), we found 17 known miRNA families in both T1 and T2 samples (Figure 1b and Table S3). Fourteen of the stress responsive miRNA families were common for both analyzed times (T1 and T2). The differential expression of these miRNA families was validated by qRT-PCR (Figure S4). Although, most of the stress-responsive miRNAs identified at T1 and T2 were coincident with miRNAs responsive to long-term treatment (Sanz-Carbonell et al., 2019), three of the miRNAs differentially expressed (miR399 and miR894 in both T1 and T2 and miR7130 in T2) were specific for early stress response.

**Figure 1:**
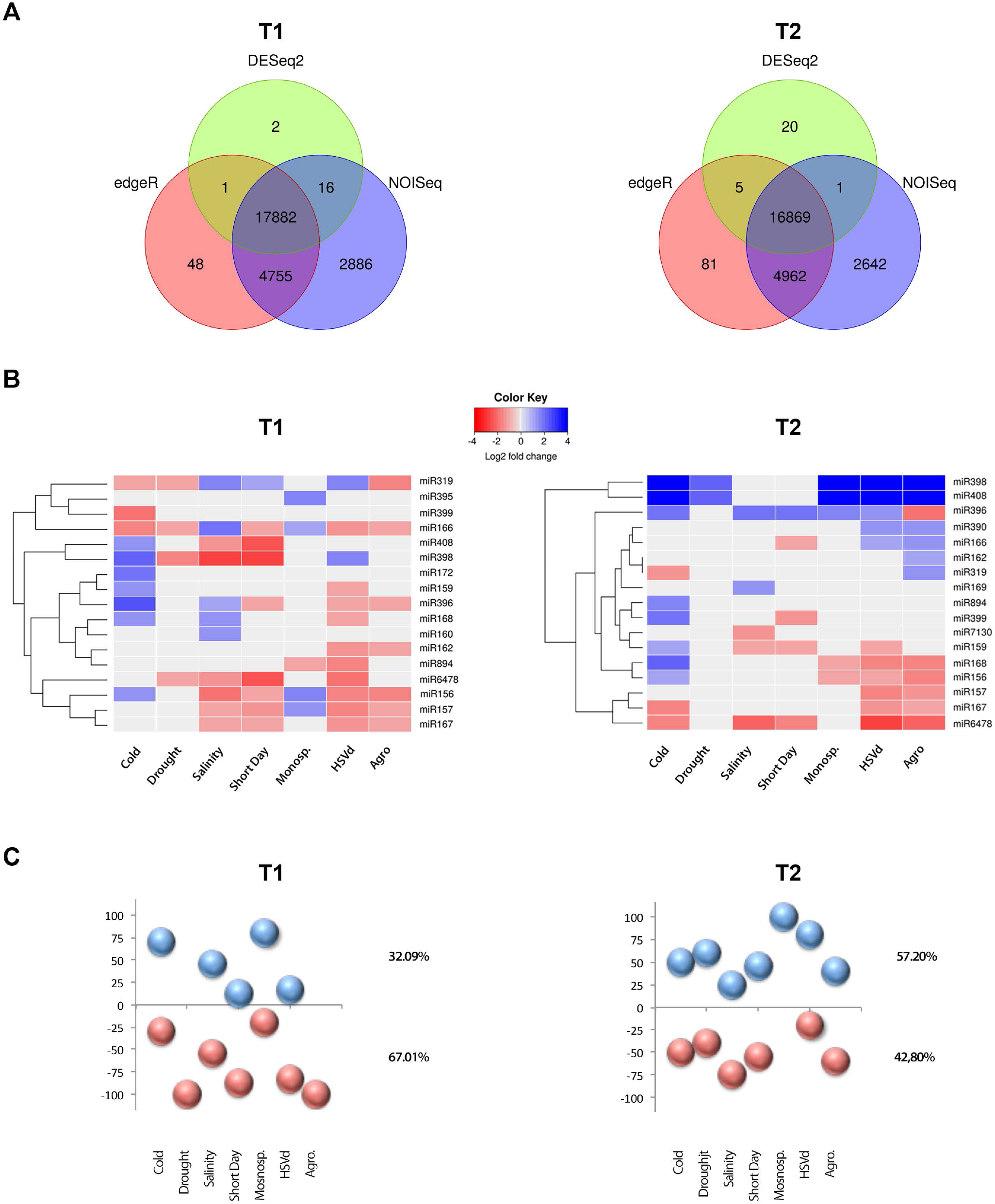
Analysis of stress-responsive miRNAs. **A)** Venn diagram comparing the number of the differential sRNAs -estimated by DESeq2 (green), edgeR (red) and NOISeq (blue)-expressed in melon in response to cold, drought, salinity, short day, *Monosporascus* (Mon), HSVd and *Agrobacterium* (Agro) treatments at both analyzed times. Only the sRNAs predicted as differential by all three analysis methods were considered as true stress-responsive miRNAs. **B)** Heat map of 17 miRNAs families differentially expressed in melon plants in response to stress at T1 and T2. The differential expression values represented correspond to the median of the Log_2_FC values obtained by DESeq2 analysis in each miRNA family, except for miR396 in HSVd (T2) and miR166 in cold and short day (T1) were we considered the expression value of the most highly represented miRNA-related sequences. **C)** Graphic representation of the relative accumulation (in percentage) of melon miRNAs up-(blue balls) and down-regulated (red balls) in plants exposed to different treatments.

The general response at T1 consisted in a general (67%) down-regulation of miRNA accumulation (Figure 1c). This phenomenon was more evident in drought and *Agrobacterium* treatments with 100% of the differential miRNAs being down regulated. However, in T2 approximately 57% of the stress-responsive miRNAs were up-regulated (Figure 1c). Plants exposed to *Monosporascus* and HSVd showed an accumulation of up-regulated stress-responsive miRNAs clearly above average (100% and 80% respectively, Figure 1c). In general, members in each miRNA family respond in a coordinated manner to each analyzed stress condition (Figure S5). The only exception to this rule was the expression of miR396-related sequences in HSVd-infected plants (T2) and miR166 in cold and short day treatments (T1). However, sequences with antagonist behaviour were close to the limits established for differential expression (LFC near to −1 or 1) and poorly recovered from analyzed samples (Table S4), for those families we only considered the expression value of the most highly represented miRNA family-related sequence.

### miRNA-regulated targets in response to stress in melon

The majority (16 out of 19) of the miRNA-targeted mRNAs with differential expression at both T1 and T2 agree with the previously described and validated miRNA-targeted mRNAs in melon plants exposed to longer stress period (Sanz-Carbonell et al., 2019). These targets were mainly transcription factors (TFs, 9 out of 19, including: SPL, BEH4, ARF17, ATHB14, ARF, AP2, TCP, GRF and NFY) (Table S5). The miRNA-mediated regulation of TFs under stress conditions in diverse plants species reinforce these results (Matthews *et al*., 2019; Zhou and Tang, 2019; Arshad *et al*., 2017 a and b; Li *et al*., 2017; Chen *et al*., 2015). Other transcripts regulated by stress-responsive miRNAs were functionally associated with oxidation-reduction processes (miR398, miR408 and miR6478), RNA silencing (miR168), photosynthesis-related processes (miR162), and metal metabolism (miR395) (Table S5). We further validated by 5’-RLM-RACE, the transcripts regulated by the stress responsive miRNAs specific for T1 and T2 (miR894, miR399 and miR7130, Figure S6b). Our analysis indicated that these three miRNAs regulate mRNAs involved at different levels of the stress-response homologous to Cysteine-rich/transmembrane domain A-like, Beta-glucosidase 11-like and Constitutive photomorphogenesis 9 signalosome (sub-unity 6a) proteins (Figure S6a) that regulate cell death, antioxidant accumulation and hormonal balance (Singh and Chamovitz, 2019; Baba et al., 2017; Yadeta et al., 2017).

To test the functional role of the stress-responsive miRNAs, we analyzed the correlation between miRNA levels and their target accumulation. We focused on miRNAs reactive to early stress conditions (miR156, miR166, miR172, miR319, miR398 and miR408) with strong differential expression at both analyzed times (T1 and T2). A significant negative correlation was obtained when we compared the expression values of stress-responsive miRNAs with the accumulation of their target estimated by qRT-PCR (Figure 2).

**Figure 2:**
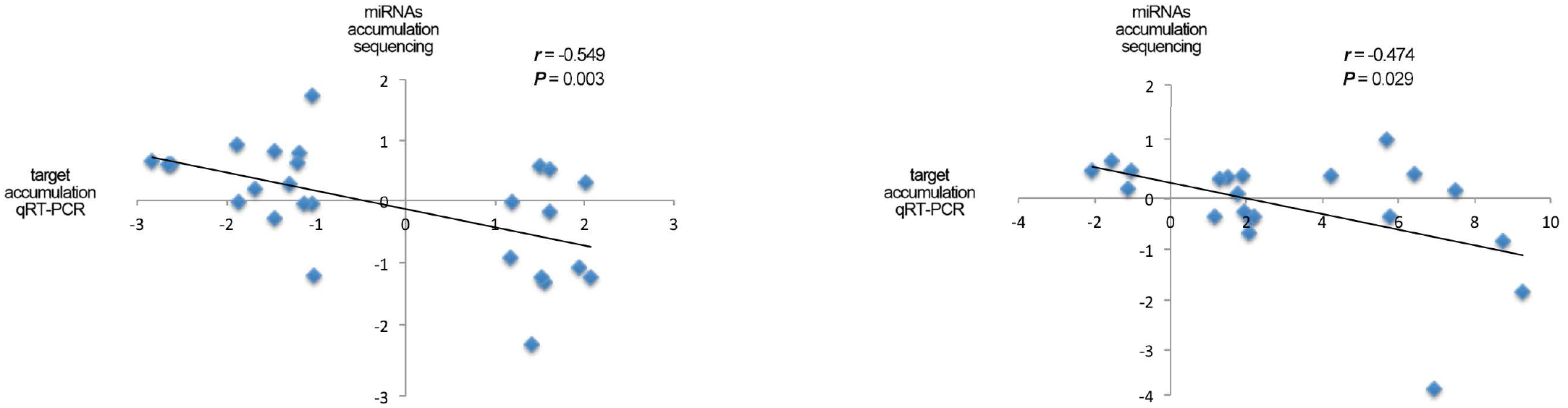
Validation of functional miRNA-targets interactions by qRT-PCR assay. Scatter plot showing the significant negative correlation (estimated by *Pearson correlation coefficient*) between the expression levels of selected stress-responsive miRNAs with differential accumulation determined by sequencing data and the accumulation of their predicted targets in the corresponding stress situations estimated by qRT-PCR (detailed information in Table S15).

### Melon miRNAs show common architecture of response-range to stress

We used a principal components analysis (PCA) to infer the organization of the miRNA-mediated response to stress in melon. Our results support that at both analyzed T1 and T2 time points stress-responsive miRNAs were organized into three different groups according to their responsiveness to stress (Figure 3). These groups include a group of miRNAs (six in T1: miR156, miR157, miR166, miR319, miR396 and miR398; and four in T2: miR396, miR398, miR408 and miR6478) reactive to a broad range of stress conditions (with modified expression in five or more stresses). A second cluster includes those miRNAs responsive to an intermediate range (3 or 4) of adverse environmental conditions (miR267, miR168, miR408 and miR6478 in T1; and miR156, miR159, miR166, miR167 and miR168 in T2). A more abundant group of miRNAs differentially expressed under a narrow number (1 or 2) of stress conditions were clustered independently (seven in T1 and eight in T2). The statistical significance of the differences between the predicted groups was estimated considering the Euclidean distances between components (accumulative proportion of variance for PC1 and PC2 of 77.02% for T1 and 74.85% for T2) (Figure 3).

**Figure 3:**
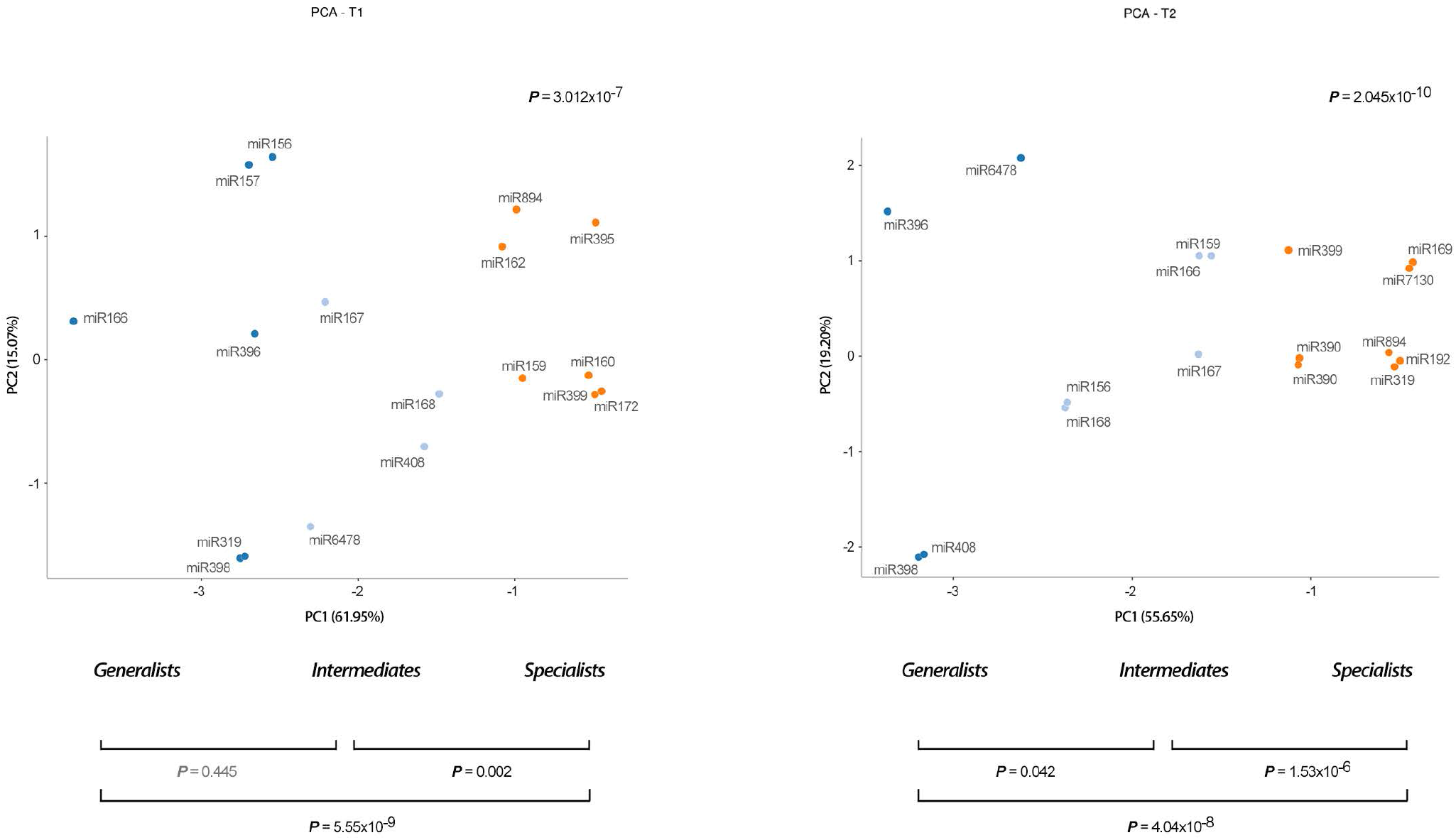
Stress-responsive miRNAs are hierarchically organized in relation to its response-range to stress. Principal component analysis of stress-responsive miRNAs detected in melon plants at both analyzed times T1 (left panel) and T2 (right panel). Values of proportion of variance for PC1 and PC2 are showed y the X and Y-axis. The statistical significance of the identified clusters was estimated by Mann-Whitney-Wilcoxon test, considering the inter- and intra-group Euclidean distances (*P*-values are showed in the graphic). The different groups of stress-responsive miRNAs detected in melon are identified by colors: *broad response-range* (deep blue), *intermediate* (light blue) *and narrow* (orange).

### The stress-responsive network mediated by miRNAs in melon

To infer the general architecture of melon miRNA network at T1 and T2, we grouped all stress-responsive miRNAs in pairs and searched for the stress conditions to which both were responsive (i.e., either up- or down-regulated). If the two miRNAs reacted to a particular stress condition were considered connected. This information (about miRNAs behaviour under adverse environmental conditions) was used to construct a stress-response miRNA network (Figure 4). On these networks, nodes represented reactive miRNAs and links indicated the relation between miRNAs with common response to at least one stress condition (Figure 4). The width of the links in the networks represented the strength of the connection between the two miRNAs connected by that link (the higher the proportion of stress that was common in the response of two miRNAs, the thicker was the edge connecting these miRNAs).

**Figure 4:**
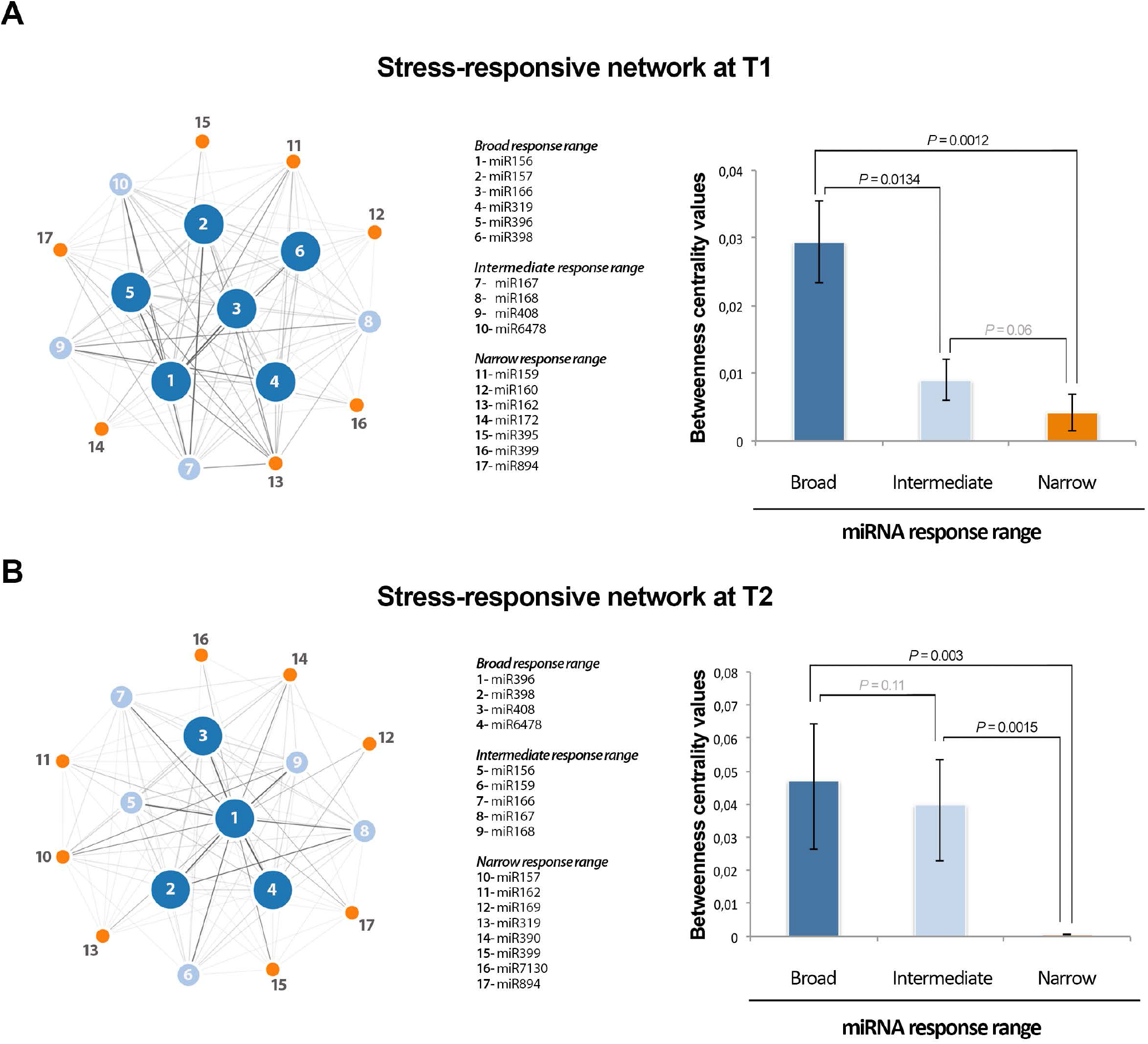
Network of stress-responsive miRNAs in melon. **Left panels)** Nodes in the network represent differentially expressed miRNAs. Colors and numbers depict the different groups of stress-responsive miRNAs detected in melon. Node size is proportional to the number of stress conditions where a particular miRNA is differentially expressed (5 or more for broad, 3 or 4 for intermediate, and 1 or 2 for narrow response range). Edges represent weighted associations between the terms based on response to common stress conditions. **Right panels)** Graphic representation of the average *betweenness* of the nodes calculated for broad, intermediate, and narrow response range miRNAs in melon. The statistical significance of the differences was estimated by Mann-Whitney-Wilcoxon test, significant *P*-values (< 0.05) are highlighted in bold, and non-significant values in gray. Error bars represent the standard error in *betweenness* values.

The obtained miRNA responsive networks revealed an architecture evidencing two main layers (Figure 4a and b, left panels). One central module that includes highly connected miRNAs (broad response range), and another peripheral layer composed by stress-related miRNAs with lower connectivity (narrow response-range). Finally, the miRNAs that exhibit an intermediate response-range were distributed in a more undefined layer dispersed between both principal network components. The statistical robustness of the inferred network architecture was estimated by analysis of the *Betweenness* values obtained for each miRNA in the network (Figure 4b, right panels).

### The amplitude of the miRNA-mediated response is increased over time

To draw a map of the dynamic evolution of the miRNA-mediated response along exposition to stress, we combined the results obtained here, with the previous data obtained from melon plants exposed to identical stress conditions during a longer time (11 days - T3) (Sanz-Carbonell et al., 2019). Box-plot analysis of all stress-responsive miRNA families provided evidence about two main characteristics of the temporal evolution of the response to stress in melon (Figure 5). First, the general shift of the miRNA expression (up- or down-regulation) is variable along the analyzed period of exposition to stress and the stress responsive miRNAs mainly down regulated in T1 (−1.04) and T3 (−1.39) and up regulated at T2 (1.51). Second, the amplitude (estimated by the variance of the differential expression values) and diversity (estimated by the number of miRNA sequences with differential expression) was clearly increased during the exposition to the stress conditions. The increase in the diversity of the global miRNA-related response in melon plants is a consequence of two different but complementary phenomena, *i*) an increase in the number of reactive miRNAs (17 in earlier phases of the stress response and 24 miRNAs in T3) and *ii*) differential expression of additional members in each miRNA family (48 family-related sequences in T1 and 96 sequences in T3) (Table S6).

**Figure 5:**
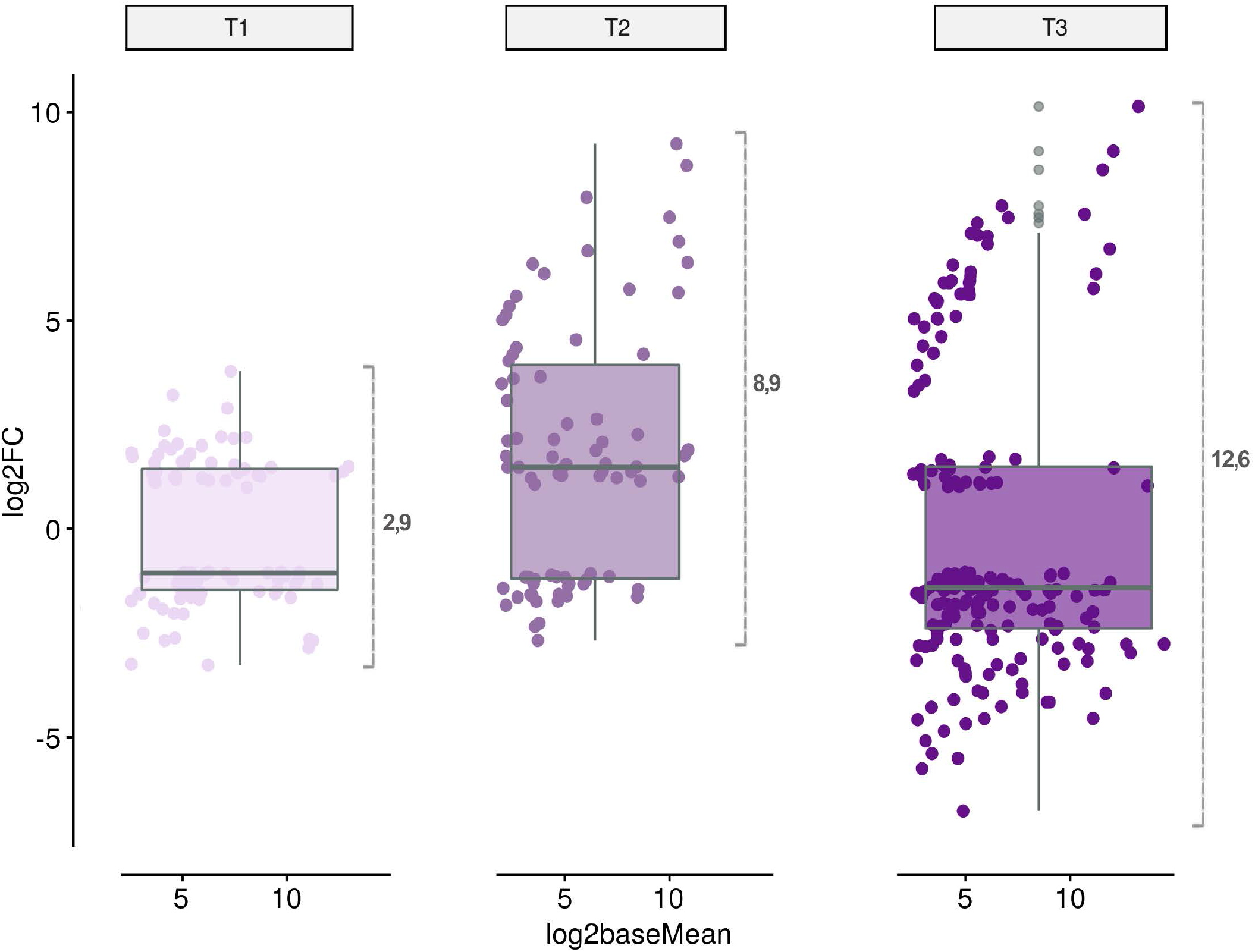
The amplitude and diversity of the miRNA-mediated stress response is increased over time. Analysis of the temporal evolution of the stress-response in melon. The dots represent each one of the miRNA-family related sequences reactive to analyzed stress conditions at two days (T1), four days (T2) and eleven days (T3) post treatment. The differential expression (LFC, in the Y axis) and accumulation (Log2 of the base mean, in the X axis) values of the stress-responsive miRNAs represented in the figure correspond to the data obtained by DESeq2 analysis. The internal box-line represents the median of the miRNA expression levels in each analyzed time. At the right of the box-plot is represented the variance of the differential expression values in each analyzed time (used as indicator of response amplitude).

### The response-range of the stress-associated miRNAs is consistent during the time exposition to stress

Clustering analysis supports that under the stress conditions tested here responsive miRNAs may be functionally organized in three differentiated groups identified as broad-, intermediate- and narrow response range. Our results reveal that this architecture is applicable to both early (Figure 3) and long-term response to adverse environmental conditions (Sanz-Carbonell et al., 2019), when the stress-responsive miRNAs are considered at each exposition time individually and under a static viewing. However, global expression of stress-related miRNAs is expected to be a dynamic phenomenon related to, besides the type of stress, additional variables as for example, the time of exposition to the adverse condition.

To obtain a more dynamic view of the general architecture of the stress response mediated by miRNAs, we used a modification of the *Sankey* diagram (www.sankeymatic.com), where we represented the identified response ranges (broad, intermediate, and narrow) in each one of the three analyzed times (T1, T2 and T3). The results, presented in Figure 6, revealed that the range of response of a particular miRNA to stress is maintained constant across the time of exposition. For example, miRNAs identified as responsive to a broad-range of stress conditions maintain this characteristic (or an intermediate response range) during the analyzed period with the exception of miR157 and miR319 (identified as narrow-range in T2). A more evident conservation of the range of response to stress through this study was observed for miRNAs showing a narrow response-range. A high proportion of them (11 out 14) strictly maintain this functional category during the entire period of exposition to the diverse stress conditions (boxed miRNAs in Figure 6). Furthermore, five out of eight miRNAs identified as stress-responsive in a particular analyzed time (miR7130 in T2 and miR164, miR165, miR394 and miR1515 in T3) were also clustered as narrow response-range miRNAs, suggesting that these reactive miRNAs have a specific functional role (considering both, the type and the duration of the adverse environmental condition) in the regulation of the response to stress in melon.

**Figure 6:**
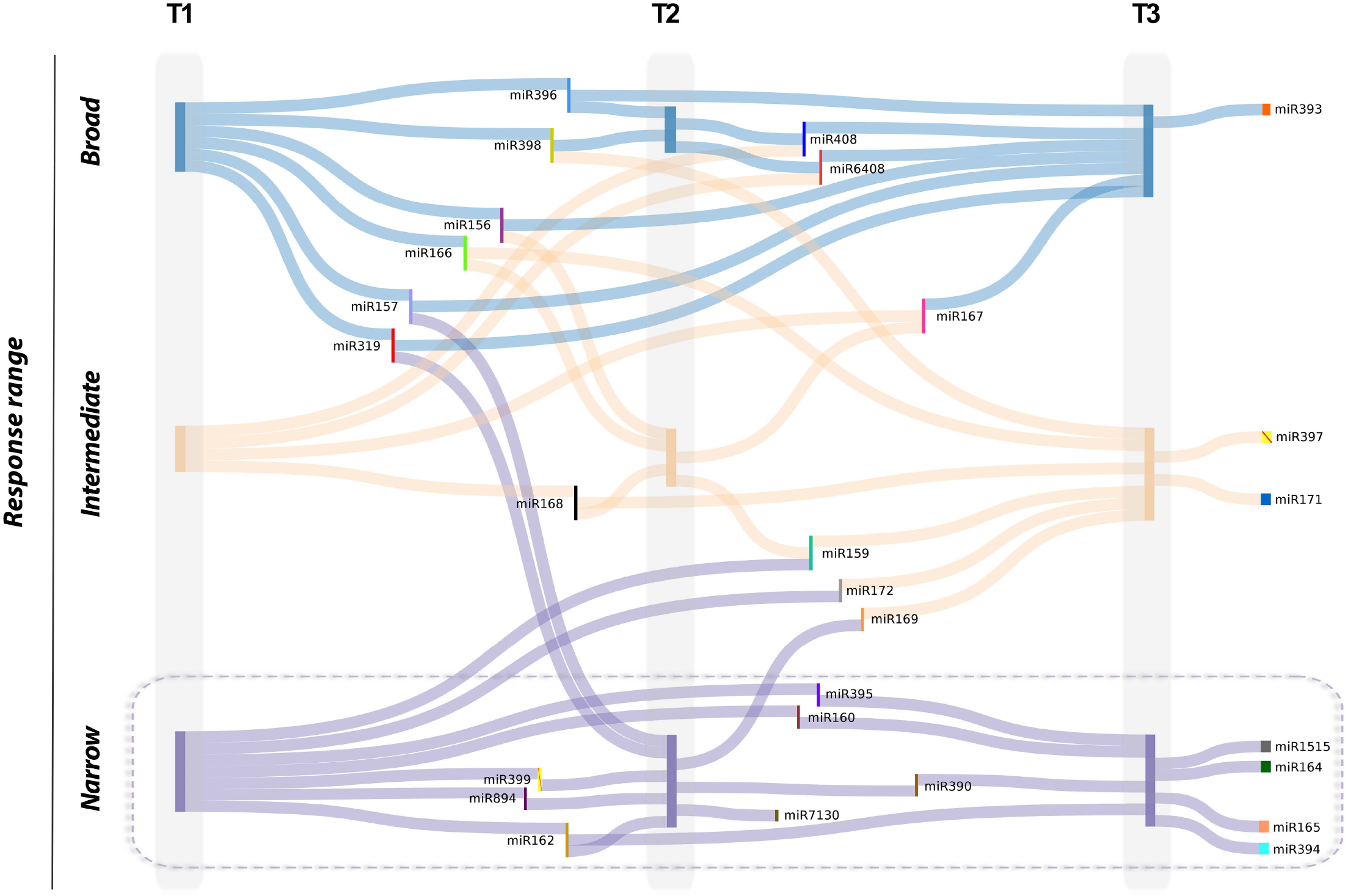
The response range of miRNAs reactive to stress is in general constant during the stress treatment. Viewing of the response-range evolution for stress-related miRNAs through the three analyzed time periods. miRNAs with broad-response range are represented by blue lines). Vertical bars represent nodes for miRNAs reactive to *broad* (blue), *intermediate* (orange) and *narrow* (magenta) range of stress conditions at each one of the analyzed times: two days (T1), four days (T2) and eleven days (T3) post treatment. Thin vertical bars in colors are used to integrate miRNA-labeling. Strict *narrow response-range* miRNAs are boxed.

Regarding the functional aspects of the miRNAs playing a more generalist or specialist role in the regulation of the stress response (defined by the biological role of their targets), it was evident that miRNAs with a narrow response-range are mainly characterized as regulators of transcripts associated to stress response and basic cell functions such as metabolism (of metals, carbohydrates, and carotenoids), reproduction, RNA silencing and photosynthesis (Table S7). Although further studies are needed in order to elucidate the roles played by miRNAs with narrow response-range in melon, the observation that miRNAs such as miR395 (Du et al., 2018), miR894 (Xie et al., 2014) or miR394 (Song et al., 2013) -for example- have been previously associated to diverse biotic and/or abiotic stress responses allow us to speculate about a potential selective stress-related regulatory activity. In contrast, miRNAs that exhibit a broad response-range seem to be master regulators of central hubs in the control of the gene expression, targeting predominantly (70% of the identified targets) TFs related with plant development (Table S7).

### miR398 and miR408 are coordinately altered in all stress conditions

To analyze the patterns in miRNA levels at different time intervals in relation to each particular stress treatment, we performed a time-course analysis of the expression of the reactive miRNAs (considering only families recovered in all analyzed times, including T0 as convergent starting point). This analysis detected 26 miRNA families whose expression profiles were significantly altered in at least one of the analyzed times in response to, at least, one stress condition. The cluster analysis identified six groups of stress-responsive miRNAs associated to drought and salinity treatments, five for cold, four for short day, monosporascus and HSVd infections and three different groups for melon plants infected with agrobacterium (Figure 7). The general expression pattern of miRNAs reactive to the various analyzed conditions was diverse, being constantly altered (up- or down-regulated) during stress in some groups or alternating increased, non-differential and/or decreased expression in others. To simplify the identification of common dynamic patterns of miRNA expression in response to stress, we used these data to perform an additional “*soft*” clustering analysis in which each data (miRNAs in this case) can belong to more than one cluster (Lu et al., 2013). Next, we searched the clusters in which every pair of miRNAs was grouped. If two miRNAs were related to a particular cluster then we considered them to be connected. This information (Table S8) was used to construct a cluster-related miRNA network in which nodes represented miRNAs and edges indicated the links between miRNAs with common estimated clustering in response to stress (Figure 8a). Notably, miR398 and miR408 (two miRNAs predominantly characterized, as *broad response range* were the most strongly related miRNAs, sharing cluster under all analyzed stress-conditions. In addition to a highly coordinated expression in response to stress, both miRNAs also showed a specific response pattern, sharing clustering only with miR157 and miR167 (under salinity conditions) and with miR6478 (short day). According to their expression patterns the miR398-miR408 response can be ordered in four different stress-related groups (Figure 8b), supporting that, despite being highly coordinated, the miR398-miR408 response in melon may be determined by the stress conditions.

**Figure 7:**
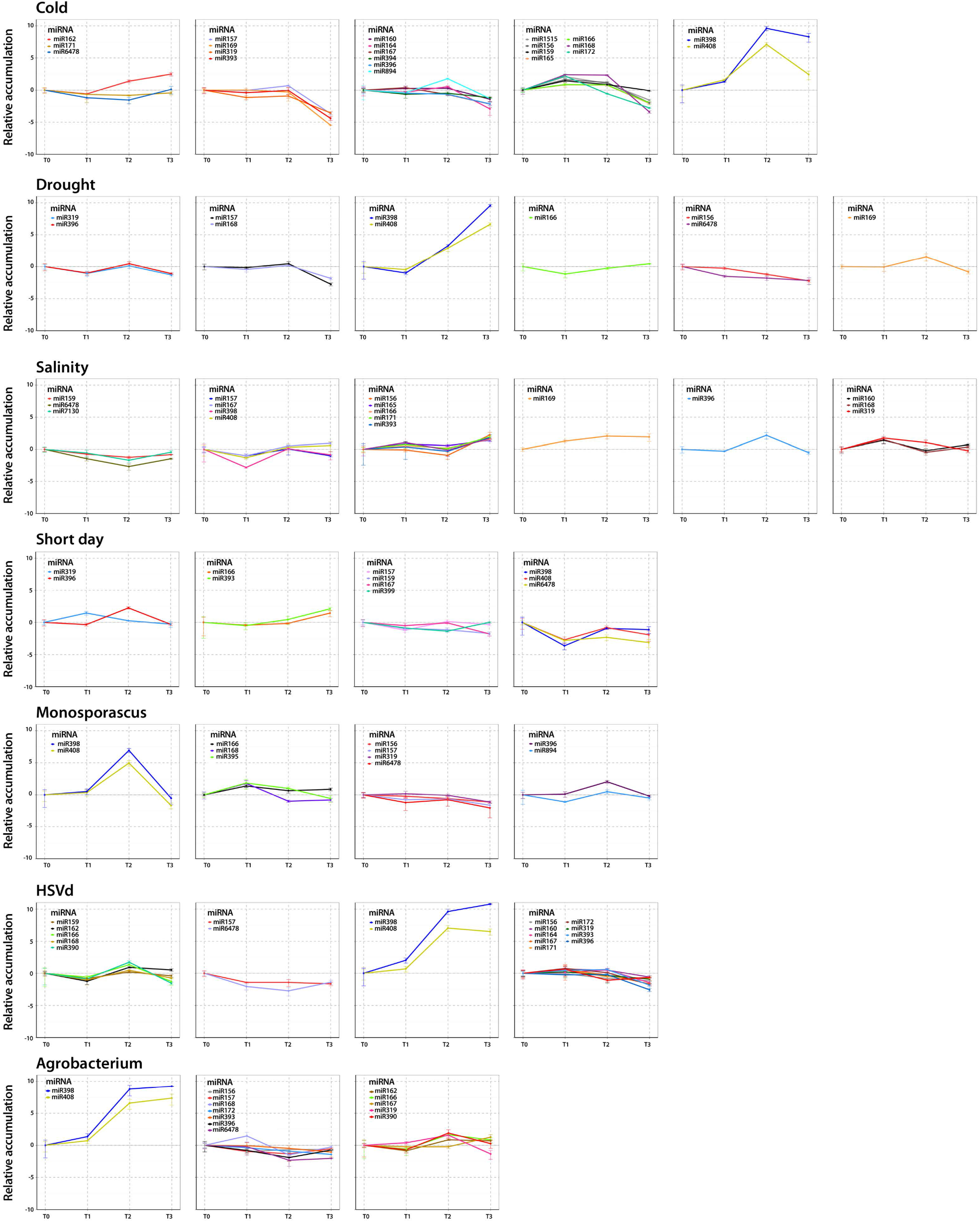
Comparative analysis of the expression patterns of responsive miRNAs in each analyzed stress condition. Clustering analysis of time-course expression profiling of responsive miRNAs in each one of the biotic and abiotic stress condition evaluated in this work. Error bars indicate standard error between replicates.

**Figure 8:**
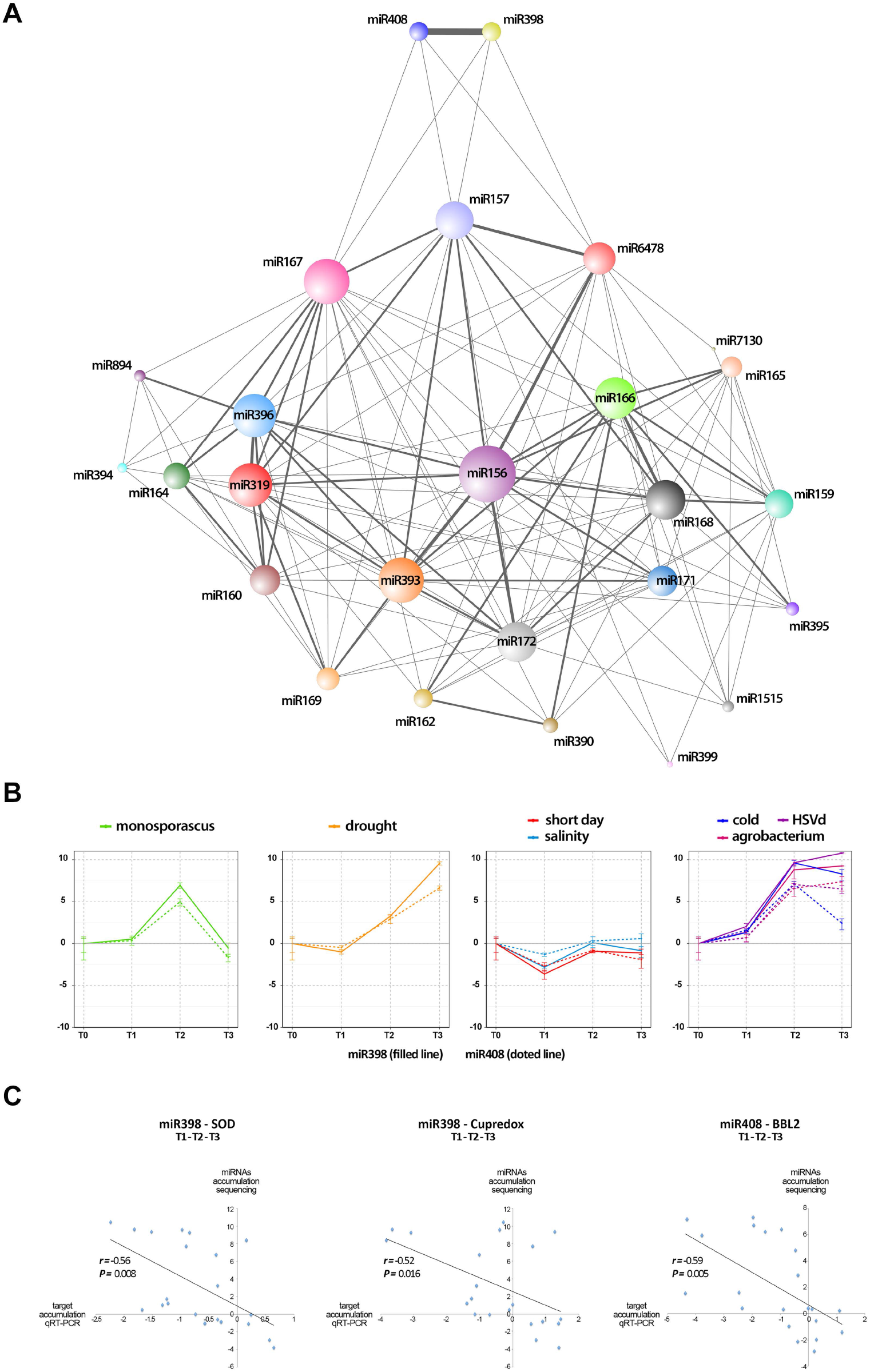
Network of expression profiling of stress-responsive miRNAs. A) Nodes in the network represent differentially expressed miRNAs. Edges represent weighted associations between the terms based on response to common expression profiles in response to analyzed stress conditions. Edges thickness is proportional to the number of shared expression profiles. B) Time-course expression profiling of miR398 and miR408 clustered according to their expression profiles in response to the seven stress conditions analyzed here. C) Scatter plot showing the significant negative correlation (estimated by *Pearson correlation coefficient*) between the expression levels of Cu-miRNAs reactive to stress conditions (miR398 and miR408) according sequencing data and the accumulation of its predicted targets (estimated by qRT-PCR) during the period of exposition to the biotic and abiotic stress conditions analyzed here (detailed information in Table S16).

### Fine modulation of Cu-related proteins is a general phenomenon associated to the stress-response in melon

Although miR398 and miR408 are commonly known as Copper (Cu) miRNAs involved in the regulation of Cu metabolism (Burkhead et al., 2009), a large body of evidence links the differential expression of these Cu-miRNAs to a general stress response in plants (Pilon, 2017). Melon miR398 is predicted to mediate the cleavage of (Cu) Superoxide dismutase (SOD) and Cupredoxin transcripts, while miR408 regulates the expression of a putative Basic blue protein-like transcript (BBL) (Sanz-Carbonell et al., 2019). In order to establish a more precise functional correlation between miR398 and miR408 levels and the response to stress in melon, we analyzed the temporal expression of the three targets across the time-exposition to the diverse stress conditions described here. Quantitative real time-PCR (qRT-PCR) assays revealed a significant negative correlation between the global expression of miR398 and miR408 and the accumulation of SOD, Cupredoxin and BBL transcripts (Figure 8c) and (Figure S7).

### The architecture of the miRNA-mediated response to stress is conserved in other crops

To explore the potential conservation of stress-responsive miRNA networks in other plant species we analyzed publicly available datasets. Concretely, we performed a differential expression analysis, using sRNA-data obtained from rice (*Oryza sativa*) and soybean (*Glycine max*) plants exposed to diverse biotic and abiotic stress conditions (Table S9). According to the analysis parameters described in File S2 we identified 85 and 72 stress-responsive miRNA families in rice and soybean respectively (Figure S8). These families are also functionally organized into the three different groups identified for stress-responsive miRNAs in melon (Figure S9).

In both species, our results revealed a network architecture with a central module of broad response-range miRNAs, a peripheral layer composed by miRNAs with lower connectivity and narrow response-range and an undefined layer dispersed between both principal network components containing miRNAs with an intermediate response-range (Figure S10). The presence of a comparable stress-responsive miRNA network in melon, rice and soybean, supported the notion that the global architecture of the stress response mediated by miRNAs might be conserved, at least for the crops analyzed here. We further analyzed the conservation of the miRNA families included in each group. To address this issue we grouped stress-responsive miRNAs identified in melon, rice and soybean according to their response range (broad, intermediate and narrow) and considered as connected the miRNAs families sharing response range. According to this we constructed a response-range network in which nodes represented the analyzed crops and links indicate the connection between miRNA families. The results showed in Figure 9 revealed that the functional categorization of stress-responsive miRNA families is conserved in the analyzed crops. A great proportion (14 out of 20) of broad response-range miRNA families maintain this behavior (or an intermediate response-range) in the analyzed crops. Regarding miRNAs that exhibit a more specific stress-related activity (narrow response-range), the percentage of them that strictly maintain this functional category in any of the three crops analyzed is higher than 90% (Figure 9). To obtain a more quantitative vision about this phenomenon we analyze the connectivity level for miRNAs grouped in the three functional categories in melon, rice and soybean (Table S10). The data obtained from this analysis (detailed in File S1), evidenced that miRNAs reactive to a broad range of stress conditions, showed the higher connectivity level (CV=1.3). In contrast narrow response range miRNAs had a lower connectivity (CV=0.036) and are predominantly specie-specific. The connectivity (CV=0.3) for the intermediates miRNAs was between both main groups (Figure S11).

**Figure 9:**
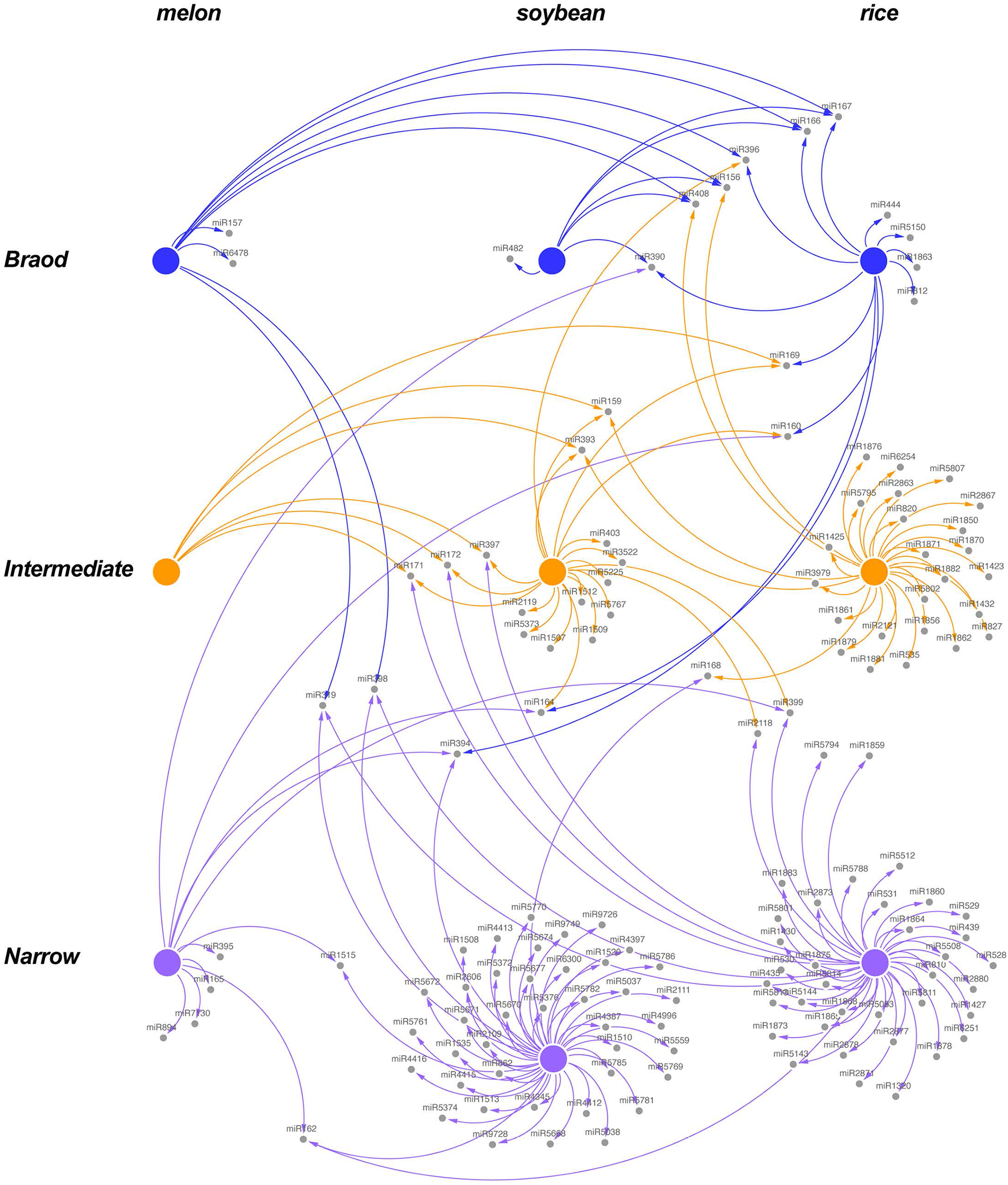
The general architecture of the miRNA-mediated response to stress is conserved in other crops. Graphic representation of the connectivity between stress responsive miRNAs identified in melon, soybean and rice according it categorized range of response to stress. Interactions between stress responsive miRNAs support that the functional categorization of the stress responsive miRNA families is comparable in the three analyzed crops. Nodes represent functional groups (Broad - blue, Intermediate - orange and Narrow – magenta) of stress responsive miRNAs in melon, soybean and rice. Edges represent weighted associations between miRNAs based on response-range in each analyzed crop.

## DISCUSSION

MiRNAs fine-tune the transcriptional response triggered in plants exposed to stress (Sunkar et al., 2012; Nogoy et al., 2018). Although the role of miRNAs is well established there was a question that remained unsolved: what kind of miRNA-based regulatory architectures do plants use to connect miRNAs and plant responses with the surrounding environment? (Reis et al., 2015; Zhang, 2015). A recent work provided evidences supporting that stress-responsive miRNAs in melon plants exposed to diverse stress conditions were hierarchically structured (Sanz-Carbonell et al., 2019). However, this previous analysis could not exclude that this structural organization was biased due to the lack of the time component in the experiment. Here, we have addressed this question by analyzing the dynamic evolution of the miRNA-mediated response to stress in melon plants at earlier phases of the exposition to stress (two -T1- and four -T2-days after stress exposition). Our analysis identified 20 stress-responsive miRNA families (14 miRNAs common for both T1 and T2, 3 restricted to T1, and 3 to T2) that mainly target well-described TFs (SPL, BEH4, ARF, ATHB14, TCP, AP2, GRF and NFY). This agrees with the observed in diverse plants (arabidopsis, rice, maize, sorghum, sunflower, etc.) where it is well established that that many of the stress-responsive miRNAs target TFs (Sunkar et al., 2007; Kumar, 2014; Zhang, 2015; Samad et al., 2017). This data contributes to reinforce the notion that in plants the role of miRNAs during response to stress is conserved (Rubio-Somoza and Weigel, 2011; Megraw et al., 2016; Zhao et al., 2016).

Following the identification of stress-responsive miRNA families we inferred the structure of this response. Our results, obtained by clustering and network analysis, revealed that miRNAs are hierarchically organized in 3 well-defined functional groups:

1) miRNAs showing a broad response-range share response with many other miRNAs, and were central elements in the network, while 2) miRNAs with a narrow response-range appear in the periphery of the network and exhibit a lower connectivity and 3), intermediate miRNAs, that due to their bordering condition, show a less consistent range of response to stress along the analyzed period. This result complemented our previous data from plants exposed to a longer stress period (Sanz-Carbonell et al., 2019). Indeed, the dynamic analysis of the structure of stress-responsive miRNAs using three time points (termed T1, T2 and T3) evidenced that the majority of miRNAs clustering in the broad or narrow response-range maintain this clustering along time. Indeed ≈ 70% of the miRNAs of narrow response-range-in at least one of the three analyzed times- were strictly limited to this group along the entire period of exposition to stress (boxed miRNAs in Figure 8) and only two (miR319 and miR157) of the 10 miRNAs that cluster in the broad response-range group change their clustering state to narrow response-range during time. Altogether these results showed that, although the global expression of stress-related miRNAs is expected to be dynamic and susceptible to the duration of the stress, the basic architecture of the miRNA-mediated network was conserved, at least under the analyzed period.

Interestingly, miRNAs with broad or narrow response-ranges regulate genes involved in different processes. While narrow-response miRNAs regulated genes associated to stress response and basic cell functions, broad response-range targeted master regulators of central hubs, predominantly (70% of the identified targets) TFs related with plant development (Table S7).

Narrow-response miRNAs include miR395, miR894 or miR394, which have been previously associated to diverse biotic and/or abiotic stress responses (Kawashima et al., 2011; Yuan et al., 2016; Du et al., 2018; Xie et al., 2014; Song et al., 2013). On the other hand, broad-response miRNAs regulate genes involved in the responses to various biotic and abiotic stresses in different species and include miR396, miR156, miR166 and miR167. Several studies have shown that the miR396-GRF module is involved in the responses to various biotic and abiotic stresses, including drought, salt, alkali, UV-B radiation, osmotic stresses, and pathogen infection (Gao *et al*., 2010; Kim *et al*., 2012; Casadevall *et al*., 2013; Chen *et al*., 2015). Interaction between miR156 and SPL improves salinity (Arshad *et al*., 2017 a), drought (Arshad *et al*., 2017 b) and heat (Matthews *et al*., 2019) tolerance in *Medicago sativa* and modulate the response to low temperatures in different plant species, including monocotyledonous, dicotyledonous, and gymnosperms (Zhou and Tang, 2019). In addition was also demonstrated that miR156 regulate immunity to bacterial infection in *N. benthamiana* (Padmanabhan *et al*. 2013). Increasing evidences suggest that miR166 family might play crucial roles in response to abiotic and biotic stresses. For example, miR166 up-regulation upon salinity stress was observed in potato (Kitazumi *et al*., 2015) and cold tolerance in soybean (Li *et al*., 2017). Similarly, miR166 was induced by *Phytophthora sojae* infection in soybean, indicating that it may be implicated in basal defense processes (Wong *et al*., 2014). Finally miR167-mediated regulation of ARFs has been involved in response a diverse stress conditions in barley (Kantar *et al*., 2010), arabidopsis (Liu *et al*., 2008), populous (Jia *et al*., 2009), triticum (Kantar *et al*., 2011) and rice (Zhou *et al*., 2010).

Additionally, we validated our analysis focusing in the role of two miRNAs (miR398 and miR408) known as copper-related and behaving as generalists in our analysis. These miRNAs show a highly coordinated expression pattern throughout the analyzed period. Cu-miRNAs belong to a conserved family of miRNAs that are involved in the regulation of Cu-related proteins and Cu homeostasis (Burkhead *et al*., 2009). However the existence of a large body of evidence linking differential expression of Cu-miRNAs to the response to abiotic (Dugas and Bartel, 2008; Li *et al*., 2010; Kantar *et al*., 2010; Trindade *et al*, 2010; Guan *et al*., 2013; Jovanovic *et al*., 2014; Liang *et al*., 2015) and biotic (Ravet *et al*., 2011; De Luis *et al*., 2012; Naya *et al*., 2014; Thiebaut *et al*., 2014; Xu *et al*., 2014) stress conditions argues against the idea that these miRNAs may have evolved exclusively as an adaptation to Cu-limited soils (Pilon, 2017). Interestingly, it has been proposed that cu-miRNAs may act as mobile (at both cell-to-cell and long distance levels) signals able to modulate at local and systemic level the distribution of Cu in the cell allowing plants to coordinate development and response to stress (Pilon, 2017). The highly synchronized expression pattern observed for both miR398 and miR408 and their copper-related targets, during the continuous stress treatment performed in melon plants is in coincidence with this possibility.

The analysis of the action range and function of stress-responsive miRNAs lead us to speculate about a potential selective stress-related regulatory activity in melon, at least under the range of stress conditions analyzed in this work. Taking into account our results, it is possible to envision a scenario, where a change in the expression of broad-response miRNAs can have a large impact on plant physiology through the regulation of TFs working as amplifiers of the stress response (Megraw et al., 2016; Samad et al., 2017; Hernández and Sanan-Mishran, 2017). Additionally, TFs and miRNAs can regulate each other establishing feedback and feedforward loops (Rubio-Somoza and Weigel, 2011; Megraw et al., 2016), and thus increasing the potential regulatory role for these broad response-range miRNAs as key modulators of the interaction between plant and environment.

All together, our results evidence the existence of a complex, hierarchical and dynamic miRNA-mediated regulatory network in melon that underlies the responses to environmental stimuli. On the other hand, the observation that the stress response mediated by miRNAs in rice and soybean (monocotyledonous and dicotyledonous species, respectively) maintain the global structure (at both functional and architectural levels) observed in melon plants, evidence that the general structure of the miRNA-network of response to stress inferred in melon, is a non exclusive feature for this crop. Our work may contribute to elucidate how miRNA-mediated regulatory pathways control gene expression and functionally connect plant responses with stress conditions. This knowledge is pivotal for a better understanding of the molecular mechanisms that enable plants to respond and eventually adapt to the environmental changes (Carrera et al., 2009).

## EXPERIMENTAL PROCEDURES

### Plant material, growth conditions, and stress treatments

Melon seeds of cv. *Piel de Sapo* were germinated in Petri dishes at 37°C/48h darkness followed by 24h/25°C (16/8 light/darkness). Melon seedlings were sown in pots and maintained for 10 days under controlled conditions ( 28°C/16h light and 20°C/8h darkness) in a growing chamber. Only plants homogeneously developed were selected for stress treatment. At day 11, selected plants were exposed to the seven biotic and abiotic stress treatments, as previously described (Sanz-Carbonell et al., 2019) (detailed in the Table S1). At two (T1) and four (T2) days post-treatment, the first leaf under the apical end per plant was collected in liquid nitrogen and maintained at –80°C until processing. Each analyzed sample corresponds to a pool of three treated plants. Three biological replicates were performed for each treatment. Leaves recovered from non-treated melon plants at the starting-point of the experiment were considered as T0.

### RNA extraction and small RNA (sRNA) purification

Total RNA was extracted from leaves (~0.1 g) recovered from treated and control melon plants using the TRI reagent (SIGMA, St. Louis, MO, USA) according to the manufacturer’s instructions. The low-molecular weight RNA (<200 nt) fraction was enriched from total RNA using TOTAL-miRNA (miRNA isolation Kit, REAL) according to the manufacturer’s instructions.

### sRNA sequencing

Production and sequencing of the libraries were carried out by SISTEMAS GENOMICOS (https://www.sistemasgenomicos.com). Fifty one (24 each for T1 and T2 and three for T0) cDNA libraries were obtained by following ILLUMINA’s recommendations. Briefly, 3’ and 5’ adaptors were sequentially ligated to the RNA prior to reverse transcription and cDNA generation. cDNAs were enriched by PCR to create the indexed double stranded cDNA library. Size selection was performed using 6% polyacrylamide gel. The quantity of the libraries was determined by quantitative real-time PCR (qRT-PCR) in a LightCycler 480 (ROCHE). Prior to cluster generation in cbot (ILLUMINA), an equimolar pooling of the libraries was performed. The pool of the cDNA libraries was sequenced by paired-end sequencing (100 bp each) in a HiSeq 2000 (ILLUMINA). Adaptors and low quality reads were trimmed by using the *Cutadapt* software (Martin, 2011). To analyze the dynamic evolution of the miRNA-mediated response along a determined period of time exposition to adverse environmental conditions, we also include in this study previous data (Sanz-Carbonell et al., 2019) obtained from melon plants exposed to identical stress conditions during 11 days and identified here as T3.

### Bioinformatic analysis of miRNA sequences

We use pair-wise Spearman rank to infer the correlation rate between sequencing data obtained from the different conditions analyzed and their biological replicates using the *stats* R-package (R Core Team, 2013). Spearman correlation coefficient ρ (rho) was estimated to determine the linear association between all libraries with “cor” function.

Differential expression of melon sRNAs was estimated by three different statistical testing methods: *NOISeq* (Tarazona et al., 2015), *DESeq2* (Love et al., 2014), and *edgeR* (Robinson and Oshlack, 2010) R-packages for pairwise differential expression analysis of expression data. Differentially expressed sRNAs were filtered using three criteria: *i*) log2-fold change (log2FC) ≥1 or ≤-1 for biological significance, *ii*) P value <0.05, and *iii*) base mean ≥5, which is the mean of normalized counts of all samples. Small RNAs identified as differentially expressed by the three methods were aligned against miRNA sequences in miRBase (release 22) (Kozomara and Griffiths-Jones, 2014), using *blastall* software by command line in Linux with the number of mismatches allowed set to zero. Sequences fully homologous to previously described mature miRNAs were identified as known stress-responsive miRNAs. Heat maps were generated with log_2_FC values (obtained by DESeq2 analysis for the most frequent sequence in each miRNA family) using the *gplots* R package.

### qRT-PCR assays

We selected to validation by qRT-PCR, differential and non-differential melon miRNAs that exhibit a relatively higher accumulation ratio (base mean ≥50) in order to guaranty their appropriate detection. Quantification of selected miRNAs was performed starting from low-molecular weight RNA (< 200 nt) fractions obtained as described above. A slightly modified stem-loop-specific reverse transcription protocol for miRNAs detection (Czimmerer et al., 2013) was performed as previously described (Sanz-Carbonell et al., 2019). Primers used are listed in Table S11.

To analyze target expression, total melon RNA (1.5 μg) was subjected to DNase treatment (EN0525, Thermo Scientific™) followed by reverse transcription using RevertAid First Strand cDNA Synthesis Kit (Thermo Scientific™) according to the manufacturer’s instructions for use with oligo-dT. cDNAs were amplified by conventional end point RT-PCR using specific primers to assess for sequence specificity. Then, real-time PCR was performed as described previously (Bustamante et al., 2018). All analyses were performed in triplicate on an ABI 7500 Fast-Real Time qPCR instrument (Applied Biosystems) using a standard protocol. The efficiency of PCR amplification was derived from a standard curve generated by four 10-fold serial dilution points of cDNA obtained from a mix of all the samples. RNA expression was quantified by the comparative ΔΔCt method. Primers used are listed in Table S11.

Relative RNA expression was determined by using the comparative ΔΔCT method (Livak and Schmittgen, 2001) and normalized to the geometric mean of Profilin (NM_001297545.1) expression, as reference. The statistical significance of the observed differences was evaluated by the paired t-Test.

### Degradome assay

Libraries for high-scale degradome assay were constructed following the protocol previously described (Zhai et al., 2014) with minor modifications. Poly (A+) RNA (≈200 ng) purified from total RNA using the Oligotex mRNA kit (Qiagen) was used. A 5’ RNA oligonucleotide adaptor (PARE 5’ Adaptor) containing a *Mme* I recognition site was ligated to the 5’-phosphate of the truncated poly (A+) RNA by T4 RNA ligase. The ligated products were purified by ethanol precipitation and subjected to a reverse transcription reaction with an oligo dT primer with a known tail (dT primer). Following RNA degradation by alkaline lysis, the cDNA was amplified by PCR using adaptor and oligo dT-specific primers (RP1S and GR3’, respectively), digested with *Mme* I and ligated to a 3’ double DNA adaptor (dsDNA-top and dsDNA-bottom). The ligated products were separated by electrophoresis in polyacrylamide, and those with the expected size were gel-purified and amplified with 30 PCR cycles to finally fill out the sequence of the Illumina 5’ (RP1M) and 3’ adaptors (RPIndex). PCR products of the expected size were gel-purified and subjected to sequencing by Illumina technology. Primers used are listed in Table S11.

For determination of miRNAs response range Principal Component Analysis (PCA) was carried out. First stress-responsive miRNAs were organized in a binary table of presence and absence (Table S12), in which the values “1” and “0” represent whether or not, respectively, a miRNA is reactive (with both either increased or decreased expression) to a stress condition. Next, the *prcomp* function was used to compute principal components (scale = T, center = F), included in *stats* R-package (R Core Team, 2013). The statistical significance was estimated with a Mann-Whitney-Wilcoxon test. Analyses were performed with Euclidean distance metric and Ward linkage with the “ward.D2” algorithm.

### Analysis of miRNA-mediated networks

The network represents the relationship between stress-responsive miRNAs and common stress conditions at both analyzed times (T1 and T2). The miRNAs relationship was established by pairing miRNAs and the common stresses they share. The nodes of the network correspond to the miRNAs and the edges represent the common stresses between both miRNAs. To build the network, two input tables were needed: the one containing information about the nodes (Table S13) and the other, about the edges (Table S14). The network was visualized with the *d3Network* package from R (Pink and Vogel, 2014), creating a D3 JavaScript network useful for studying complex networks and integrating them with any type of attribute data, such that nodes represent families of responsive miRNAs and edges link miRNAs that respond to at least one common stress. To account for the strength of the link between two miRNAs (i.e., the number of stresses to which they both respond), we made the thickness of the edge proportional to the number of stresses in common between linked miRNAs with the thickness of the edge being greater for miRNAs sharing more stresses (divided by the number of total stresses). The size of the nodes is a visual representation of those considered as broad, intermediate, and narrow response range miRNAs.

With the aim of measuring the centrality and influence of a miRNA or a group of miRNAs in a common context, the *igraph* package from R (Csárdi and Nepusz, 2006) was used to calculate the betweenness centrality for all nodes of the network. The statistical significance of the differences between the different groups (broad, intermediate, and narrow response range) was estimated with a Mann-Whitney-Wilcoxon test.

### Time course of miRNA expression

To analyze melon miRNAs reactive to stress under a dynamic viewpoint, we performed a clustered time-course analysis of miRNA expression profiles. For this, counts of the reads for each miRNA were converted to RPM. The expression values at each time point were normalized to the corresponding expression value of their control (except for T0 that was normalized to themselves) and Log2-transformed.

The complete method in the *NbClust* R-package considering Euclidean distances was used to identify the optimal number of clusters (Charrad et al., 2012), the *mode* of the clusters value (most representative number of clusters in that the miRNA expression profiles were grouped for each stress condition) was selected as the optimal input parameter for the number of cluster. Soft clustering analysis was performed with the function “mfuzz” of *Mfuss* R-package (Kumar and Futschik, 2007), the number of iterations was set at one million. The resulting clusters (grouping similar dynamic miRNA expression profiles in each stress condition) were generated by *ggplot2* R-package.

To evaluate common dynamic patterns of miRNA expression during the response to stress, we used clustering data to construct a cluster-related non-directional network. We searched for every pair of miRNAs (source-target), the number of clusters in which both were grouped (two miRNAs sharing a particular cluster were considered connected). In this network, built with *Cystoscape* software (Shannon et al., 2003), nodes represented stress-responsive miRNAs (source in Supplementary Table 4) and edges indicate the links between miRNAs with common clustering in response to stress (link value in Supplementary Table 5). The nodes size and links thickness were estimated as is described in the Supplementary Table 4 and 5.

### Data analysis and visualization software

The local computational analyzes were carried out on an own server based on Ubuntu 16.04.5 LTS (Intel® Xeon (R) CPU E5-2630 v4 @ 2.20GHz × 20 & 32 GB of RAM 2133MHz). The adapters were removed by Cutadapt v.1.10 (https://cutadapt.readthedocs.io/en/stable/guide.html) and FastQC v.011.3 (www.bioinformatics.babraham.ac.uk/projects/) for some quality control checks. The alignments of the sequences of sRNAs were made with Blastall v. 2.2.26 (http://gensoft.pasteur.fr/docs/blast/2.2.26) and the representation of the network of clusters with Cystoscape v.3.4 (https://cytoscape.org/).

Most of analysis and visualization were performed using R v.3.4.4, assisted by the Rstudio IDE (Integrated Development Environment for R v.1.0.153; https://www.rstudio.com/) as well as the following R-packages.

#### For differential expression analysis

DESeq2 v.1.18.1 (https://bioconductor.org/packages/release/bioc/html/DESeq2.html) edgeR v.3.20.9 (https://bioconductor.org/packages/release/bioc/html/edgeR.html) NOISeq v.2.22.1 (http://bioconductor.org/packages/release/bioc/html/NOISeq.html) Reshaping and statistical functions for data.tables in R and: reshape2 v.1.4.3 (https://cran.r-project.org/web/packages/reshape2/index.html) stats v.3.4.4 (included in R)

#### For cluster analysis

cluster v.2.07-1 (https://cran.r-project.org/web/packages/cluster/index.html) Factoextra v.1.0.5 (https://cran.r-project.org/web/packages/factoextra/index.html) NbClust v.3.0 (https://cran.r-project.org/web/packages/NbClust/index.html) Mfuzz v.2.38.0 (https://bioconductor.org/packages/release/bioc/html/Mfuzz.html)

#### For the generation of images and functions extended

ggExtra v.0.8 (https://cran.r-project.org/web/packages/ggExtra/index.html) ggplot2 v.3.0.0 (https://cran.r-project.org/web/packages/ggplot2/index.html) ggrepel v.0.8.0 (https://cran.r-project.org/web/packages/ggrepel/index.html) ggthemes v.3.5.0 (https://cran.r-project.org/web/packages/ggthemes/index.html) gplots v.3.0.1 (https://cran.r-project.org/web/packages/gplots/index.html) gridExtra v.2.3 (https://cran.r-project.org/web/packages/gridExtra/index.html) venn v.1.6 (https://cran.r-project.org/web/packages/venn/index.html)

#### For viewing and exporting JavaScript networks

d3Network v.0.5.2.1 (https://cran.r-project.org/web/packages/d3Network/index.html) and rgl v.0.99.16 (https://cran.r-project.org/web/packages/rgl/index.html).

### Accession Numbers

Melon miRNA sequences used in this study (T0, T1, T2 and T3) have been submitted to the genomic repository SRA of the NCBI and are available in the BioProject (PRJNA551387).

## ACKNOWLEDGMENTS

The authors thank Dr. A. Monforte (IBMCP) and Dra. B. Pico (Cucurbits Group - COMAV) for providing melon seeds and *Monosporascus* isolate respectively. Degradome sequencing was performed by the SNP&SEQ Technology Platform in Uppsala. The facility is part of the National Genomics Infrastructure (NGI) Sweden and Science for Life Laboratory. The SNP&SEQ Platform is also supported by the Swedish Research Council and the Knut and Alice Wallenberg Foundation.

## DISCLAIMER

The funders had no role in study design, data collection and analysis, decision to publish, or preparation of the manuscript.

