## Supplemental Figures 1 to 9 for "Dynamic architecture and regulatory implications of the miRNA network underlying the response to stress in melon"

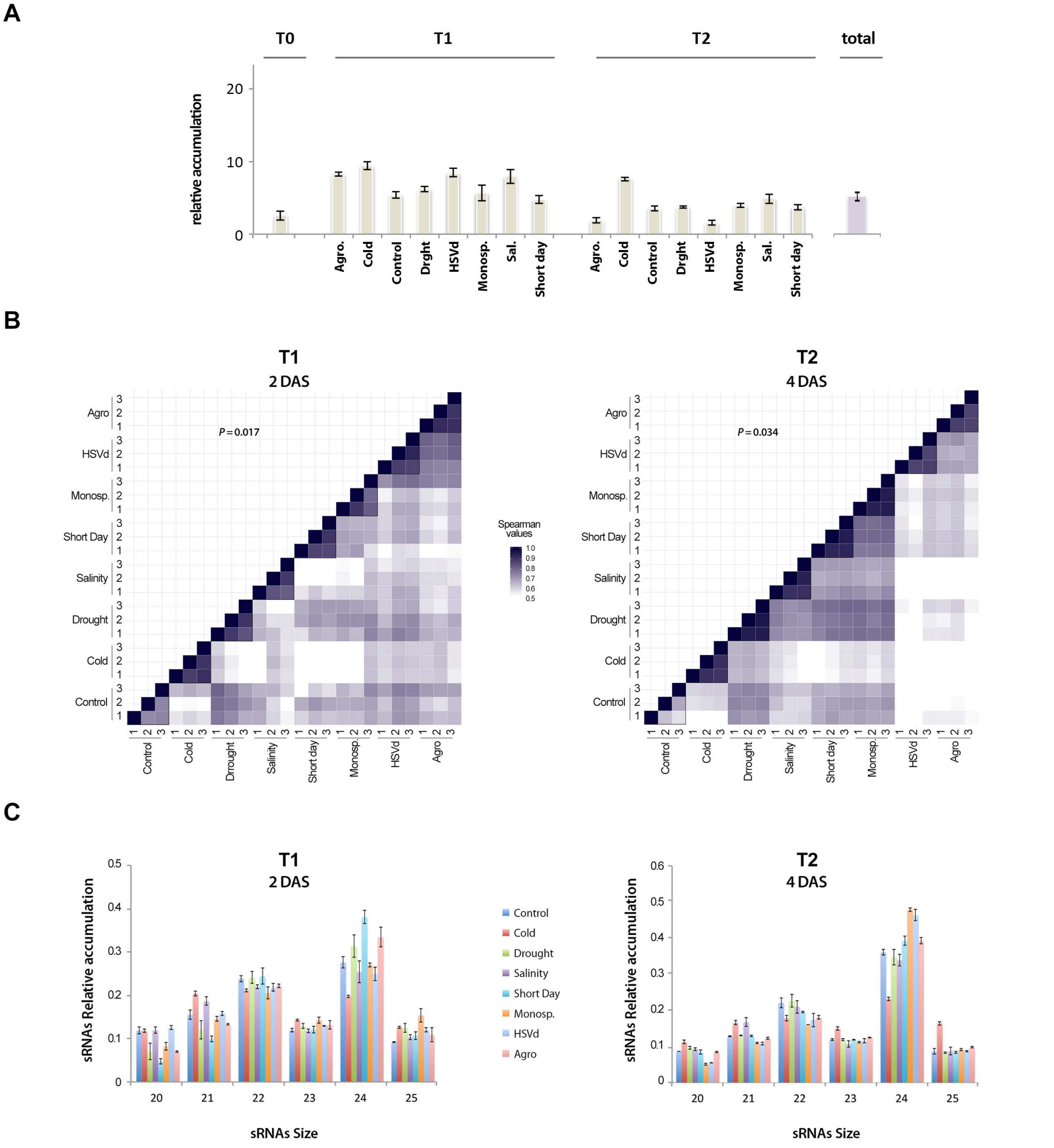


**Figure S1: Analysis of the sRNA population recovered from analyzed melon libraries. A)** Diagram showing the means of the accumulation of rRNA, tRNA, snoRNA and snRNA derived sRNAs in all analysed samples. In magent is represented the means of all analized samples. Error bars show the confidence interval of the difference between means. **B)** Heat map showing the pair-wise Spearman rank correlation between the sRNAs read of the 24 libraries analyzed at T1 (left) and T2 (right). As is showed in the figure, high correlation values (>0,75) were obtained between replicates (identified as 1, 2 and 3) of the same treatments. The color key represents the correlation coefficient range. **C)** Diagram showing the relative accumulation and distribution of the total clean reads of melon sRNAs ranging between 20 and 25 nt obtained from the sequenced libraries at T1 and T2. The control and the different analyzed treatments are represented with colors. The shown values represent the mean of all repetitions. Bars represent the standard error between replicates values.

**A**

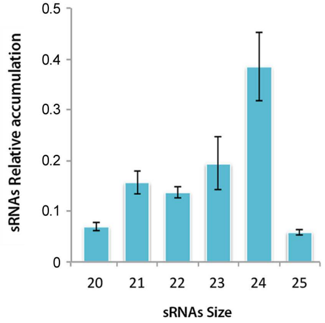


**B**

| sRNA size | Percentage T0-R1 | Percentage T0-R2 | Percentage T0-R3 | Means | SE |
|---|---|---|---|---|---|
| 20 | 7,35 | 5,46 | 7,97 | 6,93 | 0,76 |
| 21 | 15,79 | 11,55 | 19,55 | 15,63 | 2,31 |
| 22 | 15,85 | 12,25 | 13,05 | 13,72 | 1,09 |
| 23 | 29,41 | 16,67 | 12,05 | 19,38 | 5,20 |
| 24 | 25,38 | 47,38 | 42,59 | 38,45 | 6,69 |
| 25 | 6,22 | 6,68 | 4,78 | 5,89 | 0,57 |

**Figure S2. A)** Histogram showing the relative accumulation and distribution of the total clean reads of melon sRNAs ranging between 20 and 25 nt obtained from the sequenced libraries at T0. The control and the different analyzed treatments are represented with colors. Bars represent the standard error between replicates values. **B)** Details of the data graphied in A.

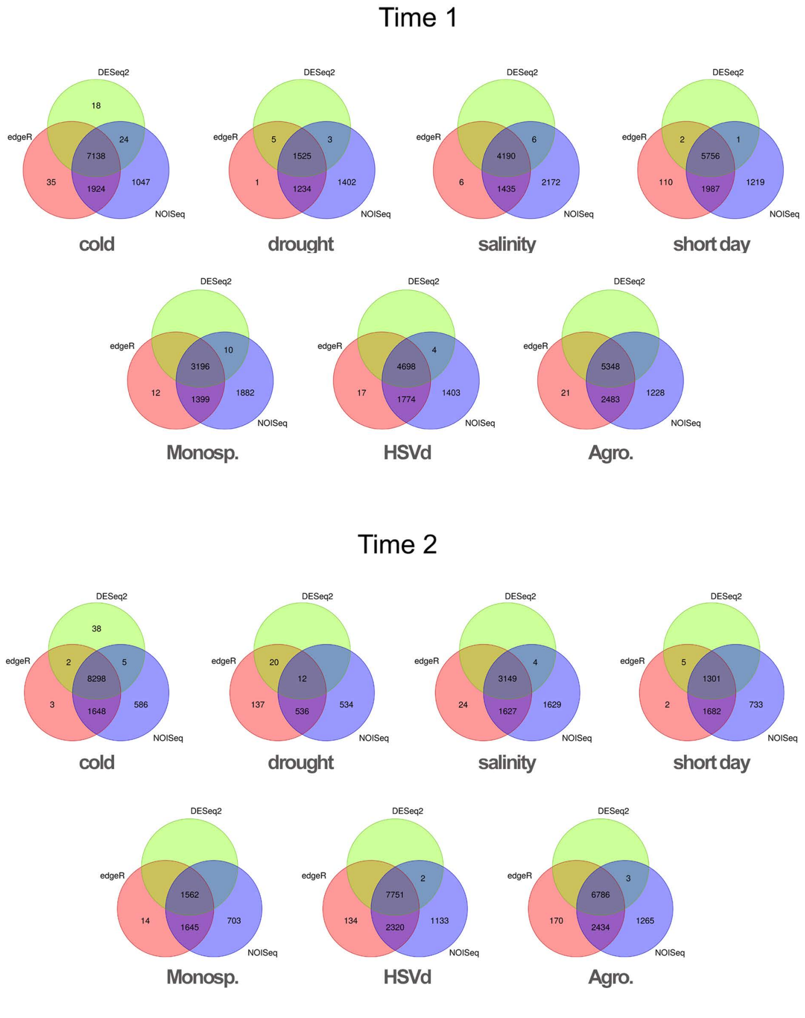


**Figure S3:** Venn diagram comparing the number of the differential sRNAs -estimated by DESeq2 (green), edgeR (red) and NOISeq (blue)- expressed in melon in response to cold, drought, salinity, short day, Monosporascus (Monosp.), HSVd and Agrobacterium (Agro.) treatments, at T1 and T2. Only the sRNAs predicted as differential by all three analysis methods were considered as true stress-responsive miRNAs.

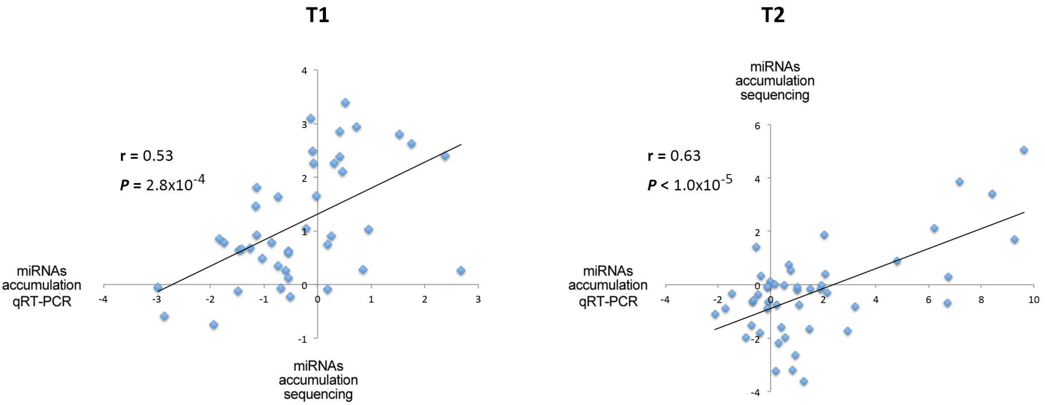


**Figure S4:** Validation of sequencing data by stem-loop qRT-PCR assay. Scatterplot showing the significant positive correlation -estimated by Pearson correlation coefficient- between the accumulation data obtained by sequencing and qRT-PCR for representative miRNAs in all analyzed stress conditions in T1 and T2 (detailed information Table S17).

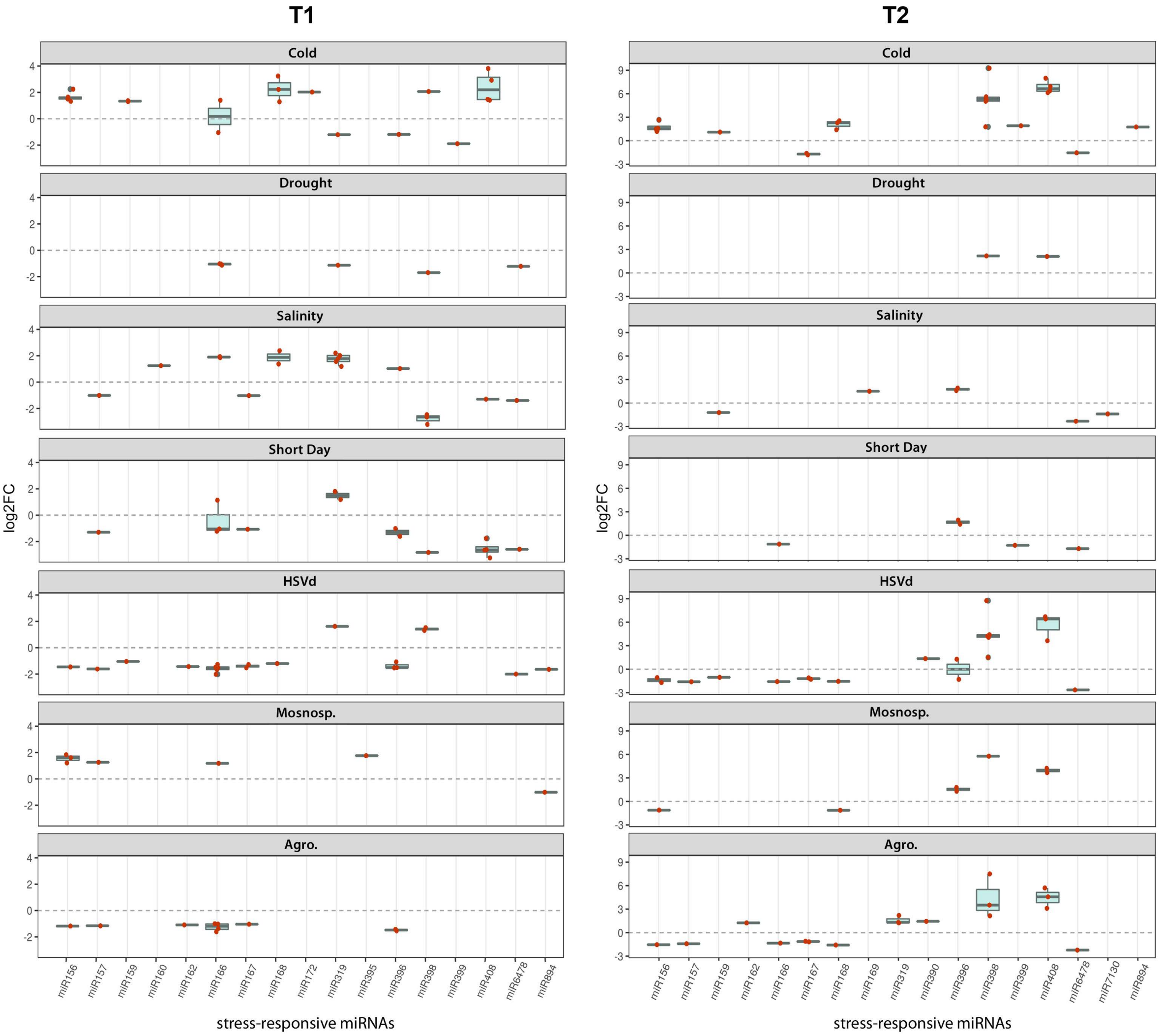


**Figure S5:** To determine the general sense of the expression for each miRNA family in both analyzed times, T1 and T2, we employed the median value of expression (represented by internal box-line) estimated by box-plot analysis of all family-related sequences (red dots). The differential expression values represented in the figure correspond to the log2FC obtained by DESeq2 analysis. In those cases with bordering discordant expression (miR166 in short day and cold treatment T1, and miR366 in HSVd infection) we considered the expression value of the most representative sequence.

**A**

| miRNA | Target | expect | start | end | Target description |
|---|---|---|---|---|---|
| *miR7130* | MU54233 | 3.5 | 798 | 821 | PREDICTED: Cucumis melo COP9 signalosome complex subunit 6a (LOC103490191), transcript variant X1, mRNA |
| *miR399* | METC013060 | 4.0 | 68 | 89 | PREDICTED: Cucumis melo beta-glucosidase 11-like (LOC103494776), transcript variant X2, mRNA |
| *mir894* | METC023591 | 4.5 | 244 | 263 | PREDICTED: Cucumis melo cysteine-rich and transmembrane domain-containing protein A-like (LOC103489653) |

**B**

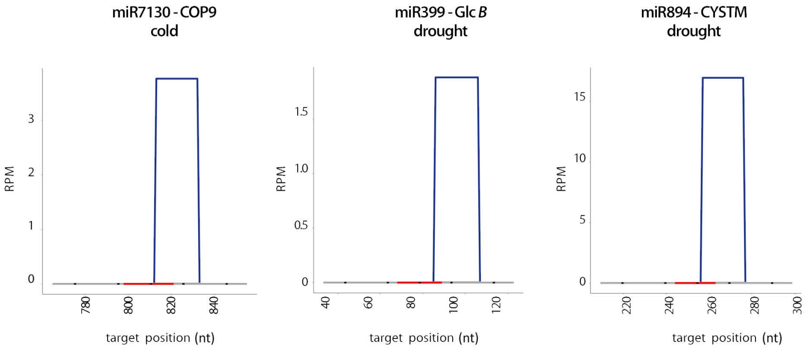


**Figure S6:** Validation of predicted targets for miR7130, miR399 and miR894 in melon. **A)** Detail of the targets prediction. The parameter expect represent the values asigned by psRNA*target* to the interaction miRNA/trancript. **B)** Graphic representation of miRNA-cleaved transcripts detected by high-scale degradome assay in representative stress-exposed samples. The obtained sequences were plotted (allowing 100% homologous matching) onto the predicted-targets sequences. The red lines on the X-axis indicate the position of the predicted miRNA recognition site in the melon transcripts. The values on the Y-axis represent the number of obtained reads (normalized in reads per million).

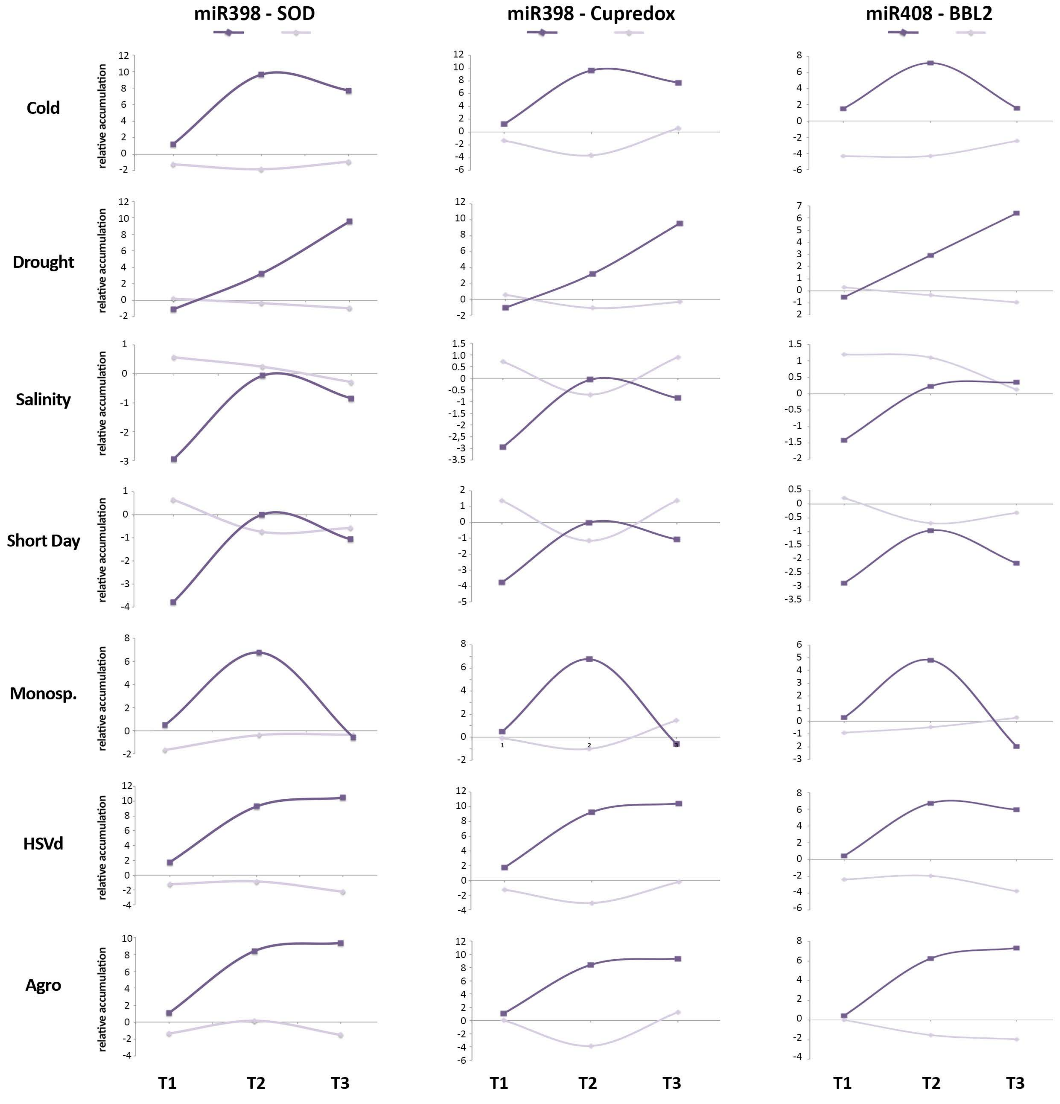


**Figure S7:** Graphic representation of comparative time course of expression levels of miR398 and miR408 (determined by sequencing) and their transcripts targets in melon (inferred by qRT-PCR) during each stress treatment analyzed in this work. The values on the Y-axis represent the relative accumulation (expressed in Log2FC), in the X-axis is represented the temporal scale of the stress treatments. SOD: (Cu) Superoxide dismutase, BBL2: Basic blue protein-like.

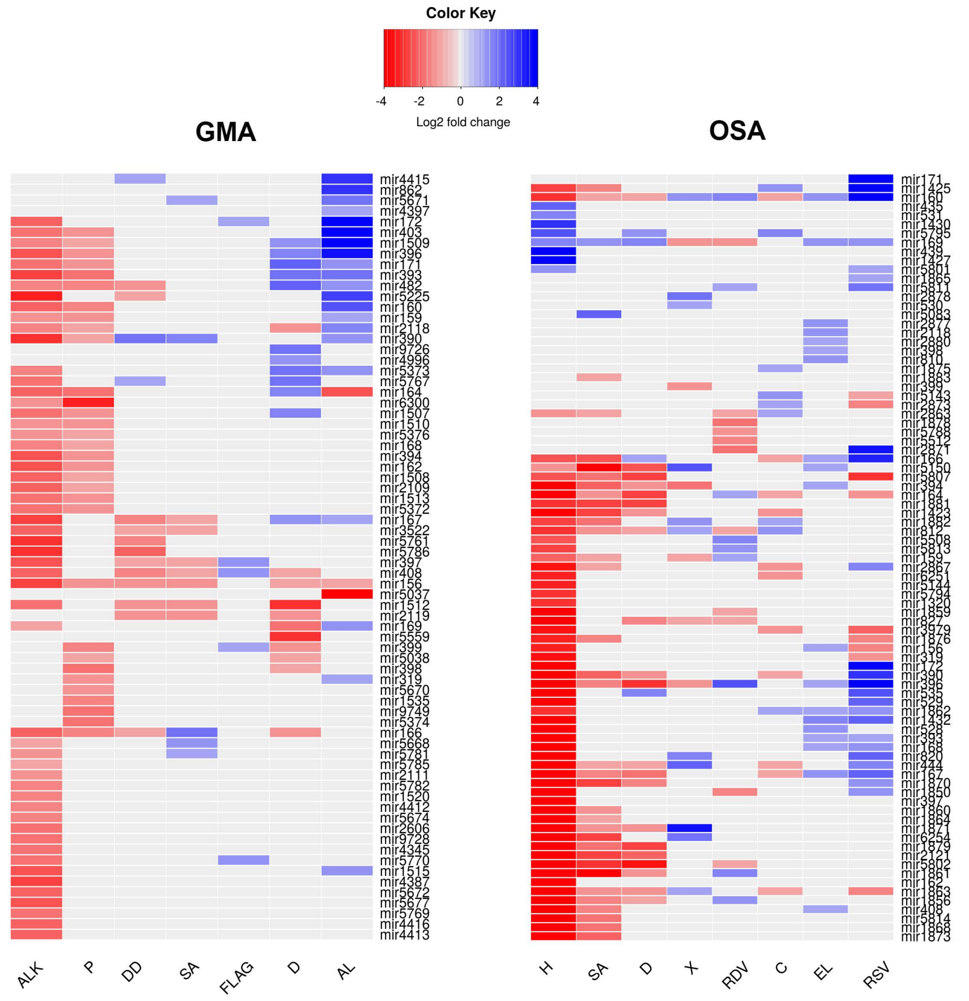


**Figure S8:** Heat map of the 72 and 85 miRNAs differentially expressed in soybean (GMA) and rice (OSA) plants respectively in response to tested stress conditions. The differential expression values represented correspond to the median of the Log2FC values obtained by edgeR analysis in each miRNA family. Stress treatments. **GMA:** ALK (alkalinity,), P (*Pseudomona sojae*), DD (moderate drought tratment), SA (salinity), FLAG (flageline treatment), D (strong drought treatment), AL (aluminiun). **OSA:** H (heat), SA (salinity), D (drought), X (*Xanthomonas oryzae*), RDV (*Rice dwarf virus*), C (cold), EL (fungal elicitor), RSV (*Rice stripe virus*). Detailed information about the stress tretaments should be recovered from the refrences listed in the Table S8.

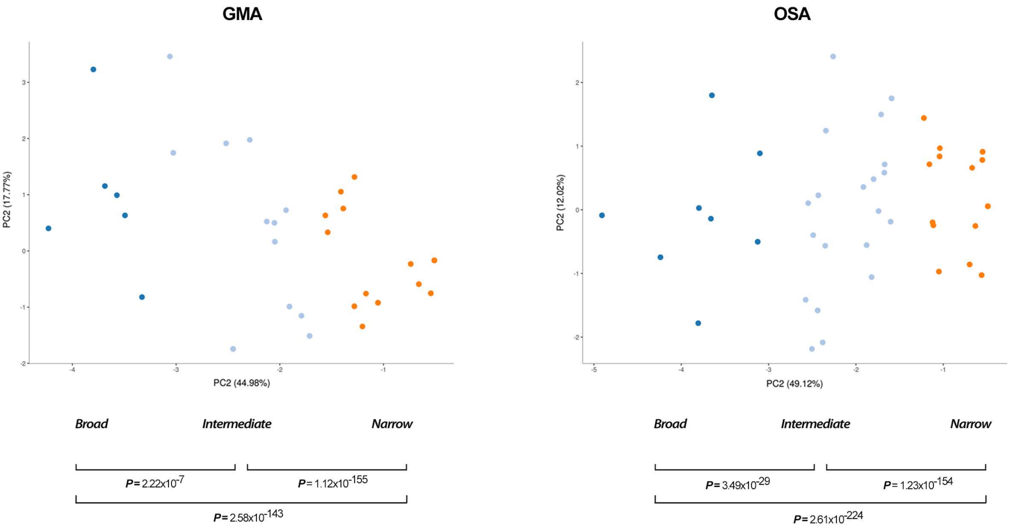


**Figure S9: Soybean and rice stress-responsive miRNAs are hierarchically organized in relation to its response-range to stress.** Principal component analysis of stress-responsive miRNAs detected in soybean (GMA - left panel ) and rice (OSA - right panel) plants. Values of proportion of variance for PC1 and PC2 are showed y the X and Y-axis, respectively. The statistical significance of the identified clusters was estimated by Mann-Whitney-Wilcoxon test, considering the inter- and intra-group Euclidean distances (P-values are showed in the graphic). The different groups of stress-responsive miRNAs are identified by colors: broad response-range (deep blue), intermediate (light blue) and narrow (orange).

### OSA

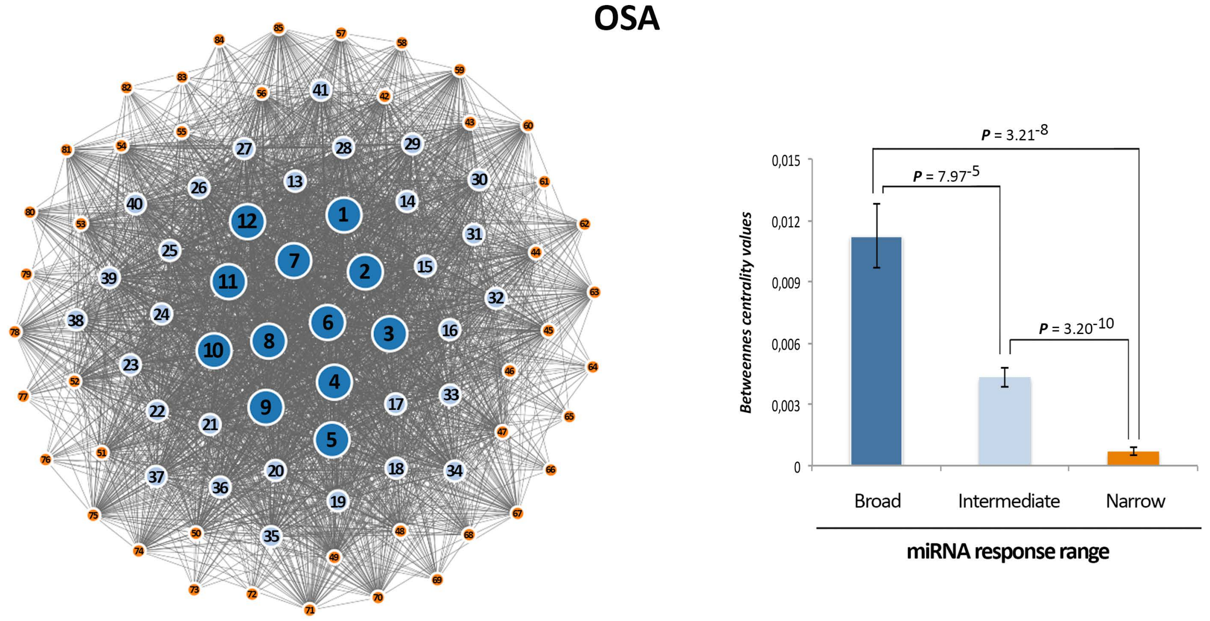


| | miRNA | Number-code |
|---|---|---|
| Broad | mir160 | 1 |
| | mir396 | 2 |
| | mir169 | 3 |
| | mir812 | 4 |
| | mir1863 | 5 |
| | mir167 | 6 |
| | mir444 | 7 |
| | mir166 | 8 |
| | mir164 | 9 |
| | mir390 | 10 |
| | mir5150 | 11 |
| | mir394 | 12 |
| Intermediate | mir1862 | 13 |
| | mir1882 | 14 |
| | mir1423 | 15 |
| | mir2867 | 16 |
| | mir1425 | 17 |
| | mir2863 | 18 |
| | mir5807 | 19 |
| | mir5802 | 20 |
| | mir1871 | 21 |
| | mir1861 | 22 |
| | mir1856 | 23 |
| | mir1870 | 24 |
| | mir827 | 25 |
| | mir159 | 26 |
| | mir5795 | 27 |

| | miRNA | Number-code |
|---|---|---|
| Intermediate | mir3979 | 28 |
| | mir2121 | 29 |
| | mir1881 | 30 |
| | mir535 | 31 |
| | mir1879 | 32 |
| | mir1432 | 33 |
| | mir408 | 34 |
| | mir393 | 35 |
| | mir168 | 36 |
| | mir156 | 37 |
| | mir1876 | 38 |
| | mir820 | 39 |
| | mir6254 | 40 |
| | mir1850 | 41 |
| Narrow | mir2873 | 42 |
| | mir5143 | 43 |
| | mir6251 | 44 |
| | mir528 | 45 |
| | mir5813 | 46 |
| | mir5814 | 47 |
| | mir5801 | 48 |
| | mir319 | 49 |
| | mir529 | 50 |
| | mir172 | 51 |
| | mir1860 | 52 |
| | mir5508 | 53 |
| | mir1873 | 54 |

| | miRNA | Number-code |
|---|---|---|
| Narrow | mir1868 | 55 |
| | mir1864 | 56 |
| | mir1859 | 57 |
| | mir5811 | 58 |
| | mir2871 | 59 |
| | mir2880 | 60 |
| | mir2118 | 61 |
| | mir810 | 62 |
| | mir398 | 63 |
| | mir531 | 64 |
| | mir397 | 65 |
| | mir162 | 66 |
| | mir5794 | 67 |
| | mir1427 | 68 |
| | mir1320 | 69 |
| | mir1430 | 70 |
| | mir439 | 71 |
| | mir435 | 72 |
| | mir1878 | 73 |
| | mir5512 | 74 |
| | mir171 | 75 |
| | mir1883 | 76 |
| | mir530 | 77 |
| | mir2878 | 78 |
| | mir1875 | 79 |
| | mir2877 | 80 |
| | mir5144 | 81 |
| | mir5788 | 82 |
| | mir1865 | 83 |
| | mir5083 | 84 |
| | mir399 | 85 |

### GMA

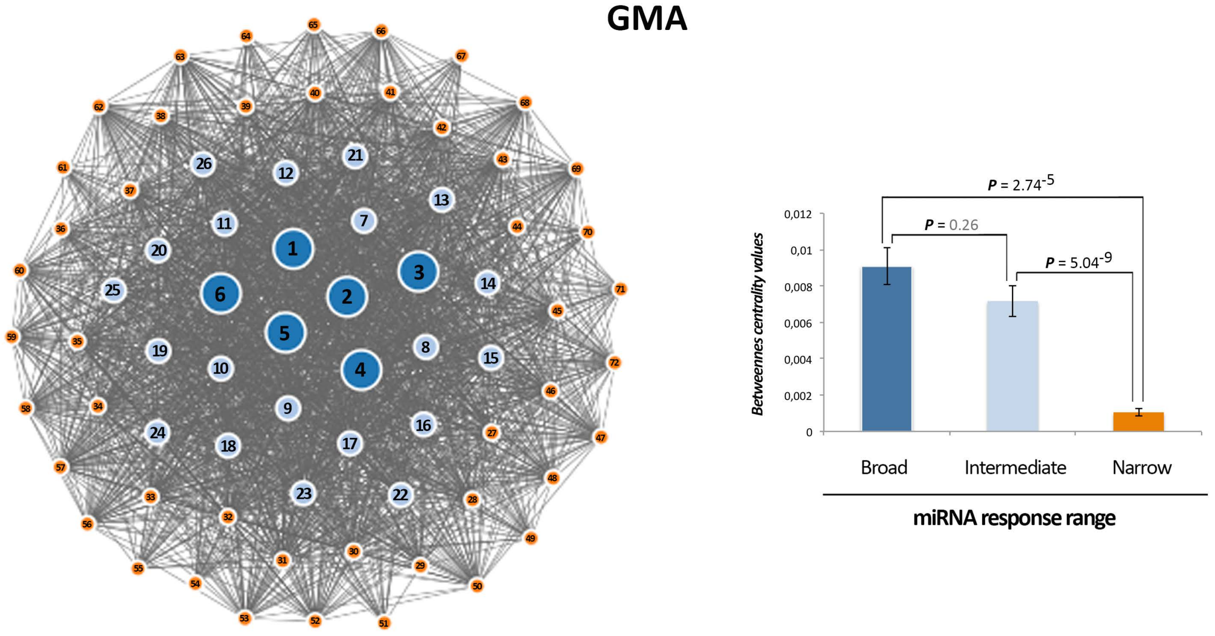


| | miRNA | Number-code |
|---|---|---|
| Broad | *mir156* | 1 |
| | *mir390* | 2 |
| | *mir482* | 3 |
| | *mir167* | 4 |
| | *mir408* | 5 |
| | *mir166* | 6 |
| Intermediate | *mir171* | 7 |
| | *mir396* | 8 |
| | *mir164* | 9 |
| | *mir1509* | 10 |
| | *mir393* | 11 |
| | *mir2118* | 12 |
| | *mir1512* | 13 |
| | *mir397* | 14 |
| | *mir169* | 15 |
| | *mir159* | 16 |
| | *mir5225* | 17 |
| | *mir172* | 18 |
| | *mir160* | 19 |
| | *mir5373* | 20 |
| | *mir403* | 21 |
| | *mir3522* | 22 |
| | *mir1507* | 23 |
| | *mir5767* | 24 |
| | *mir2119* | 25 |
| | *mir399* | 26 |

| | miRNA | Number-code |
|---|---|---|
| Narrow | *mir5671* | 27 |
| | *mir1515* | 28 |
| | *mir4415* | 29 |
| | *mir319* | 30 |
| | *mir5668* | 31 |
| | *mir394* | 32 |
| | *mir1510* | 33 |
| | *mir5781* | 34 |
| | *mir2109* | 35 |
| | *mir6300* | 36 |
| | *mir1508* | 37 |
| | *mir162* | 38 |
| | *mir168* | 39 |
| | *mir5376* | 40 |
| | *mir1513* | 41 |
| | *mir5786* | 42 |
| | *mir5770* | 43 |
| | *mir5372* | 44 |
| | *mir5761* | 45 |
| | *mir5038* | 46 |
| | *mir398* | 47 |
| | *mir4397* | 48 |
| | *mir5037* | 49 |
| | *mir2606* | 50 |
| | *mir1520* | 51 |
| | *mir5672* | 52 |

| | miRNA | Number-code |
|---|---|---|
| Narrow | *mir9728* | 53 |
| | *mir4416* | 54 |
| | *mir2111* | 55 |
| | *mir4345* | 56 |
| | *mir4413* | 57 |
| | *mir5674* | 58 |
| | *mir5782* | 59 |
| | *mir5785* | 60 |
| | *mir5769* | 61 |
| | *mir4387* | 62 |
| | *mir4412* | 63 |
| | *mir5559* | 64 |
| | *mir4996* | 65 |
| | *mir5670* | 66 |
| | *mir1535* | 67 |
| | *mir9749* | 68 |
| | *mir862* | 69 |
| | *mir5677* | 70 |
| | *mir9726* | 71 |
| | *mir5374* | 72 |

**Figure S10: Network of stress-responsive miRNAs in rice (upper) and soybean (lower).** Left panels) Nodes in the network represent differentially expressed miRNAs. Colors and numbers depict the different groups of stress-responsive miRNAs detected in melon. Node size is proportional to the number of stress conditions where a particular miRNA is differentially expressed (see Supp. File 1). Edges represent weighted associations between the terms based on response to common stress conditions. Right panels) Graphic representation of the average betweenness of the nodes calculated for broad, intermediate, and narrow response range miRNAs in rice (upper) and soybean (lower). The statistical significance of the differences was estimated by Mann-Whitney-Wilcoxon test, significant P-values are showed in bold (< 0.05), and non-significant values in gray. Error bars represent the standard error in betweenness values.

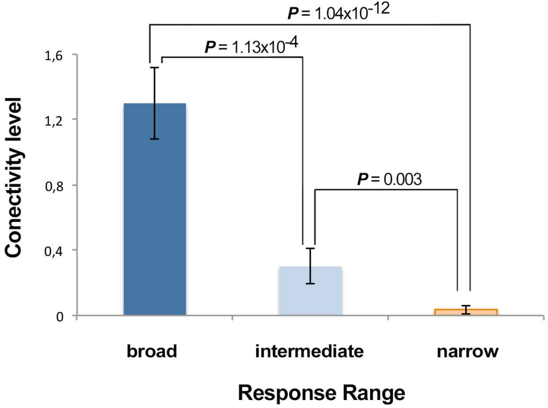


**Figure S11: Analisys of Connectivity values:** Graphic representation of the average Connectivity values (CV) (between functionally categorized miRNAs in melon, rice and soybean) obtained for stress responsive miRNAs included in the three established functional groups of response range to stress conditions analized here. The statistical significance of the differences was estimated by Mann-Whitney-Wilcoxon test, significant P-values are showed. Error bars represent the standard error in connectivity values.
