## Supplemental Tables 1 to 19 for "Dynamic architecture and regulatory implications of the miRNA network underlying the response to stress in melon"

**Table S1:** Detail of the stress treatments in melon plants. In salinity treatment, we used LiCl instead of NaCl to reduce the osmotic effect (Goossens, et al., 2000).

| Stress Conditions | Treatments at 11 days post emergency | |
|---|---|---|
| *Cold* | Irrigated with 50 ml Hoagland's solution<br>20 ºC/16 h-light --- 14 ºC/8 h-darkness | Except for drought treatment, plants were irrigated alternatively (water and Hoagland's solution) by inundation (1500 mL/48 h). |
| *Drought* | Not irrigated<br>28 ºC/16 h-light --- 20 ºC/8 h-darkness | |
| *Salinity* | Irrigated with 50 ml of LiCl (200 mM)<br>28ºC/16 h-light --- 20 ºC/8 h-darkness | |
| *Short-day* | Irrigated with 50 ml Hoagland's solution<br>28 ºC/8 h-light --- 20 ºC/16 h-darkness | |
| *HSVd* | Irrigated with 50 ml Hoagland's solution<br>Inoculated with viroid-RNA in both cotyledons<br>28 ºC/16 h-light --- 20 ºC/8 h-darkness | |
| *Monosporascus* | Irrigated with 50 ml Hoagland's solution plus *M. cannonballus* mycelium (1000 UFC)<br>28 ºC/8 h-light --- 20 ºC/16 h-darkness | |
| *Agrobacterium* | Irrigated with 50 ml Hoagland's solution<br>infiltrated in cotyledons with *A. tumefaciens* (0.8 OD)<br>28 ºC/16 h-light --- 20 ºC/8 h-darkness | |
| *Control* | Irrigated with 50 ml Hoagland's solution<br>28 ºC/16 h-light --- 20 ºC/8 h-darkness | |

**Table S2**: Detail of the sequenced reads.

| Time stress exposition | Type of stress | Analyzed samples | Filtered sRNA reads | Reads by analized time |
|---|---|---|---|---|
| **T0** | *prior to treatment starting* | *R1* | 3.216.440 | 9.693.430 |
| | | *R2* | 3.217.277 | |
| | | *R3* | 3.259.713 | |
| **T1** | *cold* | *R1* | 1.573.058 | 45.929.350 |
| | | *R2* | 1.281.676 | |
| | | *R3* | 1.802.635 | |
| | *drought* | *R1* | 1.986.110 | |
| | | *R2* | 2.201.443 | |
| | | *R3* | 1.774.052 | |
| | *salinity* | *R1* | 2.643.989 | |
| | | *R2* | 2.131.931 | |
| | | *R3* | 1.922.773 | |
| | *short day* | *R1* | 1.912.305 | |
| | | *R2* | 1.953.817 | |
| | | *R3* | 1.722.858 | |
| | *Agrobacterium* | *R1* | 2.011.023 | |
| | | *R2* | 1.608.476 | |
| | | *R3* | 2.333.327 | |
| | *HSVd* | *R1* | 1.112.168 | |
| | | *R2* | 1.579.777 | |
| | | *R3* | 1.428.136 | |
| | *Monosporascus* | *R1* | 1.636.809 | |
| | | *R2* | 2.248.825 | |
| | | *R3* | 1.324.404 | |
| | *control* | *R1* | 2.387.425 | |
| | | *R2* | 2.328.463 | |
| | | *R3* | 3.023.870 | |
| **T2** | *cold* | *R1* | 1.407.153 | 61.219.206 |
| | | *R2* | 1.422.247 | |
| | | *R3* | 1.279.661 | |
| | *drought* | *R1* | 1.861.865 | |
| | | *R2* | 2.229.770 | |
| | | *R3* | 2.613.321 | |
| | *salinity* | *R1* | 2.175.703 | |
| | | *R2* | 2.557.117 | |
| | | *R3* | 3.372.067 | |
| | *short day* | *R1* | 2.642.516 | |
| | | *R2* | 2.927.338 | |
| | | *R3* | 2.673.733 | |
| | *Agrobacterium* | *R1* | 3.055.638 | |
| | | *R2* | 3.613.207 | |
| | | *R3* | 3.118.311 | |
| | *HSVd* | *R1* | 2.990.357 | |
| | | *R2* | 3.397.996 | |
| | | *R3* | 3.344.113 | |
| | *Monosporascus* | *R1* | 2.593.347 | |
| | | *R2* | 2.351.910 | |
| | | *R3* | 3.014.594 | |
| | *control* | *R1* | 2.184.690 | |
| | | *R2* | 2.029.912 | |
| | | *R3* | 2.362.640 | |
| ***Total analized samples*** | | **51** | | |
| ***Total filtered reads*** | | | | **116.841.986** |

**Table S3:** Stress-responsive miRNA-family related sequences recovered from melon libraries.

| Time | miRNA | sRNA | nt |
|---|---|---|---|
| T1 | *miR156* | TGACAGAAGAGAGTGAGCAC | 20 |
| | | TGACAGAAGAGAGTGAGCACA | 21 |
| | | TGACAGAAGAGAGTGAGCACT | 21 |
| | | TTGACAGAAGAGAGTGAGCAC | 21 |
| | | TTGACAGAAGATAGAGAGCAC | 21 |
| | | TTGACAGAAGATAGAGGGCAC | 21 |
| | *miR157* | GCTCTCTATACTTCTGTCACC | 21 |
| | | GCTCTCTATGCTTCTGTCATC | 21 |
| | *miR159* | TTTGGACTGAAGGGAGCTCTA | 21 |
| | | TTTGGATTGAAGGGAGCTCTA | 21 |
| | | TTTGGATTGAAGGGAGCTCTT | 21 |
| | *miR160* | TGCCTGGCTCCCTGTATGCC | 20 |
| | *miR162* | TCGATAAACCTCTGCATCCAG | 21 |
| | *miR166* | CGGACCAGGCTTCATTCCCC | 20 |
| | | TCGGACCAGGCTTCATTCCC | 20 |
| | | TCGGACCAGGCTTCATTCCT | 20 |
| | | TCGGACCAGGCTTCATTCCTC | 21 |
| | | TCTCGGACCAGGCTTCATTC | 20 |
| | | TCTCGGACCAGGCTTCATTCC | 21 |
| | | TTGGACCAGGCTTCATTCCCC | 21 |
| | *miR167* | TGAAGCTGCCAGCATGATCT | 20 |
| | | TGAAGCTGCCAGCATGATCTGC | 22 |
| | *miR168* | CCCGCCTTGCATCAACTGAAT | 21 |
| | | TCGCTTGGTGCAGGTCGGGA | 20 |
| | | TCGCTTGGTGCAGGTCGGGAA | 21 |
| | *miR172* | AGAATCTTGATGATGCTGCAT | 21 |
| | *miR319* | CTTGGACTGAAGGGAGCTCCC | 21 |
| | | TTGGACTGAAGGGAGCTCCC | 20 |
| | | TTGGACTGAAGGGAGCTCCCT | 21 |
| | | TTGGACTGAAGGGAGCTCCT | 20 |
| | | TTGGACTGAAGGGAGCTCCTTC | 22 |
| | *miR395* | TGAAGTGTTTGGGGGAACTC | 20 |
| | *miR396* | TTCCACAGCTTTCTTGAACT | 20 |
| | | TTCCACAGCTTTCTTGAACTG | 21 |
| | | TTCCACAGCTTTCTTGAACTT | 21 |
| | | TTCCACGGCTTTCTTGAACTG | 21 |
| | | TTCCACGGCTTTCTTGAACTT | 21 |
| | *miR398* | TGTGTTCCCAGGTCGCCCCTG | 21 |
| | | TGTGTTCTCAGGTCACCCCT | 20 |
| | | TGTGTTCTCAGGTCGCCCCTG | 21 |
| | *miR399* | AGGGCTTCTCTCCATTGGCAGG | 22 |
| | *miR408* | ATGCACTGCCTCTTCCCTGGC | 21 |
| | | TGCACTGCCTCTTCCCTGGC | 20 |
| | | TGCACTGCCTCTTCCCTGGCT | 21 |
| | | TGCACTGCCTCTTCCCTGGCTG | 22 |
| | *miR6478* | CCGACCTTAGCTCAGTTGGTG | 21 |
| | *miR894* | CGTTTCACGTCGGGTTCACC | 20 |
| T2 | *miR156* | TGACAGAAGAGAGTGAGCAC | 20 |
| | | TTGACAGAAGAGAGTGAGCAC | 21 |
| | | TTGACAGAAGATAGAGAGCAC | 21 |
| | | TTGACAGAAGATAGAGGGCAC | 21 |
| | *miR157* | GCTCTCTATGCTTCTGTCATC | 21 |
| | *miR159* | CTTGGATTGAAGGGAGCTCTA | 21 |
| | | TTTGGATTGAAGGGAGCTCTG | 21 |
| | *miR162* | TCGATAAACCTCTGCATCCA | 20 |
| | *miR166* | TCGGACCAGGCTTCATTCCCG | 21 |
| | *miR167* | TGAAGCTGCCAGCATGATCTA | 21 |
| | | TGAAGCTGCCAGCATGATCTC | 21 |
| | | TGAAGCTGCCAGCATGATCTGC | 22 |
| | *miR168* | CCCGCCTTGCATCAACTGAAT | 21 |
| | | TCGCTTGGTGCAGGTCGGGA | 20 |
| | | TCGCTTGGTGCAGGTCGGGAA | 21 |
| | *miR169* | TAGCCAAAAATGACTTGCCTGC | 22 |
| | *miR319* | CTTGGACTGAAGGGAGCTCC | 20 |
| | | CTTGGACTGAAGGGAGCTCCC | 21 |
| | | TTGGACTGAAGGGAGCTCCCT | 21 |
| | *miR390* | AAGCTCAGGAGGGATAGCGCC | 21 |
| | *miR396* | TTCCACAGCTTTCTTGAACTA | 21 |
| | | TTCCACAGCTTTCTTGAACTG | 21 |
| | | TTCCACAGCTTTCTTGAACTT | 21 |
| | *miR398* | CGTGTTCTCAGGTCGCCCCTG | 21 |
| | | TGTGTTCCCAGGTCGCCCCTG | 21 |
| | | TGTGTTCTCAGGTCACCCCT | 20 |
| | | TGTGTTCTCAGGTCACCCCTT | 21 |
| | | TGTGTTCTCAGGTCGCCCCCG | 21 |
| | | TGTGTTCTCAGGTCGCCCCTG | 21 |
| | *miR399* | AGGGCTTCTCTCCATTGGCAGG | 22 |
| | *miR408* | ATGCACTGCCTCTTCCCTGGC | 21 |
| | | TGCACTGCCTCTTCCCTGGC | 20 |
| | | TGCACTGCCTCTTCCCTGGCT | 21 |
| | | TGCACTGCCTCTTCCCTGGCTG | 22 |
| | *miR6478* | CCGACCTTAGCTCAGTTGGTG | 21 |
| | *miR7130* | GTTTGGAATGTGCGAGATGTGTGC | 24 |
| | *miR894* | CGTTTCACGTCGGGTTCACC | 20 |

**Table S4:** Detail of the miRNA family related sequences with antagonic expression in response to identical stress conditions. The values represented correspon to the obtained by DeSeq analisys. The selected values correspponding to the most highly represented sequences are in gray. ID: Internal identification code.

| miRNA | sequence | stress | time | base mean | log2FC | P value | size | ID |
|---|---|---|---|---|---|---|---|---|
| *miR166* | TCTCGGACCAGGCTTCATTC | Cold | T1 | 36,067 | -1,054 | 0,02023 | 20 | SNCMel000025245 |
| | TTGGACCAGGCTTCATTCCCC | | | 11,866 | 1,404 | 0,02317 | 21 | SNCMel000030146 |
| | TCGGACCAGGCTTCATTCCT | Shot day | T1 | 60,878 | -1,047 | 0,01486 | 20 | SNCMel000011869 |
| | TCTCGGACCAGGCTTCATTC | | | 37,136 | -1,220 | 0,00532 | 20 | SNCMel000025245 |
| | TTGGACCAGGCTTCATTCCCC | | | 11,505 | 1,139 | 0,04458 | 21 | SNCMel000030146 |
| *miR396* | TTCCACAGCTTTCTTGAACTG | HSVd | T2 | 75,975 | -1,294 | 0,00192 | 21 | SNCMel000020454 |
| | TTCCACAGCTTTCTTGAACTT | | | 1770,236 | 1,275 | 0,00008 | 21 | SNCMel000003352 |

**Table S5:** Description and detailed information of targets for stress-responsive miRNAs identified en melon plants at T1 and T2. The GO terms were estimated in base to information for homologous transcripts in arabidopsis.

| miRNA | Diferential expression | Predicted Target | Accesion number | ID | GO | GO Annotation | Main GO Functions |
|---|---|---|---|---|---|---|---|
| *miR156, miR157* | T1 - T2 | Squamosa promoter-binding-like protein | METC038709 | SPL | GO:0006351 | transcription, DNA-templated | Transcription regulation Developmen |
| | | | | | GO:0006355 | regulation of transcription, DNA-templated | |
| | | | | | GO:0010358 | leaf shaping | |
| | | | | | GO:0045893 | positive regulation of transcription, DNA-templated | |
| | | | | | GO:0048510 | regulation of transition from vegetative to reproductive... | |
| *miR159* | T1 - T2 | BES1/BZR1 homolog protein 4 | METC032818 | BEH4 | GO:0006351 | transcription, DNA-templated | Transcription regulation Development |
| | | | | | GO:0006355 | regulation of transcription, DNA-templated | |
| | | | | | GO:0009742 | brassinosteroid mediated signaling pathway | |
| *miR160* | T1 | Auxin response factor 17 | METC021734 | ARF17 | GO:0006351 | transcription, DNA-templated | Transcription regulation Development |
| | | | | | GO:0006355 | regulation of transcription, DNA-templated | |
| | | | | | GO:0009555 | pollen development | |
| | | | | | GO:0009653 | anatomical structure morphogenesis | |
| | | | | | GO:0009725 | response to hormone | |
| | | | | | GO:0009734 | auxin-activated signaling pathway | |
| | | | | | GO:0048830 | adventitious root development | |
| *miR162* | T1 - T2 | Chorophyllase 2 | MU44640 | CLH2 | GO:0015996 | chlorophyll catabolic process | Photosynthesis metabolism |
| *miR166* | T1 - T2 | Homeobox-leucine zipper protein ATHB-14-like | XM_008441332.2 | ATHB14 | GO:0006351 | transcription, DNA-templated | Transcription regulation Developmen |
| | | | | | GO:0006355 | regulation of transcription, DNA-templated | |
| | | | | | GO:0009855 | determination of bilateral symmetry | |
| | | | | | GO:0009944 | polarity specification of adaxial/abaxial axis | |
| | | | | | GO:0009955 | adaxial/abaxial pattern specification | |
| | | | | | GO:0010014 | meristem initiation | |
| | | | | | GO:0030154 | cell differentiation | |
| *miR167* | T1 - T2 | Auxin response factor 6 | XM_008441364.2 | ARF6 | GO:0009908 | flower development | Transcription regulation Developmen |
| | | | | | GO:0006351 | transcription, DNA-templated | |
| | | | | | GO:0009725 | response to hormone | |
| | | | | | GO:0009734 | auxin-activated signaling pathway | |
| *miR168* | T1 - T2 | Argonaute 1 | XM_008440788.2 | AGO1 | GO:0006355 | regulation of transcription, DNA-templated | RNA silencing |
| | | | | | GO:0006952 | defense response | |
| | | | | | GO:0009616 | virus induced gene silencing | |
| | | | | | GO:0016441 | posttranscriptional gene silencing | |
| | | | | | GO:0035195 | gene silencing by miRNA | |
| | | | | | GO:0045087 | innate immune response | |
| | | | | | GO:0060145 | viral gene silencing in virus induced gene silencing | |
| *miR172* | T1 | AP2-like ethylene-responsive transcription factor | METC002019 | AP2 | GO:0006351 | transcription, DNA-templated | Transcription regulation Development |
| | | | | | GO:0006355 | regulation of transcription, DNA-templated | |
| | | | | | GO:0007275 | multicellular organism development | |
| | | | | | GO:0009873 | ethylene-activated signaling pathway | |
| *miR319* | T1 - T2 | Transcription factor TCP4 | METC043926 | TCP4 | GO:0006351 | transcription, DNA-templated | Transcription regulation Development |
| | | | | | GO:0006355 | regulation of transcription, DNA-templated | |
| | | | | | GO:0007275 | multicellular organism development | |
| | | | | | GO:0009793 | embryo development ending in seed dormancy | |
| *miR395* | T1 | ATP sulfurylase 1 | METC031431 | APS1 | GO:0000103 | sulfate assimilation | Metals metabolism |
| | | | | | GO:0001887 | selenium compound metabolic process | |
| | | | | | GO:0016310 | phosphorylation | |
| | | | | | GO:0046686 | response to cadmium ion | |
| *miR396* | T1 - T2 | Growth-regulating factor | METC031668 | GRF | GO:0006351 | transcription, DNA-templated | Transcription regulation Developmen Stress response |
| | | | | | GO:0006355 | regulation of transcription, DNA-templated | |
| | | | | | GO:0032502 | developmental process | |
| | | | | | GO:0048366 | leaf development | |
| *miR398* | T1 - T2 | Superoxide dismutase [Cu-Zn] | METC018907 | SOD | GO:0055114 | oxidation-reduction process | Response to oxidative stress |
| | | | | | GO:0019430 | removal of superoxide radicals | |
| | | | | | GO:0006979 | response to oxidative stress | |
| | | | | | GO:0034599 | cellular response to oxidative stress | |
| *miR408* | T1 - T2 | Basic blue protein-like | METC003548 | BABL | GO:0055114 | oxidation-reduction process | Oxidation-reduction process |
| | | | | | GO:0022900 | electron transport chain | |
| | | | | | GO:0048653 | anther development | |
| *miR6478* | T1 - T2 | Ferredoxin-like protein | MU49954 | FER | GO:0055114 | oxidation-reduction process | Oxidation-reduction process |
| | | | | | GO:0009643 | photosynthetic acclimation | |
| | | | | | GO:0009767 | photosynthetic electron transport chain | |
| *miR894* | T1 - T2 | Cysteine-rich and transmembrane domain-containing protein A-like | METC023591 | CYSTM | GO:0003006 | developmental process involved in reproduction | Development |
| *miR399* | T1 - T2 | Beta-glucosidase 11-like | METC013060 | BGlu11 | GO:0004553 | hydrolase activit ( O-glycosyl compounds) | Carbohydrates metabolism |
| | | | | | GO:0008422 | beta-glucosidase activity | |
| *miR169* | T2 | Nuclear transcription factor Y subunit A-1-like | METC025602 | NFYA1 | GO:0006351 | transcription, DNA-templated | Transcription regulation Developmen |
| | | | | | GO:0006355 | regulation of transcription, DNA-templated | |
| | | | | | GO:0009793 | embryo development ending in seed dormancy | |
| | | | | | GO:0010262 | somatic embryogenesis | |
| | | | | | GO:0048316 | seed development | |
| | | | | | GO:0048510 | regulation of transition from vegetative to reproductive... | |
| | | | | | GO:0055046 | microgametogenesis | |
| *miR7130* | T2 | COP9 signalosome complex subunit 6A | MU54233 | CSN6A | GO:0010387 | COP9 signalosome assembly | Photomorphogenesis |
| | | | | | GO:0007275 | multicellular organism development | |
| | | | | | GO:0030163 | protein catabolic process | |

**Table S6:** Details of the diversity of the melon miRNA family-related sequences during the exposition to stress conditions analyzed here.

| *miRNAs* | Stress responsive miRNA families | | | sRNA sequences | Stress responsive Family members | | |
|---|---|---|---|---|---|---|---|
| | T1 | T2 | T3 | | T1 | T2 | T3 |
| ***miR156*** | 1 | | | TGACAGAAGAGAGTGAGCAC | 1 | | |
| | | | | TGACAGAAGAGAGTGAGCACA | 2 | | |
| | | | | TGACAGAAGAGAGTGAGCACT | 3 | | |
| | | | | TTGACAGAAGAGAGTGAGCAC | 4 | | |
| | | | | TTGACAGAAGATAGAGAGCAC | 5 | | |
| ***miR157*** | 2 | | | GCTCTCTATACTTCTGTCACC | 6 | | |
| | | | | GCTCTCTATGCTTCTGTCATC | 7 | | |
| ***miR159*** | 3 | | | TTTGGACTGAAGGGAGCTCTA | 8 | | |
| | | | | TTTGGATTGAAGGGAGCTCTA | 9 | | |
| | | | | TTTGGATTGAAGGGAGCTCTT | 10 | | |
| ***miR160*** | 4 | | | TGCCTGGCTCCCTGTATGCC | 11 | | |
| ***miR162*** | 5 | | | TCGATAAACCTCTGCATCCAG | 12 | | |
| ***miR166*** | 6 | | | CGGACCAGGCTTCATTCCCC | 13 | | |
| | | | | TCGGACCAGGCTTCATTCCC | 14 | | |
| | | | | TCGGACCAGGCTTCATTCCT | 15 | | |
| | | | | TCGGACCAGGCTTCATTCCTC | 16 | | |
| | | | | TCTCGGACCAGGCTTCATTC | 17 | | |
| | | | | TCTCGGACCAGGCTTCATTCC | 18 | | |
| | | | | TCTCGGACCAGGCTTCATTCT | 19 | | |
| | | | | TTGGACCAGGCTTCATTCCCC | 20 | | |
| ***miR167*** | 7 | | | TGAAGCTGCCAGCATGATCT | 21 | | |
| | | | | TGAAGCTGCCAGCATGATCTGC | 22 | | |
| ***miR168*** | 8 | | | CCCGCCTTGCATCAACTGAAT | 23 | | |
| | | | | TCGCTTGGTGCAGGTCGGGA | 24 | | |
| | | | | TCGCTTGGTGCAGGTCGGGAA | 25 | | |
| ***miR172*** | 9 | | | AGAATCTTGATGATGCTGCAT | 26 | | |
| ***miR319*** | 10 | | | CTTGGACTGAAGGGAGCTCCC | 27 | | |
| | | | | TTGGACTGAAGGGAGCTCCC | 28 | | |
| | | | | TTGGACTGAAGGGAGCTCCCT | 29 | | |
| | | | | TTGGACTGAAGGGAGCTCCT | 30 | | |
| | | | | TTGGACTGAAGGGAGCTCCTTC | 31 | | |
| ***miR395*** | 11 | | | TGAAGTGTTTGGGGGAACTC | 32 | | |
| ***miR396*** | 12 | | | TTCCACAGCTTTCTTGAACT | 33 | | |
| | | | | TTCCACAGCTTTCTTGAACTG | 34 | | |
| | | | | TTCCACAGCTTTCTTGAACTT | 35 | | |
| | | | | TTCCACGGCTTTCTTGAACT | 36 | | |
| | | | | TTCCACGGCTTTCTTGAACTG | 37 | | |
| | | | | TTCCACGGCTTTCTTGAACTT | 38 | | |
| ***miR398*** | 13 | | | TGTGTTCCCAGGTCGCCCCTG | 39 | | |
| | | | | TGTGTTCTCAGGTCACCCCT | 40 | | |
| | | | | TGTGTTCTCAGGTCGCCCCTG | 41 | | |
| ***miR399*** | 14 | | | AGGGCTTCTCTCCATTGGCAGG | 42 | | |
| ***miR408*** | 15 | | | ATGCACTGCCTCTTCCCTGGC | 43 | | |
| | | | | TGCACTGCCTCTTCCCTGGC | 44 | | |
| | | | | TGCACTGCCTCTTCCCTGGCT | 45 | | |
| | | | | TGCACTGCCTCTTCCCTGGCTG | 46 | | |
| ***miR6478*** | 16 | | | CCGACCTTAGCTCAGTTGGTG | 47 | | |
| ***miR894*** | 17 | | | CGTTTCACGTCGGGTTCACC | 48 | | |
| ***miR156*** | | 1 | | TGACAGAAGAGAGTGAGCAC | | 1 | |
| | | | | TTGACAGAAGAGAGTGAGCAC | | 2 | |
| | | | | TTGACAGAAGATAGAGAGCAC | | 3 | |
| | | | | TTGACAGAAGATAGAGGGCAC | | 4 | |
| ***miR157*** | | 2 | | GCTCTCTATGCTTCTGTCATC | | 5 | |
| ***miR159*** | | 3 | | CTTGGATTGAAGGGAGCTCTA | | 6 | |
| | | | | TTTGGATTGAAGGGAGCTCTG | | 7 | |
| ***miR162*** | | 4 | | TCGATAAACCTCTGCATCCA | | 8 | |
| ***miR166*** | | 5 | | TCGGACCAGGCTTCATTCCCG | | 9 | |
| ***miR167*** | | 6 | | TGAAGCTGCCAGCATGATCTA | | 10 | |
| | | | | TGAAGCTGCCAGCATGATCTC | | 11 | |
| | | | | TGAAGCTGCCAGCATGATCTGC | | 12 | |
| ***miR168*** | | 7 | | CCCGCCTTGCATCAACTGAAT | | 13 | |
| | | | | TCGCTTGGTGCAGGTCGGGA | | 14 | |
| | | | | TCGCTTGGTGCAGGTCGGGAA | | 15 | |
| ***miR169*** | | 8 | | TAGCCAAAAATGACTTGCCTGC | | 16 | |
| ***miR319*** | | 9 | | CTTGGACTGAAGGGAGCTCC | | 17 | |
| | | | | CTTGGACTGAAGGGAGCTCCC | | 18 | |
| | | | | TTGGACTGAAGGGAGCTCCCT | | 19 | |
| ***miR390*** | | 10 | | AAGCTCAGGAGGGATAGCGCC | | 20 | |
| ***miR396*** | | 11 | | TTCCACAGCTTTCTTGAACTA | | 21 | |
| | | | | TTCCACAGCTTTCTTGAACTG | | 22 | |
| | | | | TTCCACAGCTTTCTTGAACTT | | 23 | |
| ***miR398*** | | 12 | | CGTGTTCTCAGGTCGCCCCTG | | 24 | |
| | | | | TGTGTTCCCAGGTCGCCCCTG | | 25 | |
| | | | | TGTGTTCTCAGGTCACCCCT | | 26 | |
| | | | | TGTGTTCTCAGGTCACCCCTT | | 27 | |
| | | | | TGTGTTCTCAGGTCGCCCCCG | | 28 | |
| | | | | TGTGTTCTCAGGTCGCCCCTG | | 29 | |
| ***miR399*** | | 13 | | AGGGCTTCTCTCCATTGGCAGG | | 30 | |
| ***miR408*** | | 14 | | ATGCACTGCCTCTTCCCTGGC | | 31 | |
| | | | | TGCACTGCCTCTTCCCTGGC | | 32 | |
| | | | | TGCACTGCCTCTTCCCTGGCT | | 33 | |
| | | | | TGCACTGCCTCTTCCCTGGCTG | | 34 | |
| ***miR6478*** | | 15 | | CCGACCTTAGCTCAGTTGGTG | | 35 | |
| ***miR7130*** | | 16 | | GTTTGGAATGTGCGAGATGTGTGC | | 36 | |
| ***miR894*** | | 17 | | CGTTTCACGTCGGGTTCACC | | 37 | |
| ***miR1515*** | | | 1 | TCATTTTTGCGTGCAATGATCC | | | 1 |
| ***miR156*** | | | 2 | CTGACAGAAGAGAGTGAGCAC | | | 2 |
| | | | | TGACAGAAGAGAGTGAGCAC | | | 3 |
| | | | | TGACAGAAGAGAGTGAGCACA | | | 4 |
| | | | | TGACAGAAGAGAGTGAGCACT | | | 5 |
| | | | | TGACAGAAGAGAGTGGGCAC | | | 6 |
| | | | | TGACAGAAGATAGAGAGCAC | | | 7 |
| | | | | TTGACAGAAGAGAGTGAGCAC | | | 8 |
| | | | | TTGACAGAAGATAGAGAGCAC | | | 9 |
| | | | | TTGACAGAAGATAGAGGGCAC | | | 10 |
| ***miR157*** | | | 3 | GCTCTCTATACTTCTGTCACC | | | 11 |
| | | | | GCTCTCTATGCTTCTGTCATC | | | 12 |
| ***miR159*** | | | 4 | TTTGGACTGAAGGGAGCTCTA | | | 13 |
| | | | | TTTGGATTGAAGGGAGCTCCT | | | 14 |
| | | | | TTTGGATTGAAGGGAGCTCTC | | | 15 |
| | | | | TTTGGATTGAAGGGAGCTCTG | | | 16 |
| | | | | TTTGGATTGAAGGGAGCTCTT | | | 17 |
| ***miR160*** | | | 5 | TGCCTGGCTCCCTGTATGCCA | | | 18 |
| ***miR162*** | | | 6 | TCGATAAGCCTCTGCATCCAG | | | 19 |
| ***miR164*** | | | 7 | TGGAGAAGCAGGGCACGTGCA | | | 20 |
| | | | | TGGAGAAGCAGGGCACGTGCT | | | 21 |
| ***miR165*** | | | 8 | TCGGACCAGGCTTCATCCCCC | | | 22 |
| ***miR166*** | | | 9 | CCGGACCAGGCTTCATTCCCC | | | 23 |
| | | | | CGGACCAGGCTTCATTCCCC | | | 24 |
| | | | | TCGAACCAGGCTTCATTCCCC | | | 25 |
| | | | | TCGGACCAGGCTTCATTCCC | | | 26 |
| | | | | TCGGACCAGGCTTCATTCCCC | | | 27 |
| | | | | TCGGACCAGGCTTCATTCCCCC | | | 28 |
| | | | | TCGGACCAGGCTTCATTCCCCT | | | 29 |
| | | | | TCGGACCAGGCTTCATTCCCG | | | 30 |
| | | | | TCGGACCAGGCTTCATTCCCT | | | 31 |
| | | | | TCGGACCAGGCTTCATTCCT | | | 32 |
| | | | | TCGGACCAGGCTTCATTCCTC | | | 33 |
| | | | | TCGGACCAGGCTTCATTCCTT | | | 34 |
| | | | | TCTCGGACCAGGCTTCATTC | | | 35 |
| | | | | TCTCGGACCAGGCTTCATTCC | | | 36 |
| | | | | TCTCGGACCAGGCTTCATTCT | | | 37 |
| | | | | TTGGACCAGGCTTCATTCCCC | | | 38 |
| ***miR167*** | | | 10 | TGAAGCTGCCAACATGATCTG | | | 39 |
| | | | | TGAAGCTGCCAGCATGATCT | | | 40 |
| | | | | TGAAGCTGCCAGCATGATCTA | | | 41 |
| | | | | TGAAGCTGCCAGCATGATCTC | | | 42 |
| | | | | TGAAGCTGCCAGCATGATCTG | | | 43 |
| | | | | TGAAGCTGCCAGCATGATCTGA | | | 44 |
| | | | | TGAAGCTGCCAGCATGATCTGC | | | 45 |
| | | | | TGAAGCTGCCAGCATGATCTT | | | 46 |
| | | | | TGAAGCTGCCAGCATGATCTTA | | | 47 |
| ***miR168*** | | | 11 | TCGCTTGGTGCAGGTCGGGA | | | 48 |
| | | | | TCGCTTGGTGCAGGTCGGGAA | | | 49 |
| | | | | TCGCTTGGTGCAGGTCGGGAAC | | | 50 |
| ***miR169*** | | | 12 | TAGCCAAAAATGACTTGCCTG | | | 51 |
| | | | | TAGCCAAAAATGACTTGCCTGC | | | 52 |
| | | | | TAGCCAAAGATGACTTGCCTG | | | 53 |
| | | | | TGAGCCAAGAATGACTTGCCGGC | | | 54 |
| ***miR171*** | | | 13 | TGATTGAGCCGCGCCAATATC | | | 55 |
| | | | | TGATTGAGCCGTGCCAATATC | | | 56 |
| | | | | TTGAGCCGCGTCAATATCTCT | | | 57 |
| | | | | TTGAGCCGTGCCAATATCACA | | | 58 |
| | | | | TTGAGCCGTGCCAATATCACG | | | 59 |
| ***miR172*** | | | 14 | AGAATCTTGATGATGCTGCAT | | | 60 |
| ***miR319*** | | | 15 | ATTGGACTGAAGGGAGCTCC | | | 61 |
| | | | | CTTGGACTGAAGGGAGCTCCC | | | 62 |
| | | | | CTTGGACTGAAGGGAGCTCCT | | | 63 |
| | | | | TTGGACTGAAGGGAGCTCCC | | | 64 |
| | | | | TTGGACTGAAGGGAGCTCCCA | | | 65 |
| | | | | TTGGACTGAAGGGAGCTCCCT | | | 66 |
| | | | | TTGGACTGAAGGGAGCTCCT | | | 67 |
| | | | | TTGGACTGAAGGGAGCTCCTTC | | | 68 |
| ***miR390*** | | | 16 | AAGCTCAGGAGGGATAGCGCC | | | 69 |
| ***miR393*** | | | 17 | ATCATGCTATCTCTTTGGATT | | | 70 |
| | | | | TCCAAAGGGATCGCATTGATC | | | 71 |
| ***miR394*** | | | 18 | TTGGCATTCTGTCCACCTCC | | | 72 |
| ***miR395*** | | | 19 | CTGAAGTGTTTGGGGGAACTC | | | 73 |
| | | | | TGAAGTGTTTGGGGGAACTCT | | | 74 |
| ***miR396*** | | | 20 | TTCCACAGCTTTCTTGAACGT | | | 75 |
| | | | | TTCCACAGCTTTCTTGAACT | | | 76 |
| | | | | TTCCACAGCTTTCTTGAACTA | | | 77 |
| | | | | TTCCACAGCTTTCTTGAACTG | | | 78 |
| | | | | TTCCACAGCTTTCTTGAACTGT | | | 79 |
| | | | | TTCCACAGCTTTCTTGAACTT | | | 80 |
| | | | | TTCCACGGCTTTCTTGAACTT | | | 81 |
| ***miR397*** | | | 21 | ATTGAGTGCAGCGTTGATGT | | | 82 |
| | | | | TCATTGAGTGCAGCGTTGATG | | | 83 |
| ***miR398*** | | | 22 | CGTGTTCTCAGGTCGCCCCTG | | | 84 |
| | | | | TATGTTCTCAGGTCGCCCCTG | | | 85 |
| | | | | TGTGTTCCCAGGTCGCCCCTG | | | 86 |
| | | | | TGTGTTCTCAGGTCACCCCT | | | 87 |
| | | | | TGTGTTCTCAGGTCACCCCTG | | | 88 |
| | | | | TGTGTTCTCAGGTCGCCCCCG | | | 89 |
| | | | | TGTGTTCTCAGGTCGCCCCTG | | | 90 |
| | | | | TTGTGTTCTCAGGTCACCCCT | | | 91 |
| ***miR408*** | | | 23 | ATGCACTGCCTCTTCCCTGGC | | | 92 |
| | | | | TGCACTGCCTCTTCCCTGGC | | | 93 |
| | | | | TGCACTGCCTCTTCCCTGGCT | | | 94 |
| | | | | TGCACTGCCTCTTCCCTGGCTG | | | 95 |
| ***miR6478*** | | | 24 | CCGACCTTAGCTCAGTTGGTG | | | 96 |

**Table S7:** Functional relation between narrow- and broad- response range stress-responsive miRNAs identified en melon plants during this assay. Main functional GO terms are grouped in colours according their predicted functions in melon.

| *Narrow response-range* | | |
|---|---|---|
| miRNAs | targets | main GO term |
| *miR394* | EG45 | Stress response |
| *miR399* | BGlu11 | Carbohydrates metabolism |
| *miR395* | APS1 | Metals metabolism |
| *miR162* | CLH2 | Photosysnthesis |
| *miR1515* | GLRY2 | Redox |
| *miR894* | CYSTM | Reproduction |
| *miR7130* | CSN6A | Photomorphogenesis |
| *miR160* | ARF17 | TF-Development |
| *miR164* | NAC100 | TF-Development |
| *miR165* | ATHB14 | TF-Development |
| *miR390* | TAS3 | RNA silencing |

| *Broad response-range* | | |
|---|---|---|
| miRNAs | targets | main GO term |
| *miR396* | GFR | TF - Development |
| *miR156* | SPL | TF - Development |
| *miR166* | ATBH14 | TF - Development |
| *miR167* | ARF6 | TF - Development |
| *miR159* | BEH4 | TF - Development |
| *miR319* | TCP | TF - Development |
| *miR393* | F-Box | Development |
| *miR398* | SOD | Cu homeostasis |
| *miR408* | BABL | Cu homeostasis |
| *miR6478* | FER | REDOX |

**Table S8:** Values matrix for clusters network building. ***Link***: number of stress conditions in which both miRNAs share cluster (value for links thickness), ***W. Link***: =Link/# stress conditions (value for links length), ***Node***: sum of link values for each source miRNA (node size). **C**: cold, **D**: drought, **Sal** : salinity, **SD**: short day, **Mon**: *Monosporascus*, **Agro**: *Agrobacterium*, **HSVd**: Hop stunt viroid.

| Source | Target | Link | W. Link | Node | Detail of the stress conditions |
|---|---|---|---|---|---|
| *miR398* | *miR408* | 7 | 1 | 10 | C;D;Sal;SD;Mon;Agro;HSVd |
| *miR157* | *miR6478* | 3 | 0.43 | 20 | Mon;Agro;HSVd |
| *miR6478* | *miR156* | 3 | 0.43 | 17 | D;Mon;Agro |
| *miR319* | *miR396* | 3 | 0.43 | 23 | D;SD;HSVd |
| *miR166* | *miR168* | 3 | 0.43 | 22 | C;Mon;HSVd |
| *miR166* | *miR162* | 2 | 0.29 | 22 | Agro;HSVd |
| *miR157* | *miR319* | 2 | 0.29 | 20 | C;Mon |
| *miR157* | *miR393* | 2 | 0.29 | 20 | C;Agro |
| *miR157* | *miR167* | 2 | 0.29 | 20 | Sal;SD |
| *miR157* | *miR156* | 2 | 0.29 | 20 | Mon;Agro |
| *miR157* | *miR168* | 2 | 0.29 | 20 | D;Agro |
| *miR319* | *miR393* | 2 | 0.29 | 23 | C;HSVd |
| *miR319* | *miR160* | 2 | 0.29 | 23 | Sal;HSVd |
| *miR319* | *miR167* | 2 | 0.29 | 23 | Agro;HSVd |
| *miR319* | *miR156* | 2 | 0.29 | 23 | Mon;HSVd |
| *miR396* | *miR894* | 2 | 0.29 | 23 | C;Mon |
| *miR396* | *miR156* | 2 | 0.29 | 23 | Agro;HSVd |
| *miR396* | *miR172* | 2 | 0.29 | 23 | Agro;HSVd |
| *miR166* | *miR395* | 2 | 0.29 | 22 | Sal;Mon |
| *miR166* | *miR390* | 2 | 0.29 | 22 | Agro;HSVd |
| *miR6478* | *miR162* | 1 | 0.14 | 17 | C |
| *miR319* | *miR162* | 1 | 0.14 | 23 | Agro |
| *miR396* | *miR171* | 1 | 0.14 | 23 | HSVd |
| *miR6478* | *miR319* | 1 | 0.14 | 17 | Mon |
| *miR6478* | *miR393* | 1 | 0.14 | 17 | Agro |
| *miR6478* | *miR396* | 1 | 0.14 | 17 | Agro |
| *miR6478* | *miR159* | 1 | 0.14 | 17 | Sal |
| *miR6478* | *miR168* | 1 | 0.14 | 17 | Agro |
| *miR6478* | *miR172* | 1 | 0.14 | 17 | Agro |
| *miR398* | *miR6478* | 1 | 0.14 | 10 | SD |
| *miR408* | *miR6478* | 1 | 0.14 | 10 | SD |
| *miR157* | *miR169* | 1 | 0.14 | 20 | C |
| *miR157* | *miR396* | 1 | 0.14 | 20 | Agro |
| *miR157* | *miR159* | 1 | 0.14 | 20 | SD |
| *miR157* | *miR172* | 1 | 0.14 | 20 | Agro |
| *miR157* | *miR398* | 1 | 0.14 | 20 | Sal |
| *miR157* | *miR408* | 1 | 0.14 | 20 | Sal |
| *miR319* | *miR164* | 1 | 0.14 | 23 | HSVd |
| *miR319* | *miR166* | 1 | 0.14 | 23 | Agro |
| *miR319* | *miR168* | 1 | 0.14 | 23 | Sal |
| *miR319* | *miR172* | 1 | 0.14 | 23 | HSVd |
| *miR319* | *miR390* | 1 | 0.14 | 23 | Agro |
| *miR396* | *miR168* | 1 | 0.14 | 23 | Agro |
| *miR166* | *miR172* | 1 | 0.14 | 22 | C |
| *miR156* | *miR172* | 3 | 0.43 | 30 | C;Agro;HSVd |
| *miR156* | *miR171* | 2 | 0.29 | 30 | Sal;HSVd |
| *miR156* | *miR165* | 2 | 0.29 | 30 | C;Sal |
| *miR156* | *miR166* | 2 | 0.29 | 30 | C;Sal |
| *miR156* | *miR168* | 2 | 0.29 | 30 | C;Agro |
| *miR168* | *miR172* | 2 | 0.29 | 21 | C;Agro |
| *miR168* | *miR162* | 1 | 0.14 | 21 | HSVd |
| *miR156* | *miR159* | 1 | 0.14 | 30 | C |
| *miR156* | *miR395* | 1 | 0.14 | 30 | Sal |
| *miR168* | *miR395* | 1 | 0.14 | 21 | Mon |
| *miR168* | *miR390* | 1 | 0.14 | 21 | HSVd |
| *miR393* | *miR156* | 3 | 0.43 | 24 | Sal;A;HSVd |
| *miR393* | *miR171* | 2 | 0.29 | 24 | Sal;HSVd |
| *miR393* | *miR396* | 2 | 0.29 | 24 | Agro;HSVd |
| *miR393* | *miR166* | 2 | 0.29 | 24 | Sal;SD |
| *miR393* | *miR172* | 2 | 0.29 | 24 | Agro;HSVd |
| *miR167* | *miR396* | 2 | 0.29 | 24 | C;HSVd |
| *miR167* | *miR162* | 1 | 0.14 | 24 | Agro |
| *miR393* | *miR160* | 1 | 0.14 | 24 | HSVd |
| *miR393* | *miR164* | 1 | 0.14 | 24 | HSVd |
| *miR393* | *miR167* | 1 | 0.14 | 24 | HSVd |
| *miR393* | *miR165* | 1 | 0.14 | 24 | Sal |
| *miR393* | *miR168* | 1 | 0.14 | 24 | Agro |
| *miR393* | *miR395* | 1 | 0.14 | 24 | Sal |
| *miR167* | *miR394* | 1 | 0.14 | 24 | C |
| *miR167* | *miR894* | 1 | 0.14 | 24 | C |
| *miR167* | *miR156* | 1 | 0.14 | 24 | HSVd |
| *miR167* | *miR159* | 1 | 0.14 | 24 | SD |
| *miR167* | *miR166* | 1 | 0.14 | 24 | Agro |
| *miR167* | *miR172* | 1 | 0.14 | 24 | HSVd |
| *miR167* | *miR398* | 1 | 0.14 | 24 | Sal |
| *miR167* | *miR408* | 1 | 0.14 | 24 | Sal |
| *miR167* | *miR399* | 1 | 0.14 | 24 | SD |
| *miR167* | *miR390* | 1 | 0.14 | 24 | Agro |
| *miR169* | *miR319* | 2 | 0.29 | 12 | C;HSVd |
| *miR169* | *miR393* | 2 | 0.29 | 12 | C;HSVd |
| *miR159* | *miR166* | 2 | 0.29 | 15 | C;HSVd |
| *miR159* | *miR168* | 2 | 0.29 | 15 | C;HSVd |
| *miR159* | *miR162* | 1 | 0.14 | 15 | HSVd |
| *miR169* | *miR171* | 1 | 0.14 | 12 | HSVd |
| *miR169* | *miR160* | 1 | 0.14 | 12 | HSVd |
| *miR169* | *miR164* | 1 | 0.14 | 12 | HSVd |
| *miR169* | *miR167* | 1 | 0.14 | 12 | HSVd |
| *miR169* | *miR396* | 1 | 0.14 | 12 | HSVd |
| *miR169* | *miR156* | 1 | 0.14 | 12 | HSVd |
| *miR169* | *miR172* | 1 | 0.14 | 12 | HSVd |
| *miR159* | *miR165* | 1 | 0.14 | 15 | C |
| *miR159* | *miR172* | 1 | 0.14 | 15 | C |
| *miR159* | *miR7130* | 1 | 0.14 | 15 | Sal |
| *miR159* | *miR399* | 1 | 0.14 | 15 | SD |
| *miR159* | *miR390* | 1 | 0.14 | 15 | HSVd |
| *miR160* | *miR164* | 2 | 0.29 | 16 | C;HSVd |
| *miR160* | *miR167* | 2 | 0.29 | 16 | C;HSVd |
| *miR160* | *miR396* | 2 | 0.29 | 16 | C;HSVd |
| *miR162* | *miR171* | 1 | 0.14 | 10 | C |
| *miR171* | *miR6478* | 1 | 0.14 | 16 | C |
| *miR171* | *miR319* | 1 | 0.14 | 16 | HSVd |
| *miR160* | *miR171* | 1 | 0.14 | 16 | HSVd |
| *miR171* | *miR167* | 1 | 0.14 | 16 | HSVd |
| *miR171* | *miR166* | 1 | 0.14 | 16 | Sal |
| *miR172* | *miR171* | 1 | 0.14 | 21 | HSVd |
| *miR160* | *miR156* | 1 | 0.14 | 16 | HSVd |
| *miR160* | *miR168* | 1 | 0.14 | 16 | Sal |
| *miR160* | *miR172* | 1 | 0.14 | 16 | HSVd |
| *miR390* | *miR162* | 2 | 0.29 | 8 | Agro;HSVd |
| *miR164* | *miR167* | 2 | 0.29 | 14 | C;HSVd |
| *miR164* | *miR396* | 2 | 0.29 | 14 | C;HSVd |
| *miR165* | *miR166* | 2 | 0.29 | 11 | C;Sal |
| *miR164* | *miR171* | 1 | 0.14 | 14 | HSVd |
| *miR165* | *miR171* | 1 | 0.14 | 11 | Sal |
| *miR395* | *miR171* | 1 | 0.14 | 7 | Sal |
| *miR894* | *miR160* | 1 | 0.14 | 6 | C |
| *miR164* | *miR394* | 1 | 0.14 | 14 | C |
| *miR164* | *miR894* | 1 | 0.14 | 14 | C |
| *miR164* | *miR156* | 1 | 0.14 | 14 | HSVd |
| *miR164* | *miR172* | 1 | 0.14 | 14 | HSVd |
| *miR165* | *miR168* | 1 | 0.14 | 11 | C |
| *miR165* | *miR172* | 1 | 0.14 | 11 | C |
| *miR165* | *miR395* | 1 | 0.14 | 11 | Sal |
| *miR7130* | *miR6478* | 1 | 0.14 | 2 | Sal |
| *miR399* | *miR157* | 1 | 0.14 | 3 | SD |
| *miR394* | *miR160* | 1 | 0.14 | 5 | C |
| *miR394* | *miR396* | 1 | 0.14 | 5 | C |
| *miR394* | *miR894* | 1 | 0.14 | 5 | C |
| *miR1515* | *miR156* | 1 | 0.14 | 6 | C |
| *miR1515* | *miR159* | 1 | 0.14 | 6 | C |
| *miR1515* | *miR165* | 1 | 0.14 | 6 | C |
| *miR1515* | *miR166* | 1 | 0.14 | 6 | C |
| *miR1515* | *miR168* | 1 | 0.14 | 6 | C |
| *miR1515* | *miR172* | 1 | 0.14 | 6 | C |

**Table S9:** Description of the sRNAs libraries used to infer stress responsive sRNAs in soybean and rice plants. The diverse stress treatments are also detailed.

| Specie | BioProject | Accesion | Treatment | Code - Used | References |
|---|---|---|---|---|---|
| ***Oriza sativa*** | PRJNA146417 | GSM816687 | Control | CONTROL | JEONG, et al. (2011). The Plant Cell 23: 4185–4207. |
| | | GSM816694 | Cold | C | |
| | | GSM816690 | Drought | D | |
| | | GSM816696 | Heat | H | |
| | | GSM816692 | Salt | SA | |
| | Not available | Sequencing data available in the website indicated in the article. | Mock(RDV) | CONTROL | DU, et al. (2011) PLoS Pathogens 7: 8. |
| | | | RDV – *Rice dwarf virus* | RDV | |
| | | | Mock(RSV) | CONTROL | |
| | | | RSV – *Rice stripe virus* | RSV | |
| | PRJNA277442 | SRR1849771 | Control_120_min | CONTROL | BALDRICH, et al. (2015) RNA Biology 12: 847-863. |
| | | SRR1849774 | Elicitor_120_min | EL | |
| | PRJNA252424 | GSM1409674 | CK24h | CONTROL | ZHAO, et al. (2015) J. of Genetics and Genomics 4: 625-637. |
| | | GSM1409678 | Xoo24h | X | |
| ***Glicine max*** | PRJNA80127 | SRR330122 | Mock | CONTROL | LI, et al. (2011). BMC Plant Biology 11: 1 - 170. |
| | | SRR330121 | Alkanility | ALK | |
| | | SRR330123 | Drought | DD | |
| | | SRR330124 | Salinity | SA | |
| | PRJNA167179 | SRR499410 | Al-free | CONTROL | ZENG, et al., (2012). BMC Plant Biology 12: 182. |
| | | SRR499409 | Al-treated | AL | |
| | PRJNA244779 | SRR1240697 | Mock-inoculated (Williams) | CONTROL | ZHAO, et al. (2015). The Plant Genome 8: 1. |
| | | SRR1240707 | P. sojae inoculated (Williams) | P | |
| | PRJNA253492 | SRR1451615 | Well watered (LD003309) | CONTROL | ARIKIT, et al. (2014) The Plant Cell 86: 4584-4601. |
| | | SRR1451621 | Drought stressed (LD003309) | D | |
| | | SRR1451637 | Water 30 minutes | CONTROL | |
| | | SRR1451639 | Flagellin treated 30 minutes | FLAG | |

**Table S10:** Details of the conectivity values (CV) determined for the miRNAs categorized as Broad, Intermediate and Narrow response range in melon, soybean and rice plants.

| miRNAs | CV | Response Range | | |
|---|---|---|---|---|
| -->*mir156* | 2 | | | |
| -->*mir166* | 2 | | | |
| -->*mir167* | 2 | | | |
| -->*mir390* | 2 | | | |
| -->*mir408* | 2 | | | |
| -->*mir482* | 0 | | | |
| -->*mir160* | 2 | | | |
| -->*mir164* | 2 | | | |
| -->*mir169* | 2 | | Mean CV | SE |
| -->*mir1863* | 0 | | 1,3 | 0,21885 |
| -->*mir394* | 2 | ***Broad*** | | |
| -->*mir396* | 2 | | | |
| -->*mir444* | 0 | | | |
| -->*mir5150* | 0 | | | |
| -->*mir812* | 0 | | | |
| -->*mir157* | 0 | | | |
| -->*mir319* | 2 | | | |
| -->*mir393* | 2 | | | |
| -->*mir398* | 2 | | | |
| -->*mir6478* | 0 | | | |
| -->*mir171* | 2 | | | |
| -->*mir1509* | 0 | | | |
| -->*mir2118* | 1 | | | |
| -->*mir1512* | 0 | | | |
| -->*mir397* | 2 | | | |
| -->*mir159* | 2 | | | |
| -->*mir5225* | 0 | | | |
| -->*mir172* | 2 | | | |
| -->*mir5373* | 0 | | | |
| -->*mir403* | 0 | | | |
| -->*mir3522* | 0 | | | |
| -->*mir1507* | 0 | | | |
| -->*mir5767* | 0 | | | |
| -->*mir2119* | 0 | | | |
| -->*mir399* | 2 | | | |
| -->*mir1862* | 0 | | | |
| -->*mir1882* | 0 | | | |
| -->*mir1423* | 0 | | | |
| -->*mir2867* | 0 | | Mean CV | SE |
| -->*mir1425* | 0 | ***Intermediate*** | 0,3 | 0,1086 |
| -->*mir2863* | 0 | | | |
| -->*mir5807* | 0 | | | |
| -->*mir5802* | 0 | | | |
| -->*mir1871* | 0 | | | |
| -->*mir1861* | 0 | | | |
| -->*mir1856* | 0 | | | |
| -->*mir1870* | 0 | | | |
| -->*mir827* | 0 | | | |
| -->*mir5795* | 0 | | | |
| -->*mir3979* | 0 | | | |
| -->*mir2121* | 0 | | | |
| -->*mir1881* | 0 | | | |
| -->*mir535* | 0 | | | |
| -->*mir1879* | 0 | | | |
| -->*mir1432* | 0 | | | |
| -->*mir168* | 1 | | | |
| -->*mir1876* | 0 | | | |
| -->*mir820* | 0 | | | |
| -->*mir6254* | 0 | | | |
| -->*mir1850* | 0 | | | |
| -->*mir5671* | 0 | | | |
| -->*mir1515* | 1 | | | |
| -->*mir4415* | 0 | | | |
| -->*mir5668* | 0 | | | |
| -->*mir1510* | 0 | | | |
| -->*mir5781* | 0 | | | |
| -->*mir2109* | 0 | | | |
| -->*mir6300* | 0 | | | |
| -->*mir1508* | 0 | | | |
| -->*mir162* | 2 | | | |
| -->*mir5376* | 0 | | | |
| -->*mir1513* | 0 | | | |
| -->*mir5786* | 0 | | | |
| -->*mir5770* | 0 | | | |
| -->*mir5372* | 0 | | | |
| -->*mir5761* | 0 | | | |
| -->*mir5038* | 0 | | | |
| -->*mir4397* | 0 | | | |
| -->*mir5037* | 0 | | | |
| -->*mir862* | 0 | | | |
| -->*mir2606* | 0 | | | |
| -->*mir1520* | 0 | | | |
| -->*mir5672* | 0 | | | |
| -->*mir9728* | 0 | | | |
| -->*mir4416* | 0 | | | |
| -->*mir2111* | 0 | | | |
| -->*mir4345* | 0 | | | |
| -->*mir4413* | 0 | | | |
| -->*mir5674* | 0 | | | |
| -->*mir5782* | 0 | | | |
| -->*mir5785* | 0 | | | |
| -->*mir5769* | 0 | | | |
| -->*mir4387* | 0 | | | |
| -->*mir4412* | 0 | | | |
| -->*mir5677* | 0 | | | |
| -->*mir5559* | 0 | | | |
| -->*mir4996* | 0 | | | |
| -->*mir9726* | 0 | | | |
| -->*mir5670* | 0 | | | |
| -->*mir1535* | 0 | | | |
| -->*mir9749* | 0 | | Mean CV | SE |
| -->*mir5374* | 0 | ***Narrow*** | 0,0366 | 333,727 |
| -->*mir2873* | 0 | | | |
| -->*mir5143* | 0 | | | |
| -->*mir6251* | 0 | | | |
| -->*mir528* | 0 | | | |
| -->*mir5813* | 0 | | | |
| -->*mir5814* | 0 | | | |
| -->*mir5801* | 0 | | | |
| -->*mir529* | 0 | | | |
| -->*mir1860* | 0 | | | |
| -->*mir5508* | 0 | | | |
| -->*mir1873* | 0 | | | |
| -->*mir1868* | 0 | | | |
| -->*mir1864* | 0 | | | |
| -->*mir1859* | 0 | | | |
| -->*mir5811* | 0 | | | |
| -->*mir2871* | 0 | | | |
| -->*mir1875* | 0 | | | |
| -->*mir2880* | 0 | | | |
| -->*mir810* | 0 | | | |
| -->*mir2877* | 0 | | | |
| -->*mir531* | 0 | | | |
| -->*mir5794* | 0 | | | |
| -->*mir1427* | 0 | | | |
| -->*mir1320* | 0 | | | |
| -->*mir1430* | 0 | | | |
| -->*mir439* | 0 | | | |
| -->*mir435* | 0 | | | |
| -->*mir5144* | 0 | | | |
| -->*mir1878* | 0 | | | |
| -->*mir5512* | 0 | | | |
| -->*mir5788* | 0 | | | |
| -->*mir1865* | 0 | | | |
| -->*mir1883* | 0 | | | |
| -->*mir5083* | 0 | | | |
| -->*mir530* | 0 | | | |
| -->*mir2878* | 0 | | | |
| -->*mir165* | 0 | | | |
| -->*mir395* | 0 | | | |
| -->*mir894* | 0 | | | |
| -->*mir7130* | 0 | | | |

**Table S11**: Detail of the oligos used in this work.

Oligos for targets amplification by qRT-PCR

| miRNA | Target | FORWARD | REVERSE |
|---|---|---|---|
| *miR156* | SPL9 | 5'- GGC AGA GCA ACT CTG AGA AC -3' | 5'- GAC AGT CGT CGT TCG GTT TC -3' |
| *miR166* | ATHB14 | 5'- AAC TGC TGT CGA CTG GGT CC -3' | 5'- CTG CAA CCT TTG TCG GCT CTA G -3' |
| *miR169* | NFY | 5'- AAT ATG GGC TGC CTC TCT TGG G -3' | 5'- AAC TAG AAC TAA GGT CCA AAT GGC -3' |
| *miR172* | AP2 | 5'- ATT CCG AGG GGT AGA AGC AG -3' | 5'- AAT TGG CCC ATT CGA GCT TC -3' |
| *miR319* | TCP2 | 5'- TCA ACA TTT CTC GTT CGT CCC -3' | 5'- ATG CAG CTC CGA TGA AAA AGG -3' |
| *miR396* | GRF9 | 5'- CGG TAA CGA CTC CGC TCA C -3' | 5'- GAT TGT GGT GTC GGT AAC GG -3' |
| *miR398* | SOD | 5'- CGT TCC CAT TCA ATC TCA CCT GC -3' | 5'- AAC AGT GGT TGG ACC ATC TCC -3' |
| | CUP | 5'-CTCACAACGTCGGCGATTC-3' | 5'-CCGGTGGTGAAATTGAAAAAGAG-3' |
| *miR408* | BBL | 5'- TAC GCC TCC TGC AAA ACT CC -3' | 5'- AGA TCA GAG AGC ATT GAC GGC -3' |

Oligos for miRNA amplification by stem loop qRT-PCR

| | | |
|---|---|---|
| *miR156* | miRNA sequencence | **TTGACAGAAGAGAGTGAGCAC** |
| | RT-miR156 | GTTGGCTCTGGTGCAGGGTCCGAGGTATTCGCACCAGAGCCAACgtgctc |
| | miR156-F | gcgcaccTTGACAGAAGAGAG |
| *miR168* | miRNA sequencence | **TCGCTTGGTGCAGGTCGGGAA** |
| | RT-miR168 | GTTGGCTCTGGTGCAGGGTCCGAGGTATTCGCACCAGAGCCAACttcccg |
| | miR168-F | gggtatTCGCTTGGTGCAGG |
| *miR169* | miRNA sequencence | **TAGCCAAAGATGACTTGCCTG** |
| | RT-miR169 | GTTGGCTCTGGTGCAGGGTCCGAGGTATTCGCACCAGAGCCAACcaggca |
| | miR169-F | gcgcaccTAGCCAAAGATGAC |
| *miR396* | miRNA sequencence | **GTTCAATAAAGCTGTGGGAAG** |
| | RT-mir396a | GTTGGCTCTGGTGCAGGGTCCGAGGTATTCGCACCAGAGCCAACcttccc |
| | mir396a-F | acggcgcaGTTCAATAAAGCTG |
| *miR398* | miRNA sequencence | **TGTGTTCTCAGGTCGCCCCTG** |
| | RT-mir398 | GTTGGCTCTGGTGCAGGGTCCGAGGTATTCGCACCAGAGCCAACcagggg |
| | mir398-F | gcgcaTGTGTTCTCAGGTCG |
| *miR408* | miRNA sequencence | **ATGCACTGCCTCTTCCCTGGC** |
| | RT-miR408 | GTTGGCTCTGGTGCAGGGTCCGAGGTATTCGCACCAGAGCCAACgccagg |
| | miR408-F | ccgcaATGCACTGCCTCTTC |
| *miR166* | miRNA sequencence | TCGGACCAGGCTTCATTCCTC |
| | RT-miR166 | GTTGGCTCTGGTGCAGGGTCCGAGGTATTCGCACCAGAGCCAACgaggaa |
| | miR166-F | |
| *miR399* | miRNA sequencence | AGGGCTTCTCTCCATTGGCAGG |
| | RT-miR399 | GTTGGCTCTGGTGCAGGGTCCGAGGTATTCGCACCAGAGCCAACcctgcc |
| | miR399-F | |
| *miR7130* | miRNA sequencence | GTTTGGAATGTGCGAGATGTGTGC |
| | RT-miR7130 | GTTGGCTCTGGTGCAGGGTCCGAGGTATTCGCACCAGAGCCAACgcacac |
| | miR7130-F | |

Oligos for degradome assay

| Oligo ID | Sequence |
|---|---|
| **PARE 5' Adaptor** | 5'-GACCACCGACAGGUUCAGAGUUCUACAGUCCGAC-3' |
| **dT primer** | 5'-ATTCTAGAGGCCGAGGCGGCCGACATG-d(T)30-3' |
| **RP1S** | 5'-GACCACCGACAGGTTCAGAGTTC-3' |
| **GR3'** | 5'-ATTCTAGAGGCCGAGGCGGCCGACATG-3' |
| **dsDNA-top** | 5'-TGGAATTCTCGGGTGCCAAGGAACTCCAGTCAC-3' |
| **dsDNA_bottom** | 5'-GTGACTGGAGTTCCTTGGCACCCGAGAATTCCANN-3' |
| **RP1M** | 5'-AATGATACGGCGACCACCGACAGGTTCAGAG-3' |
| **RPIndex** | 5'-CAAGCAGAAGACGGCATACGAGATxxxxxxGTGACTGGAGTTCCTTGGCAC-3' |

**Table S12:** Matrix of presence/absence of stress-responsive miRNAs in melon plants at T1 and T2. The values "1" and "0" represent respectively if whether or not a miRNA is reactive (with both either increased or decreased expression) to a specific stress condition. 1: reactive, 0: non reactive.

**T1**

| miRNA | cold | drought | salinity | short day | Monosp. | HSVd | Agro. | Total |
|---|---|---|---|---|---|---|---|---|
| *miR156* | 1 | 0 | 1 | 1 | 1 | 1 | 1 | 6 |
| *miR157* | 0 | 0 | 1 | 1 | 1 | 1 | 1 | 5 |
| *miR159* | 1 | 0 | 0 | 0 | 0 | 1 | 0 | 2 |
| *miR160* | 0 | 0 | 1 | 0 | 0 | 0 | 0 | 1 |
| *miR162* | 0 | 0 | 0 | 0 | 0 | 1 | 1 | 2 |
| *miR166* | 1 | 1 | 1 | 1 | 1 | 1 | 1 | 7 |
| *miR167* | 0 | 0 | 1 | 1 | 0 | 1 | 1 | 4 |
| *miR168* | 1 | 0 | 1 | 0 | 0 | 1 | 0 | 3 |
| *miR172* | 1 | 0 | 0 | 0 | 0 | 0 | 0 | 1 |
| *miR319* | 1 | 1 | 1 | 1 | 0 | 1 | 1 | 6 |
| *miR395* | 0 | 0 | 0 | 0 | 1 | 0 | 0 | 1 |
| *miR396* | 1 | 0 | 1 | 1 | 0 | 1 | 1 | 5 |
| *miR398* | 1 | 1 | 1 | 1 | 0 | 1 | 0 | 5 |
| *miR399* | 1 | 0 | 0 | 0 | 0 | 0 | 0 | 1 |
| *miR408* | 1 | 0 | 1 | 1 | 0 | 0 | 0 | 3 |
| *miR6478* | 0 | 1 | 1 | 1 | 0 | 1 | 0 | 4 |
| *miR894* | 0 | 0 | 0 | 0 | 1 | 1 | 0 | 2 |

**T2**

| miRNA | cold | drought | salinity | short day | Monosp. | HSVd | Agro. | Total |
|---|---|---|---|---|---|---|---|---|
| *miR156* | 1 | 0 | 0 | 0 | 1 | 1 | 1 | 4 |
| *miR157* | 0 | 0 | 0 | 0 | 0 | 1 | 1 | 2 |
| *miR159* | 1 | 0 | 1 | 1 | 0 | 1 | 0 | 4 |
| *miR162* | 0 | 0 | 0 | 0 | 0 | 0 | 1 | 1 |
| *miR166* | 0 | 0 | 0 | 1 | 0 | 1 | 1 | 3 |
| *miR167* | 1 | 0 | 0 | 0 | 0 | 1 | 1 | 3 |
| *miR168* | 1 | 0 | 0 | 0 | 1 | 1 | 1 | 4 |
| *miR169* | 0 | 0 | 1 | 0 | 0 | 0 | 0 | 1 |
| *miR319* | 1 | 0 | 0 | 0 | 0 | 0 | 1 | 2 |
| *miR390* | 0 | 0 | 0 | 0 | 0 | 1 | 1 | 2 |
| *miR396* | 1 | 0 | 1 | 1 | 1 | 1 | 1 | 6 |
| *miR398* | 1 | 1 | 0 | 0 | 1 | 1 | 1 | 5 |
| *miR399* | 1 | 0 | 0 | 1 | 0 | 0 | 0 | 2 |
| *miR408* | 1 | 1 | 0 | 0 | 1 | 1 | 1 | 5 |
| *miR6478* | 1 | 0 | 1 | 1 | 0 | 1 | 1 | 5 |
| *miR7130* | 0 | 0 | 1 | 0 | 0 | 0 | 0 | 1 |
| *miR894* | 1 | 0 | 0 | 0 | 0 | 0 | 0 | 1 |

**Supplemental table 13:** Nodes input matrix for miRNA networks at T1 (left) and T2 (right). Column i) indicate the name of the stress-responsive miRNA, ii) group to which were assigned, and iii) number of stresses in which they are reactives. 10: represents miRNAs responsive to five or more stress conditions, 6: represents miRNAs responsive to three or four stress conditions, and 4: represents miRNAs responsive to one or two stress conditions.

**T1**

| miRNA | response range | node size |
|---|---|---|
| **miR166** | *Broad* | 10 |
| **miR156** | *Broad* | 10 |
| **miR319** | *Broad* | 10 |
| **miR157** | *Broad* | 10 |
| **miR396** | *Broad* | 10 |
| **miR398** | *Broad* | 10 |
| **miR167** | *Intermediate* | 6 |
| **miR6478** | *Intermediate* | 6 |
| **miR168** | *Intermediate* | 6 |
| **miR408** | *Intermediate* | 6 |
| **miR159** | *Narrow* | 4 |
| **miR162** | *Narrow* | 4 |
| **miR894** | *Narrow* | 4 |
| **miR160** | *Narrow* | 4 |
| **miR172** | *Narrow* | 4 |
| **miR395** | *Narrow* | 4 |
| **miR399** | *Narrow* | 4 |

**T2**

| miRNA | response range | node size |
|---|---|---|
| **miR396** | *Broad* | 10 |
| **miR398** | *Broad* | 10 |
| **miR408** | *Broad* | 10 |
| **miR6478** | *Broad* | 10 |
| **miR156** | *Intermediate* | 6 |
| **miR159** | *Intermediate* | 6 |
| **miR168** | *Intermediate* | 6 |
| **miR166** | *Intermediate* | 6 |
| **miR167** | *Intermediate* | 6 |
| **miR157** | *Narrow* | 4 |
| **miR319** | *Narrow* | 4 |
| **miR390** | *Narrow* | 4 |
| **miR399** | *Narrow* | 4 |
| **miR162** | *Narrow* | 4 |
| **miR169** | *Narrow* | 4 |
| **miR7130** | *Narrow* | 4 |
| **miR894** | *Narrow* | 4 |

**Supplemental table 14:** Edge input matrix. Values in source and target columns refer to position of the each stress responsive miRNAs in the nodes table (detailed below in miRNA codes). Edges values were established according common stress conditions between pairs (real values) and pondered in relation to the number of analyzed conditions (pondered values= real values/7).

## T1

| source | target | real value | pondered value |
|---|---|---|---|
| 0 | 1 | 6 | 0,86 |
| 0 | 2 | 6 | 0,86 |
| 0 | 3 | 5 | 0,71 |
| 0 | 4 | 5 | 0,71 |
| 0 | 5 | 5 | 0,71 |
| 0 | 6 | 4 | 0,57 |
| 0 | 7 | 4 | 0,57 |
| 0 | 8 | 3 | 0,43 |
| 0 | 9 | 3 | 0,43 |
| 0 | 10 | 2 | 0,29 |
| 0 | 11 | 2 | 0,29 |
| 0 | 12 | 2 | 0,29 |
| 0 | 13 | 1 | 0,14 |
| 0 | 14 | 1 | 0,14 |
| 0 | 15 | 1 | 0,14 |
| 0 | 16 | 1 | 0,14 |
| 1 | 2 | 5 | 0,71 |
| 1 | 3 | 5 | 0,71 |
| 1 | 4 | 5 | 0,71 |
| 1 | 5 | 4 | 0,57 |
| 1 | 6 | 4 | 0,57 |
| 1 | 7 | 3 | 0,43 |
| 1 | 8 | 3 | 0,43 |
| 1 | 9 | 3 | 0,43 |
| 1 | 10 | 2 | 0,29 |
| 1 | 11 | 2 | 0,29 |
| 1 | 12 | 2 | 0,29 |
| 1 | 13 | 1 | 0,14 |
| 1 | 14 | 1 | 0,14 |
| 1 | 15 | 1 | 0,14 |
| 1 | 16 | 1 | 0,14 |
| 2 | 3 | 4 | 0,57 |
| 2 | 4 | 5 | 0,71 |
| 2 | 5 | 5 | 0,71 |
| 2 | 6 | 4 | 0,57 |
| 2 | 7 | 4 | 0,57 |
| 2 | 8 | 3 | 0,43 |
| 2 | 9 | 3 | 0,43 |
| 2 | 10 | 2 | 0,29 |
| 2 | 11 | 2 | 0,29 |
| 2 | 12 | 1 | 0,14 |
| 2 | 13 | 1 | 0,14 |
| 2 | 14 | 1 | 0,14 |
| 2 | 15 | 0 | 0,00 |
| 2 | 16 | 1 | 0,14 |
| 3 | 4 | 4 | 0,57 |
| 3 | 5 | 3 | 0,43 |
| 3 | 6 | 4 | 0,57 |
| 3 | 7 | 3 | 0,43 |
| 3 | 8 | 2 | 0,29 |
| 3 | 9 | 2 | 0,29 |
| 3 | 10 | 1 | 0,14 |
| 3 | 11 | 2 | 0,29 |
| 3 | 12 | 2 | 0,29 |
| 3 | 13 | 1 | 0,14 |
| 3 | 14 | 0 | 0,00 |
| 3 | 15 | 1 | 0,14 |
| 3 | 16 | 0 | 0,00 |
| 4 | 5 | 4 | 0,57 |
| 4 | 6 | 4 | 0,57 |
| 4 | 7 | 3 | 0,43 |
| 4 | 8 | 3 | 0,43 |
| 4 | 9 | 3 | 0,43 |
| 4 | 10 | 2 | 0,29 |
| 4 | 11 | 2 | 0,29 |
| 4 | 12 | 1 | 0,14 |
| 4 | 13 | 1 | 0,14 |
| 4 | 14 | 1 | 0,14 |
| 4 | 15 | 0 | 0,00 |
| 4 | 16 | 1 | 0,14 |
| 5 | 6 | 3 | 0,43 |
| 5 | 7 | 4 | 0,57 |
| 5 | 8 | 3 | 0,43 |
| 5 | 9 | 3 | 0,43 |
| 5 | 10 | 2 | 0,29 |
| 5 | 11 | 1 | 0,14 |
| 5 | 12 | 1 | 0,14 |
| 5 | 13 | 1 | 0,14 |
| 5 | 14 | 1 | 0,14 |
| 5 | 15 | 0 | 0,00 |
| 5 | 16 | 1 | 0,14 |
| 6 | 7 | 3 | 0,43 |
| 6 | 8 | 2 | 0,29 |
| 6 | 9 | 2 | 0,29 |
| 6 | 10 | 1 | 0,14 |
| 6 | 11 | 2 | 0,29 |
| 6 | 12 | 1 | 0,14 |
| 6 | 13 | 1 | 0,14 |
| 6 | 14 | 0 | 0,00 |
| 6 | 15 | 0 | 0,00 |
| 6 | 16 | 0 | 0,00 |
| 7 | 8 | 2 | 0,29 |
| 7 | 9 | 2 | 0,29 |
| 7 | 10 | 1 | 0,14 |
| 7 | 11 | 1 | 0,14 |
| 7 | 12 | 1 | 0,14 |
| 7 | 13 | 1 | 0,14 |
| 7 | 14 | 0 | 0,00 |
| 7 | 15 | 0 | 0,00 |
| 7 | 16 | 0 | 0,00 |
| 8 | 9 | 2 | 0,29 |
| 8 | 10 | 2 | 0,29 |
| 8 | 11 | 1 | 0,14 |
| 8 | 12 | 1 | 0,14 |
| 8 | 13 | 1 | 0,14 |
| 8 | 14 | 1 | 0,14 |
| 8 | 15 | 0 | 0,00 |
| 8 | 16 | 1 | 0,14 |
| 9 | 10 | 1 | 0,14 |
| 9 | 11 | 0 | 0,00 |
| 9 | 12 | 0 | 0,00 |
| 9 | 13 | 1 | 0,14 |
| 9 | 14 | 1 | 0,14 |
| 9 | 15 | 0 | 0,00 |
| 9 | 16 | 1 | 0,14 |
| 10 | 11 | 1 | 0,14 |
| 10 | 12 | 1 | 0,14 |
| 10 | 13 | 0 | 0,00 |
| 10 | 14 | 1 | 0,14 |
| 10 | 15 | 0 | 0,00 |
| 10 | 16 | 1 | 0,14 |
| 11 | 12 | 1 | 0,14 |
| 11 | 13 | 0 | 0,00 |
| 11 | 14 | 0 | 0,00 |
| 11 | 15 | 0 | 0,00 |
| 11 | 16 | 0 | 0,00 |
| 12 | 13 | 0 | 0,00 |
| 12 | 14 | 0 | 0,00 |
| 12 | 15 | 1 | 0,14 |
| 12 | 16 | 0 | 0,00 |
| 13 | 14 | 0 | 0,00 |
| 13 | 15 | 0 | 0,00 |
| 13 | 16 | 0 | 0,00 |
| 14 | 15 | 0 | 0,00 |
| 14 | 16 | 1 | 0,14 |
| 15 | 16 | 0 | 0,00 |

## T2

| source | target | real value | pondered value |
|---|---|---|---|
| 0 | 1 | 4 | 0,57 |
| 0 | 2 | 4 | 0,57 |
| 0 | 3 | 5 | 0,71 |
| 0 | 4 | 4 | 0,57 |
| 0 | 5 | 4 | 0,57 |
| 0 | 6 | 4 | 0,57 |
| 0 | 7 | 3 | 0,43 |
| 0 | 8 | 3 | 0,43 |
| 0 | 9 | 2 | 0,29 |
| 0 | 10 | 2 | 0,29 |
| 0 | 11 | 2 | 0,29 |
| 0 | 12 | 2 | 0,29 |
| 0 | 13 | 1 | 0,14 |
| 0 | 14 | 1 | 0,14 |
| 0 | 15 | 1 | 0,14 |
| 0 | 16 | 1 | 0,14 |
| 1 | 2 | 5 | 0,71 |
| 1 | 3 | 3 | 0,43 |
| 1 | 4 | 4 | 0,57 |
| 1 | 5 | 2 | 0,29 |
| 1 | 6 | 4 | 0,57 |
| 1 | 7 | 2 | 0,29 |
| 1 | 8 | 3 | 0,43 |
| 1 | 9 | 2 | 0,29 |
| 1 | 10 | 2 | 0,29 |
| 1 | 11 | 2 | 0,29 |
| 1 | 12 | 1 | 0,14 |
| 1 | 13 | 1 | 0,14 |
| 1 | 14 | 0 | 0,00 |
| 1 | 15 | 0 | 0,00 |
| 1 | 16 | 1 | 0,14 |
| 2 | 3 | 3 | 0,43 |
| 2 | 4 | 4 | 0,57 |
| 2 | 5 | 2 | 0,29 |
| 2 | 6 | 4 | 0,57 |
| 2 | 7 | 2 | 0,29 |
| 2 | 8 | 3 | 0,43 |
| 2 | 9 | 2 | 0,29 |
| 2 | 10 | 2 | 0,29 |
| 2 | 11 | 2 | 0,29 |
| 2 | 12 | 1 | 0,14 |
| 2 | 13 | 1 | 0,14 |
| 2 | 14 | 0 | 0,00 |
| 2 | 15 | 0 | 0,00 |
| 2 | 16 | 1 | 0,14 |
| 3 | 4 | 3 | 0,43 |
| 3 | 5 | 4 | 0,57 |
| 3 | 6 | 3 | 0,43 |
| 3 | 7 | 3 | 0,43 |
| 3 | 8 | 3 | 0,43 |
| 3 | 9 | 2 | 0,29 |
| 3 | 10 | 2 | 0,29 |
| 3 | 11 | 2 | 0,29 |
| 3 | 12 | 2 | 0,29 |
| 3 | 13 | 1 | 0,14 |
| 3 | 14 | 1 | 0,14 |
| 3 | 15 | 1 | 0,14 |
| 3 | 16 | 1 | 0,14 |
| 4 | 5 | 2 | 0,29 |
| 4 | 6 | 4 | 0,57 |
| 4 | 7 | 2 | 0,29 |
| 4 | 8 | 3 | 0,43 |
| 4 | 9 | 2 | 0,29 |
| 4 | 10 | 2 | 0,29 |
| 4 | 11 | 2 | 0,29 |
| 4 | 12 | 1 | 0,14 |
| 4 | 13 | 1 | 0,14 |
| 4 | 14 | 0 | 0,00 |
| 4 | 15 | 0 | 0,00 |
| 4 | 16 | 1 | 0,14 |
| 5 | 6 | 2 | 0,29 |
| 5 | 7 | 2 | 0,29 |
| 5 | 8 | 2 | 0,29 |
| 5 | 9 | 1 | 0,14 |
| 5 | 10 | 1 | 0,14 |
| 5 | 11 | 1 | 0,14 |
| 5 | 12 | 2 | 0,29 |
| 5 | 13 | 0 | 0,00 |
| 5 | 14 | 1 | 0,14 |
| 5 | 15 | 1 | 0,14 |
| 5 | 16 | 1 | 0,14 |
| 6 | 7 | 2 | 0,29 |
| 6 | 8 | 3 | 0,43 |
| 6 | 9 | 2 | 0,29 |
| 6 | 10 | 2 | 0,29 |
| 6 | 11 | 2 | 0,29 |
| 6 | 12 | 1 | 0,14 |
| 6 | 13 | 1 | 0,14 |
| 6 | 14 | 0 | 0,00 |
| 6 | 15 | 0 | 0,00 |
| 6 | 16 | 1 | 0,14 |
| 7 | 8 | 2 | 0,29 |
| 7 | 9 | 2 | 0,29 |
| 7 | 10 | 1 | 0,14 |
| 7 | 11 | 2 | 0,29 |
| 7 | 12 | 1 | 0,14 |
| 7 | 13 | 1 | 0,14 |
| 7 | 14 | 0 | 0,00 |
| 7 | 15 | 0 | 0,00 |
| 7 | 16 | 0 | 0,00 |
| 8 | 9 | 2 | 0,29 |
| 8 | 10 | 2 | 0,29 |
| 8 | 11 | 2 | 0,29 |
| 8 | 12 | 1 | 0,14 |
| 8 | 13 | 1 | 0,14 |
| 8 | 14 | 0 | 0,00 |
| 8 | 15 | 0 | 0,00 |
| 8 | 16 | 1 | 0,14 |
| 9 | 10 | 1 | 0,14 |
| 9 | 11 | 2 | 0,29 |
| 9 | 12 | 0 | 0,00 |
| 9 | 13 | 1 | 0,14 |
| 9 | 14 | 0 | 0,00 |
| 9 | 15 | 0 | 0,00 |
| 9 | 16 | 0 | 0,00 |
| 10 | 11 | 1 | 0,14 |
| 10 | 12 | 1 | 0,14 |
| 10 | 13 | 1 | 0,14 |
| 10 | 14 | 0 | 0,00 |
| 10 | 15 | 0 | 0,00 |
| 10 | 16 | 1 | 0,14 |
| 11 | 12 | 0 | 0,00 |
| 11 | 13 | 1 | 0,14 |
| 11 | 14 | 0 | 0,00 |
| 11 | 15 | 0 | 0,00 |
| 11 | 16 | 0 | 0,00 |
| 12 | 13 | 0 | 0,00 |
| 12 | 14 | 0 | 0,00 |
| 12 | 15 | 0 | 0,00 |
| 12 | 16 | 1 | 0,14 |
| 13 | 14 | 0 | 0,00 |
| 13 | 15 | 0 | 0,00 |
| 13 | 16 | 0 | 0,00 |
| 14 | 15 | 1 | 0,14 |
| 14 | 16 | 0 | 0,00 |
| 15 | 16 | 0 | 0,00 |

**miRNAs code T1**

| | | | |
|---|---|---|---|
| miR166 | 0 | miR408 | 9 |
| miR156 | 1 | miR159 | 10 |
| miR319 | 2 | miR162 | 11 |
| miR157 | 3 | miR894 | 12 |
| miR396 | 4 | miR160 | 13 |
| miR398 | 5 | miR172 | 14 |
| miR167 | 6 | miR395 | 15 |
| miR6478 | 7 | miR399 | 16 |
| miR168 | 8 | | |

**miRNAs code T2**

| | | | |
|---|---|---|---|
| miR396 | 0 | miR157 | 9 |
| miR398 | 1 | miR319 | 10 |
| miR408 | 2 | miR390 | 11 |
| miR6478 | 3 | miR399 | 12 |
| miR156 | 4 | miR162 | 13 |
| miR159 | 5 | miR169 | 14 |
| miR168 | 6 | miR7130 | 15 |
| miR166 | 7 | miR894 | 16 |
| miR167 | 8 | | |

**Supplemental table 15:** Detailed information of the values used for calculate the correlation between relative accumulation of stress responsive miRNAs (estimated by sequencing) and their targets (estimated by qRT-PCR) in melon plants exposed to stress conditions in the both -T1 and T2- times analized here. The values are in LFC for miRNAs and Log2 of ΔΔCT for targets transcripts.

**T1**

| miRNA | Target | Stress | miRNA accumulation | target accumulation |
|---|---|---|---|---|
| *miR156* | SPL13 | COLD | 1,5 | 0,56 |
| | | SALINITY | -1,87 | -0,03 |
| | | MONOSP. | 1,61 | -0,17 |
| | | HSVd | -1,46 | 0,80 |
| | | AGRO | -1,18 | 0,77 |
| *miR166* | ATHB14 | COLD | -1,89 | 0,89 |
| | | DROUGHT | -1,04 | 1,67 |
| | | SALINITY | 1,94 | -1,09 |
| | | SHORTDAY | -1,05 | -0,06 |
| | | MONOSP. | 1,18 | -0,93 |
| | | HSVd | -1,47 | -0,29 |
| | | AGRO | -1,03 | -1,21 |
| *miR172* | AP2 | COLD | 2,02 | 0,29 |
| *miR319* | TCP | COLD | -1,21 | 0,61 |
| | | DROUGHT | -1,14 | -0,04 |
| | | SALINITY | 1,56 | -1,33 |
| | | SHORTDAY | 1,19 | -0,02 |
| | | HSVd | 1,62 | 0,51 |
| *miR408* | BBL | COLD | 1,41 | -2,32 |
| | | SALINITY | -1,3 | 0,27 |
| | | SHORTDAY | -2,62 | 0,57 |
| *miR398* | SOD | COLD | 2,07 | -1,23 |
| | | DROUGHT | -1,69 | 0,20 |
| | | SALINITY | -2,65 | 0,57 |
| | | SHORTDAY | -2,83 | 0,64 |
| | | HSVd | 1,52 | -1,25 |

**T2**

| miRNA | Target | Stress | miRNA accumulation | target accumulation |
|---|---|---|---|---|
| *miR408* | BBL | COLD | 6,92 | -3,99 |
| | | DROUGHT | 2,11 | -0,40 |
| | | MONOSP. | 4,22 | 0,43 |
| | | HSVd | 6,42 | 0,48 |
| | | AGRO | 5,7 | 1,15 |
| *miR169* | NFYA1 | SALINITY | 1,51 | 0,41 |
| *miR156* | SPL13 | COLD | 1,18 | -0,35 |
| | | MONOSP. | -1,12 | 0,19 |
| | | HSVd | -1,04 | 0,52 |
| | | AGRO | -1,54 | 0,71 |
| *miR398* | SOD | COLD | 9,27 | -1,82 |
| | | DROUGHT | 2,18 | -0,35 |
| | | MONOSP. | 5,78 | -0,38 |
| | | HSVd | 8,75 | -0,84 |
| | | AGRO | 7,5 | 0,16 |
| *miR396* | GRF | COLD | 2,06 | -0,68 |
| | | SALINITY | 1,9 | 0,43 |
| | | SHORTDAY | 1,93 | -0,26 |
| | | MONOSP. | 1,78 | 0,09 |
| | | HSVd | 1,28 | 0,36 |
| | | AGRO | -2,05 | 0,54 |

**Table S16:** Detailed information of the relative accumulation of miR398 and miR408 (estimated by sequencing) and their targets (estimated by qRT-PCR), in melon plants exposed to stress conditions analyzed here. The miRNA values are represented in LFC (determined by DESeq2). The transcripts values are represented as the Log2 of the ΔΔCT.

| Time | Treatment | SOD accumulation | miR398 accumulation |
|---|---|---|---|
| T1 | *COLD* | -1,226 | 1,23 |
| | *DROUGHT* | 0,196 | -1,10 |
| | *SALINITY* | 0,566 | -2,94 |
| | *SHORTDAY* | 0,641 | -3,79 |
| | *MONOSP.* | -1,682 | 0,48 |
| | *HSVd* | -1,246 | 1,76 |
| | *AGRO* | -1,324 | 1,06 |
| T2 | *COLD* | -1,817 | 9,63 |
| | *DROUGHT* | -0,350 | 3,22 |
| | *SALINITY* | 0,241 | -0,07 |
| | *SHORTDAY* | -0,741 | 0,00 |
| | *MONOSP.* | -0,375 | 6,76 |
| | *HSVd* | -0,843 | 9,26 |
| | *AGRO* | 0,161 | 8,41 |
| T4 | *COLD* | -0,906 | 7,72 |
| | *DROUGHT* | -0,975 | 9,56 |
| | *SALINITY* | -0,289 | -0,85 |
| | *SHORTDAY* | -0,576 | -1,06 |
| | *MONOSP.* | -0,347 | -0,58 |
| | *HSVd* | -2,237 | 10,45 |
| | *AGRO* | -1,515 | 9,34 |

***pearson=*** -0,55958461
***P=*** 8,4-3

| Time | Treatment | Cupredoxin accumulation | miR398 accumulation |
|---|---|---|---|
| T1 | *COLD* | -1,39 | 1,23 |
| | *DROUGHT* | 0,55 | -1,10 |
| | *SALINITY* | 0,71 | -2,94 |
| | *SHORTDAY* | 1,40 | -3,79 |
| | *MONOSP.* | -0,11 | 0,48 |
| | *HSVd* | -1,25 | 1,76 |
| | *AGRO* | 0,00 | 1,06 |
| T2 | *COLD* | -3,65 | 9,63 |
| | *DROUGHT* | -1,10 | 3,22 |
| | *SALINITY* | -0,70 | -0,07 |
| | *SHORTDAY* | -1,15 | 0,00 |
| | *MONOSP.* | -1,01 | 6,76 |
| | *HSVd* | -3,08 | 9,26 |
| | *AGRO* | -3,83 | 8,41 |
| T4 | *COLD* | 0,61 | 7,72 |
| | *DROUGHT* | -0,32 | 9,56 |
| | *SALINITY* | 0,90 | -0,85 |
| | *SHORTDAY* | 1,38 | -1,06 |
| | *MONOSP.* | 1,45 | -0,58 |
| | *HSVd* | -0,21 | 10,45 |
| | *AGRO* | 1,28 | 9,34 |

***pearson=*** -0,517
***P=*** 0,016

| Time | Treatment | BBL2 accumulation | miR408 accumulation |
|---|---|---|---|
| T1 | *COLD* | -4,36 | 1,53 |
| | *DROUGHT* | 0,28 | -0,54 |
| | *SALINITY* | 1,19 | -1,42 |
| | *SHORTDAY* | 0,21 | -2,86 |
| | *MONOSP.* | -0,89 | 0,31 |
| | *HSVd* | -2,35 | 0,41 |
| | *AGRO* | 0,00 | 0,42 |
| T2 | *COLD* | -4,31 | 7,16 |
| | *DROUGHT* | -0,37 | 2,92 |
| | *SALINITY* | 1,10 | 0,22 |
| | *SHORTDAY* | -0,69 | -0,95 |
| | *MONOSP.* | -0,46 | 4,82 |
| | *HSVd* | -1,94 | 6,71 |
| | *AGRO* | -1,53 | 6,23 |
| T4 | *COLD* | -2,46 | 1,59 |
| | *DROUGHT* | -0,95 | 6,39 |
| | *SALINITY* | 0,13 | 0,35 |
| | *SHORTDAY* | -0,32 | -2,14 |
| | *MONOSP.* | 0,28 | -1,98 |
| | *HSVd* | -3,77 | 5,95 |
| | *AGRO* | -1,96 | 7,32 |

***pearson=*** -0,58776878
***P=*** 5,1-3

**Table S17:** Detailed information of the accumulation values of representative stress-responsive and non stress-responsive melon miRNAs at T1 and T2, established by sequencing and additionally validated by stem loop qRT-PCR. The sequencing values are represented in LFC (determined by DESeq2). The qRT-PCR values are represented as the Log2 of the ΔΔCT. NS: non sequence data (sRNA reads below the limits established to be included in the differential analysis, see materials and methods).

**T1**

| miIRNAs | stress | Accumulation | |
|---|---|---|---|
| | | sequencing | qRT-PCR |
| *miR156* | COLD | 2,38 | 2,40 |
| | DROUGHT | -0,02 | 1,65 |
| | SALINITY | -0,09 | 2,49 |
| | SHORTDAY | 0,52 | 3,39 |
| | MONOSP. | 1,75 | 2,62 |
| | HSVd | -0,13 | 3,09 |
| | AGRO | 0,72 | 2,93 |
| *miR168* | COLD | 2,68 | 0,25 |
| | DROUGHT | -0,50 | -0,22 |
| | SALINITY | -0,85 | 0,78 |
| | SHORTDAY | -1,14 | 0,93 |
| | MONOSP. | -1,47 | 0,64 |
| | HSVd | -1,74 | 0,78 |
| | AGRO | -1,84 | 0,85 |
| *miR408* | COLD | 1,53 | 2,80 |
| | DROUGHT | -0,54 | 0,11 |
| | SALINITY | -1,42 | 0,66 |
| | SHORTDAY | -2,86 | -0,59 |
| | MONOSP. | 0,31 | 2,26 |
| | HSVd | 0,41 | 2,86 |
| | AGRO | 0,42 | 2,38 |
| *miR166* | COLD | 0,85 | 0,29 |
| | DROUGHT | -1,49 | -0,13 |
| | SALINITY | 0,46 | 2,11 |
| | SHORTDAY | -1,16 | 1,45 |
| | MONOSP. | -0,07 | 2,27 |
| | HSVd | -1,13 | 1,81 |
| | AGRO | -0,74 | 1,64 |
| *miR399* | COLD | -1,94 | -0,75 |
| | DROUGHT | -0,68 | -0,07 |
| | SALINITY | -0,74 | 0,35 |
| | SHORTDAY | -1,04 | 0,50 |
| | MONOSP. | 0,96 | 1,02 |
| | HSVd | -0,60 | 0,26 |
| | AGRO | -0,55 | 0,63 |
| *miR7130* | COLD | -2,98 | -0,05 |
| | DROUGHT | 0,19 | -0,08 |
| | SALINITY | -0,54 | 0,59 |
| | SHORTDAY | 0,18 | 0,74 |
| | MONOSP. | 0,26 | 0,91 |
| | HSVd | -1,25 | 0,68 |
| | AGRO | -0,21 | 1,04 |

**T2**

| miIRNAs | stress | Accumulation | |
|---|---|---|---|
| | | sequencing | qRT-PCR |
| *miR169* | COLD | NS | 0,13 |
| | DROUGHT | 1,5 | -0,16 |
| | SALINITY | 1,9 | -0,14 |
| | SHORTDAY | 1,0 | -0,18 |
| | MONOSP. | 1,0 | -0,08 |
| | HSVd | 0,8 | -3,19 |
| | AGRO | 1,5 | -1,65 |
| *miR396* | COLD | -0,6 | 1,43 |
| | DROUGHT | 0,8 | 0,55 |
| | SALINITY | 2,1 | 0,41 |
| | SHORTDAY | 2,1 | -0,29 |
| | MONOSP. | 1,9 | -0,01 |
| | HSVd | 1,3 | -3,61 |
| | AGRO | 0,9 | -2,62 |
| *miR398* | COLD | 9,6 | 5,06 |
| | DROUGHT | 3,2 | -0,82 |
| | SALINITY | -0,1 | -0,63 |
| | SHORTDAY | NS | -0,61 |
| | MONOSP. | 6,8 | 0,32 |
| | HSVd | 9,3 | 1,71 |
| | AGRO | 8,4 | 3,42 |
| *miR408* | COLD | 7,2 | 3,88 |
| | DROUGHT | 2,9 | -1,74 |
| | SALINITY | 0,2 | -0,74 |
| | SHORTDAY | -1,0 | -1,96 |
| | MONOSP. | 4,8 | 0,91 |
| | HSVd | 6,7 | -0,69 |
| | AGRO | 6,2 | 2,12 |
| *miR166* | COLD | 1,1 | -0,76 |
| | DROUGHT | -0,1 | -0,05 |
| | SALINITY | 0,7 | 0,74 |
| | SHORTDAY | 0,5 | -0,01 |
| | MONOSP. | 0,2 | 0,04 |
| | HSVd | 0,6 | -1,97 |
| | AGRO | 0,3 | -2,17 |
| *miR399* | COLD | 2,0 | 1,89 |
| | DROUGHT | -0,7 | -0,63 |
| | SALINITY | -0,4 | 0,32 |
| | SHORTDAY | -1,5 | -0,32 |
| | MONOSP. | -0,1 | -0,10 |
| | HSVd | -0,1 | -0,88 |
| | AGRO | -0,4 | -1,81 |
| *miR7130* | COLD | -2,1 | -1,11 |
| | DROUGHT | -0,7 | -1,53 |
| | SALINITY | -1,7 | -0,88 |
| | SHORTDAY | -0,6 | -0,56 |
| | MONOSP. | -0,5 | -0,38 |
| | HSVd | 0,2 | -3,23 |
| | AGRO | 0,4 | -1,59 |

**Table S18:** Matrix of presence/absence of stress-responsive miRNAs in soybean (GMA) and rice (OSA) plants. The values "1" and "0" represent respectively if whether or not a miRNA is reactive (with both either increased or decreased expression) to a specific stress condition. 1: reactive, 0: non reactive.

**GMA**

| *miRNA* | *Sress condition* | | | | | | | |
|---|---|---|---|---|---|---|---|---|
| | AL | ALK | D | DD | FLAG | P | SA | Total value |
| *mir169* | 1 | 1 | 1 | 0 | 0 | 0 | 0 | 3 |
| *mir390* | 1 | 1 | 0 | 1 | 0 | 1 | 1 | 5 |
| *mir156* | 1 | 1 | 1 | 1 | 0 | 1 | 1 | 6 |
| *mir159* | 1 | 1 | 0 | 0 | 0 | 1 | 0 | 3 |
| *mir4397* | 1 | 0 | 0 | 0 | 0 | 0 | 0 | 1 |
| *mir5671* | 1 | 0 | 0 | 0 | 0 | 0 | 1 | 2 |
| *mir5225* | 1 | 1 | 0 | 1 | 0 | 0 | 0 | 3 |
| *mir171* | 1 | 1 | 1 | 0 | 0 | 1 | 0 | 4 |
| *mir5037* | 1 | 0 | 0 | 0 | 0 | 0 | 0 | 1 |
| *mir172* | 1 | 1 | 0 | 0 | 1 | 0 | 0 | 3 |
| *mir160* | 1 | 1 | 0 | 0 | 0 | 1 | 0 | 3 |
| *mir1515* | 1 | 1 | 0 | 0 | 0 | 0 | 0 | 2 |
| *mir396* | 1 | 1 | 1 | 0 | 0 | 1 | 0 | 4 |
| *mir5373* | 1 | 1 | 1 | 0 | 0 | 0 | 0 | 3 |
| *mir482* | 1 | 1 | 1 | 1 | 0 | 1 | 0 | 5 |
| *mir167* | 1 | 1 | 1 | 1 | 0 | 0 | 1 | 5 |
| *mir862* | 1 | 0 | 0 | 0 | 0 | 0 | 0 | 1 |
| *mir164* | 1 | 1 | 1 | 0 | 0 | 1 | 0 | 4 |
| *mir1509* | 1 | 1 | 1 | 0 | 0 | 1 | 0 | 4 |
| *mir403* | 1 | 1 | 0 | 0 | 0 | 1 | 0 | 3 |
| *mir393* | 1 | 1 | 1 | 0 | 0 | 1 | 0 | 4 |
| *mir4415* | 1 | 0 | 0 | 1 | 0 | 0 | 0 | 2 |
| *mir2118* | 1 | 1 | 1 | 0 | 0 | 1 | 0 | 4 |
| *mir319* | 1 | 0 | 0 | 0 | 0 | 1 | 0 | 2 |
| *mir2606* | 0 | 1 | 0 | 0 | 0 | 0 | 0 | 1 |
| *mir1520* | 0 | 1 | 0 | 0 | 0 | 0 | 0 | 1 |
| *mir3522* | 0 | 1 | 0 | 1 | 0 | 0 | 1 | 3 |
| *mir5668* | 0 | 1 | 0 | 0 | 0 | 0 | 1 | 2 |
| *mir394* | 0 | 1 | 0 | 0 | 0 | 1 | 0 | 2 |
| *mir1510* | 0 | 1 | 0 | 0 | 0 | 1 | 0 | 2 |
| *mir408* | 0 | 1 | 1 | 1 | 1 | 0 | 1 | 5 |
| *mir5672* | 0 | 1 | 0 | 0 | 0 | 0 | 0 | 1 |
| *mir9728* | 0 | 1 | 0 | 0 | 0 | 0 | 0 | 1 |
| *mir166* | 0 | 1 | 1 | 1 | 0 | 1 | 1 | 5 |
| *mir5781* | 0 | 1 | 0 | 0 | 0 | 0 | 1 | 2 |
| *mir4416* | 0 | 1 | 0 | 0 | 0 | 0 | 0 | 1 |
| *mir1507* | 0 | 1 | 1 | 0 | 0 | 1 | 0 | 3 |
| *mir2109* | 0 | 1 | 0 | 0 | 0 | 1 | 0 | 2 |
| *mir2111* | 0 | 1 | 0 | 0 | 0 | 0 | 0 | 1 |
| *mir6300* | 0 | 1 | 0 | 0 | 0 | 1 | 0 | 2 |
| *mir1512* | 0 | 1 | 1 | 1 | 0 | 0 | 1 | 4 |
| *mir4345* | 0 | 1 | 0 | 0 | 0 | 0 | 0 | 1 |
| *mir4413* | 0 | 1 | 0 | 0 | 0 | 0 | 0 | 1 |
| *mir5674* | 0 | 1 | 0 | 0 | 0 | 0 | 0 | 1 |
| *mir1508* | 0 | 1 | 0 | 0 | 0 | 1 | 0 | 2 |
| *mir5782* | 0 | 1 | 0 | 0 | 0 | 0 | 0 | 1 |
| *mir5785* | 0 | 1 | 0 | 0 | 0 | 0 | 0 | 1 |
| *mir397* | 0 | 1 | 0 | 1 | 1 | 0 | 1 | 4 |
| *mir162* | 0 | 1 | 0 | 0 | 0 | 1 | 0 | 2 |
| *mir168* | 0 | 1 | 0 | 0 | 0 | 1 | 0 | 2 |
| *mir5376* | 0 | 1 | 0 | 0 | 0 | 1 | 0 | 2 |
| *mir1513* | 0 | 1 | 0 | 0 | 0 | 1 | 0 | 2 |
| *mir5769* | 0 | 1 | 0 | 0 | 0 | 0 | 0 | 1 |
| *mir5767* | 0 | 1 | 1 | 1 | 0 | 0 | 0 | 3 |
| *mir5786* | 0 | 1 | 0 | 1 | 0 | 0 | 0 | 2 |
| *mir4387* | 0 | 1 | 0 | 0 | 0 | 0 | 0 | 1 |
| *mir4412* | 0 | 1 | 0 | 0 | 0 | 0 | 0 | 1 |
| *mir5770* | 0 | 1 | 0 | 0 | 1 | 0 | 0 | 2 |
| *mir5372* | 0 | 1 | 0 | 0 | 0 | 1 | 0 | 2 |
| *mir5677* | 0 | 1 | 0 | 0 | 0 | 0 | 0 | 1 |
| *mir5761* | 0 | 1 | 0 | 1 | 0 | 0 | 0 | 2 |
| *mir5559* | 0 | 0 | 1 | 0 | 0 | 0 | 0 | 1 |
| *mir4996* | 0 | 0 | 1 | 0 | 0 | 0 | 0 | 1 |
| *mir9726* | 0 | 0 | 1 | 0 | 0 | 0 | 0 | 1 |
| *mir2119* | 0 | 0 | 1 | 1 | 0 | 0 | 1 | 3 |
| *mir5038* | 0 | 0 | 1 | 0 | 0 | 1 | 0 | 2 |
| *mir399* | 0 | 0 | 1 | 0 | 1 | 1 | 0 | 3 |
| *mir398* | 0 | 0 | 1 | 0 | 0 | 1 | 0 | 2 |
| *mir5670* | 0 | 0 | 0 | 0 | 0 | 1 | 0 | 1 |
| *mir1535* | 0 | 0 | 0 | 0 | 0 | 1 | 0 | 1 |
| *mir9749* | 0 | 0 | 0 | 0 | 0 | 1 | 0 | 1 |
| *mir5374* | 0 | 0 | 0 | 0 | 0 | 1 | 0 | 1 |

**OSA**

| miRNA | Stress conditions | | | | | | | | |
|---|---|---|---|---|---|---|---|---|---|
| | C | D | EL | H | RDV | RSV | SA | X | Total value |
| *mir390* | 1 | 1 | 0 | 1 | 0 | 1 | 1 | 0 | 5 |
| *mir2873* | 1 | 0 | 0 | 0 | 0 | 1 | 0 | 0 | 2 |
| *mir1875* | 1 | 0 | 0 | 0 | 0 | 0 | 0 | 0 | 1 |
| *mir812* | 1 | 1 | 0 | 1 | 1 | 0 | 1 | 1 | 6 |
| *mir1862* | 1 | 0 | 1 | 1 | 0 | 1 | 0 | 0 | 4 |
| *mir1863* | 1 | 1 | 0 | 1 | 0 | 1 | 1 | 1 | 6 |
| *mir167* | 1 | 1 | 1 | 1 | 0 | 1 | 1 | 0 | 6 |
| *mir1882* | 1 | 0 | 0 | 1 | 0 | 0 | 1 | 1 | 4 |
| *mir1423* | 1 | 1 | 0 | 1 | 0 | 0 | 1 | 0 | 4 |
| *mir5795* | 1 | 1 | 0 | 1 | 0 | 0 | 0 | 0 | 3 |
| *mir2867* | 1 | 0 | 0 | 1 | 0 | 1 | 1 | 0 | 4 |
| *mir3979* | 1 | 0 | 0 | 1 | 0 | 1 | 0 | 0 | 3 |
| *mir160* | 1 | 1 | 1 | 1 | 1 | 1 | 1 | 1 | 8 |
| *mir444* | 1 | 1 | 0 | 1 | 0 | 1 | 1 | 1 | 6 |
| *mir166* | 1 | 1 | 1 | 1 | 0 | 1 | 1 | 0 | 6 |
| *mir1425* | 1 | 0 | 0 | 1 | 0 | 1 | 1 | 0 | 4 |
| *mir164* | 1 | 1 | 0 | 1 | 1 | 1 | 1 | 0 | 6 |
| *mir5143* | 1 | 0 | 0 | 0 | 0 | 1 | 0 | 0 | 2 |
| *mir6251* | 1 | 0 | 0 | 1 | 0 | 0 | 0 | 0 | 2 |
| *mir2863* | 1 | 0 | 0 | 1 | 1 | 0 | 1 | 0 | 4 |
| *mir2121* | 0 | 1 | 0 | 1 | 0 | 0 | 1 | 0 | 3 |
| *mir1881* | 0 | 1 | 0 | 1 | 0 | 0 | 1 | 0 | 3 |
| *mir5150* | 0 | 1 | 1 | 1 | 0 | 0 | 1 | 1 | 5 |
| *mir5807* | 0 | 1 | 0 | 1 | 0 | 1 | 1 | 0 | 4 |
| *mir5802* | 0 | 1 | 0 | 1 | 1 | 0 | 1 | 0 | 4 |
| *mir1871* | 0 | 1 | 0 | 1 | 0 | 0 | 1 | 1 | 4 |
| *mir1861* | 0 | 1 | 0 | 1 | 1 | 0 | 1 | 0 | 4 |
| *mir396* | 0 | 1 | 1 | 1 | 1 | 1 | 1 | 1 | 7 |
| *mir535* | 0 | 1 | 0 | 1 | 0 | 1 | 0 | 0 | 3 |
| *mir1879* | 0 | 1 | 0 | 1 | 0 | 0 | 1 | 0 | 3 |
| *mir1856* | 0 | 1 | 0 | 1 | 1 | 0 | 1 | 0 | 4 |
| *mir169* | 0 | 1 | 1 | 1 | 1 | 1 | 1 | 1 | 7 |
| *mir1870* | 0 | 1 | 0 | 1 | 0 | 1 | 1 | 0 | 4 |
| *mir827* | 0 | 1 | 0 | 1 | 1 | 0 | 0 | 1 | 4 |
| *mir394* | 0 | 1 | 1 | 1 | 0 | 0 | 1 | 1 | 5 |
| *mir2880* | 0 | 0 | 1 | 0 | 0 | 0 | 0 | 0 | 1 |
| *mir1432* | 0 | 0 | 1 | 1 | 0 | 1 | 0 | 0 | 3 |
| *mir528* | 0 | 0 | 1 | 1 | 0 | 0 | 0 | 0 | 2 |
| *mir2118* | 0 | 0 | 1 | 0 | 0 | 0 | 0 | 0 | 1 |
| *mir408* | 0 | 0 | 1 | 1 | 0 | 0 | 1 | 0 | 3 |
| *mir393* | 0 | 0 | 1 | 1 | 0 | 1 | 0 | 0 | 3 |
| *mir168* | 0 | 0 | 1 | 1 | 0 | 1 | 0 | 0 | 3 |
| *mir810* | 0 | 0 | 1 | 0 | 0 | 0 | 0 | 0 | 1 |
| *mir156* | 0 | 0 | 1 | 1 | 0 | 1 | 0 | 0 | 3 |
| *mir398* | 0 | 0 | 1 | 0 | 0 | 0 | 0 | 0 | 1 |
| *mir2877* | 0 | 0 | 1 | 0 | 0 | 0 | 0 | 0 | 1 |
| *mir5813* | 0 | 0 | 0 | 1 | 1 | 0 | 0 | 0 | 2 |
| *mir5814* | 0 | 0 | 0 | 1 | 0 | 0 | 1 | 0 | 2 |
| *mir5801* | 0 | 0 | 0 | 1 | 0 | 1 | 0 | 0 | 2 |
| *mir319* | 0 | 0 | 0 | 1 | 0 | 1 | 0 | 0 | 2 |
| *mir529* | 0 | 0 | 0 | 1 | 0 | 1 | 0 | 0 | 2 |
| *mir172* | 0 | 0 | 0 | 1 | 0 | 1 | 0 | 0 | 2 |
| *mir1876* | 0 | 0 | 0 | 1 | 0 | 1 | 1 | 0 | 3 |
| *mir1860* | 0 | 0 | 0 | 1 | 0 | 0 | 1 | 0 | 2 |
| *mir531* | 0 | 0 | 0 | 1 | 0 | 0 | 0 | 0 | 1 |
| *mir5508* | 0 | 0 | 0 | 1 | 1 | 0 | 0 | 0 | 2 |
| *mir1873* | 0 | 0 | 0 | 1 | 0 | 0 | 1 | 0 | 2 |
| *mir1868* | 0 | 0 | 0 | 1 | 0 | 0 | 1 | 0 | 2 |
| *mir397* | 0 | 0 | 0 | 1 | 0 | 0 | 0 | 0 | 1 |
| *mir162* | 0 | 0 | 0 | 1 | 0 | 0 | 0 | 0 | 1 |
| *mir820* | 0 | 0 | 0 | 1 | 0 | 1 | 0 | 1 | 3 |
| *mir5794* | 0 | 0 | 0 | 1 | 0 | 0 | 0 | 0 | 1 |
| *mir1427* | 0 | 0 | 0 | 1 | 0 | 0 | 0 | 0 | 1 |
| *mir6254* | 0 | 0 | 0 | 1 | 0 | 0 | 1 | 1 | 3 |
| *mir1850* | 0 | 0 | 0 | 1 | 1 | 1 | 0 | 0 | 3 |
| *mir1320* | 0 | 0 | 0 | 1 | 0 | 0 | 0 | 0 | 1 |
| *mir1430* | 0 | 0 | 0 | 1 | 0 | 0 | 0 | 0 | 1 |
| *mir439* | 0 | 0 | 0 | 1 | 0 | 0 | 0 | 0 | 1 |
| *mir435* | 0 | 0 | 0 | 1 | 0 | 0 | 0 | 0 | 1 |
| *mir5144* | 0 | 0 | 0 | 1 | 0 | 0 | 0 | 0 | 1 |
| *mir159* | 0 | 0 | 0 | 1 | 1 | 0 | 1 | 1 | 4 |
| *mir1864* | 0 | 0 | 0 | 1 | 0 | 0 | 1 | 0 | 2 |
| *mir1859* | 0 | 0 | 0 | 1 | 1 | 0 | 0 | 0 | 2 |
| *mir1878* | 0 | 0 | 0 | 0 | 1 | 0 | 0 | 0 | 1 |
| *mir5811* | 0 | 0 | 0 | 0 | 1 | 1 | 0 | 0 | 2 |
| *mir2871* | 0 | 0 | 0 | 0 | 1 | 1 | 0 | 0 | 2 |
| *mir5512* | 0 | 0 | 0 | 0 | 1 | 0 | 0 | 0 | 1 |
| *mir5788* | 0 | 0 | 0 | 0 | 1 | 0 | 0 | 0 | 1 |
| *mir171* | 0 | 0 | 0 | 0 | 0 | 1 | 0 | 0 | 1 |
| *mir1865* | 0 | 0 | 0 | 0 | 0 | 1 | 0 | 0 | 1 |
| *mir1883* | 0 | 0 | 0 | 0 | 0 | 0 | 1 | 0 | 1 |
| *mir5083* | 0 | 0 | 0 | 0 | 0 | 0 | 1 | 0 | 1 |
| *mir530* | 0 | 0 | 0 | 0 | 0 | 0 | 0 | 1 | 1 |
| *mir2878* | 0 | 0 | 0 | 0 | 0 | 0 | 0 | 1 | 1 |
| *mir399* | 0 | 0 | 0 | 0 | 0 | 0 | 0 | 1 | 1 |

**Table S19:** Nodes input matrix for miRNA networks in soybean (GMA - left) and rice (OSA - right). Column i) indicate the name of the stress-responsive miRNA, ii) group to which were assigned, and iii) number of stresses in which they are reactives. 10: represents miRNAs responsive to five or more stress conditions, 6: represents miRNAs responsive to three or four stress conditions, and 4: represents miRNAs responsive to one or two stress conditions.

**GMA**

| miRNA | Response range | node size |
|---|---|---|
| ***mir156*** | *Broad* | 10 |
| ***mir390*** | *Broad* | 10 |
| ***mir482*** | *Broad* | 10 |
| ***mir167*** | *Broad* | 10 |
| ***mir408*** | *Broad* | 10 |
| ***mir166*** | *Broad* | 10 |
| ***mir171*** | Intermediate | 6 |
| ***mir396*** | Intermediate | 6 |
| ***mir164*** | Intermediate | 6 |
| ***mir1509*** | Intermediate | 6 |
| ***mir393*** | Intermediate | 6 |
| ***mir2118*** | Intermediate | 6 |
| ***mir1512*** | Intermediate | 6 |
| ***mir397*** | Intermediate | 6 |
| ***mir169*** | Intermediate | 6 |
| ***mir159*** | Intermediate | 6 |
| ***mir5225*** | Intermediate | 6 |
| ***mir172*** | Intermediate | 6 |
| ***mir160*** | Intermediate | 6 |
| ***mir5373*** | Intermediate | 6 |
| ***mir403*** | Intermediate | 6 |
| ***mir3522*** | Intermediate | 6 |
| ***mir1507*** | Intermediate | 6 |
| ***mir5767*** | Intermediate | 6 |
| ***mir2119*** | Intermediate | 6 |
| ***mir399*** | Intermediate | 6 |
| ***mir5671*** | Narrow | 4 |
| ***mir1515*** | Narrow | 4 |
| ***mir4415*** | Narrow | 4 |
| ***mir319*** | Narrow | 4 |
| ***mir5668*** | Narrow | 4 |
| ***mir394*** | Narrow | 4 |
| ***mir1510*** | Narrow | 4 |
| ***mir5781*** | Narrow | 4 |
| ***mir2109*** | Narrow | 4 |
| ***mir6300*** | Narrow | 4 |
| ***mir1508*** | Narrow | 4 |
| ***mir162*** | Narrow | 4 |
| ***mir168*** | Narrow | 4 |
| ***mir5376*** | Narrow | 4 |
| ***mir1513*** | Narrow | 4 |
| ***mir5786*** | Narrow | 4 |
| ***mir5770*** | Narrow | 4 |
| ***mir5372*** | Narrow | 4 |
| ***mir5761*** | Narrow | 4 |
| ***mir5038*** | Narrow | 4 |
| ***mir398*** | Narrow | 4 |
| ***mir4397*** | Narrow | 4 |
| ***mir5037*** | Narrow | 4 |
| ***mir862*** | Narrow | 4 |
| ***mir2606*** | Narrow | 4 |
| ***mir1520*** | Narrow | 4 |
| ***mir5672*** | Narrow | 4 |
| ***mir9728*** | Narrow | 4 |
| ***mir4416*** | Narrow | 4 |
| ***mir2111*** | Narrow | 4 |
| ***mir4345*** | Narrow | 4 |
| ***mir4413*** | Narrow | 4 |
| ***mir5674*** | Narrow | 4 |
| ***mir5782*** | Narrow | 4 |
| ***mir5785*** | Narrow | 4 |
| ***mir5769*** | Narrow | 4 |
| ***mir4387*** | Narrow | 4 |
| ***mir4412*** | Narrow | 4 |
| ***mir5677*** | Narrow | 4 |
| ***mir5559*** | Narrow | 4 |
| ***mir4996*** | Narrow | 4 |
| ***mir9726*** | Narrow | 4 |
| ***mir5670*** | Narrow | 4 |
| ***mir1535*** | Narrow | 4 |
| ***mir9749*** | Narrow | 4 |
| ***mir5374*** | Narrow | 4 |

**OSA**

| miRNA | Response range | Node size |
|---|---|---|
| ***mir160*** | Broad | 10 |
| ***mir396*** | Broad | 10 |
| ***mir169*** | Broad | 10 |
| ***mir812*** | Broad | 10 |
| ***mir1863*** | Broad | 10 |
| ***mir167*** | Broad | 10 |
| ***mir444*** | Broad | 10 |
| ***mir166*** | Broad | 10 |
| ***mir164*** | Broad | 10 |
| ***mir390*** | Broad | 10 |
| ***mir5150*** | Broad | 10 |
| ***mir394*** | Broad | 10 |
| ***mir1862*** | Intermediate | 6 |
| ***mir1882*** | Intermediate | 6 |
| ***mir1423*** | Intermediate | 6 |
| ***mir2867*** | Intermediate | 6 |
| ***mir1425*** | Intermediate | 6 |
| ***mir2863*** | Intermediate | 6 |
| ***mir5807*** | Intermediate | 6 |
| ***mir5802*** | Intermediate | 6 |
| ***mir1871*** | Intermediate | 6 |
| ***mir1861*** | Intermediate | 6 |
| ***mir1856*** | Intermediate | 6 |
| ***mir1870*** | Intermediate | 6 |
| ***mir827*** | Intermediate | 6 |
| ***mir159*** | Intermediate | 6 |
| ***mir5795*** | Intermediate | 6 |
| ***mir3979*** | Intermediate | 6 |
| ***mir2121*** | Intermediate | 6 |
| ***mir1881*** | Intermediate | 6 |
| ***mir535*** | Intermediate | 6 |
| ***mir1879*** | Intermediate | 6 |
| ***mir1432*** | Intermediate | 6 |
| ***mir408*** | Intermediate | 6 |
| ***mir393*** | Intermediate | 6 |
| ***mir168*** | Intermediate | 6 |
| ***mir156*** | Intermediate | 6 |
| ***mir1876*** | Intermediate | 6 |
| ***mir820*** | Intermediate | 6 |
| ***mir6254*** | Intermediate | 6 |
| ***mir1850*** | Intermediate | 6 |
| ***mir2873*** | Narrow | 4 |
| ***mir5143*** | Narrow | 4 |
| ***mir6251*** | Narrow | 4 |
| ***mir528*** | Narrow | 4 |
| ***mir5813*** | Narrow | 4 |
| ***mir5814*** | Narrow | 4 |
| ***mir5801*** | Narrow | 4 |
| ***mir319*** | Narrow | 4 |
| ***mir529*** | Narrow | 4 |
| ***mir172*** | Narrow | 4 |
| ***mir1860*** | Narrow | 4 |
| ***mir5508*** | Narrow | 4 |
| ***mir1873*** | Narrow | 4 |
| ***mir1868*** | Narrow | 4 |
| ***mir1864*** | Narrow | 4 |
| ***mir1859*** | Narrow | 4 |
| ***mir5811*** | Narrow | 4 |
| ***mir2871*** | Narrow | 4 |
| ***mir1875*** | Narrow | 4 |
| ***mir2880*** | Narrow | 4 |
| ***mir2118*** | Narrow | 4 |
| ***mir810*** | Narrow | 4 |
| ***mir398*** | Narrow | 4 |
| ***mir2877*** | Narrow | 4 |
| ***mir531*** | Narrow | 4 |
| ***mir397*** | Narrow | 4 |
| ***mir162*** | Narrow | 4 |
| ***mir5794*** | Narrow | 4 |
| ***mir1427*** | Narrow | 4 |
| ***mir1320*** | Narrow | 4 |
| ***mir1430*** | Narrow | 4 |
| ***mir439*** | Narrow | 4 |
| ***mir435*** | Narrow | 4 |
| ***mir5144*** | Narrow | 4 |
| ***mir1878*** | Narrow | 4 |
| ***mir5512*** | Narrow | 4 |
| ***mir5788*** | Narrow | 4 |
| ***mir171*** | Narrow | 4 |
| ***mir1865*** | Narrow | 4 |
| ***mir1883*** | Narrow | 4 |
| ***mir5083*** | Narrow | 4 |
| ***mir530*** | Narrow | 4 |
| ***mir2878*** | Narrow | 4 |
| ***mir399*** | Narrow | 4 |
